# The functional divergence of plant ESCRT components TOL3, SNF7.1, and VPS4 during salt stress response

**DOI:** 10.1101/2025.09.06.674610

**Authors:** Madeleine Schnurer, Alois Schweighofer, Julia Hilscher, Eszter Kapusi, Barbara Korbei, Thomas Nägele, Lilian Kang, Paul Fuhrmann, Susanne Danninger, Leonie Scharl, Asem Bock, Giada Marino, Dario Leister, Verena Ibl

## Abstract

Salinity, a major abiotic stressor, impacts protein trafficking and alters transport routes. The endosomal sorting complex required for transport (ESCRT)is a key player for membrane protein sorting and is increasingly recognized as pivotal element in abiotic stress responses. However, the functional role of ESCRT proteins in cereal crops under salt stress remains unexplored. Using the as near as possible to nature DRD-BIBLOX (Dark-Root device Brick Black Box), barley (*Hordeum vulgare*, L.), although relatively salt-tolerant, exhibits impaired germination, seedling growth, and an altered root system architecture (RSA) under high salinity. Shotgun proteomic analysis revealed spatio-temporal regulation of ESCRT proteins in shoots and roots during seedling development. We identified HvSNF7.1 and HvVPS4 as critical, but differently acting modulators of salt stress responses in barley tissues. In this study we further reveal novel functions of ESCRT-0-like TOM-1-like proteins (TOLs): Previously described as gatekeepers of degradative protein sorting and as regulators of abscisic acid signaling, growth, and heat stress responses in Arabidopsis (*Arabidopsis thaliana*), HvTOLs had not been characterized for their role in salinity adaptation during plant development. Functional analyses demonstrated lethality in Hv*toldouble* and Hv*tolquadruple* mutants under standard growth conditions. In Arabidopsis, loss of At*TOL3* alone impaired germination and root growth under salt stress, underscoring the essential role of individual TOL proteins in the spatio-temporal regulation of abiotic stress adaptation. These findings highlight ESCRT-mediated trafficking as a key determinant of cereal resilience under saline conditions and provide a molecular framework for improving crop stress tolerance.

## INTRODUCTION

Endomembrane trafficking is a crucial cellular process conserved across eukaryotes, responsible for transporting materials between distinct membrane-bound organelles and carrying out essential housekeeping roles that are necessary for cellular function and development (Surpin and Raikhel, 2004). In plant cells, the primary endomembrane trafficking pathway—along secretory and endocytic routes—plays an essential role in stress responses, by ensuring targeted transport of stress-related cargo (Wang et al., 2020). Salinity, a significant abiotic stressor, drastically impacts the growth, development, and productivity of many plants throughout various stages of their life cycle (Khan et al., 2022). Specifically, high salinity interferes with regulation of protein trafficking and causes significant changes in the plant endomembrane system, including modifications to transport pathways and structural remodeling of organelles like the endoplasmic reticulum (ER) and the vacuole (Sampaio et al., 2022; Dermendjiev et al., 2021).

The endosomal sorting complex required for transport (ESCRT) serves an essential purpose as a cargo recognition and membrane deforming machinery within the plant endomembrane system. In plants the ESCRT complex consists of five subcomplexes, ESCRT-0-like, ESCRT-I, ESCRT-II, ESCRT-III, and associated proteins like VPS4. (Otegui and Spitzer, 2008; Henne et al., 2011) Besides participating in processes such as autophagy, cytokinesis, viral replication, abscisic acid (ABA) signaling, chloroplast turnover, membrane repair, multivesicular body (MVB) biogenesis, and membrane protein sorting (Otegui and Spitzer, 2008; Henne et al., 2011), further, autophagic degradation of plastid proteins in Arabidopsis (Spitzer et al., 2015), and protein sorting into protein bodies during barley endosperm development (Roustan et al., 2020), the ESCRT machinery plays a crucial role in plant stress responses and cellular processes and is assumed to be strongly affected in response to abiotic stress (Gao et al., 2017; Isono, 2021). Several ESCRT subcomplexes have been studied specifically with respect to stress conditions. So far, ESCRT-0-like constituents, Tom-1-like proteins (target of Myb-1-like proteins, TOLs) have been characterized under normal condition, where TOLs are involved in degradative sorting to the vacuole and are particularly important for auxin-mediated responses and PIN protein sorting (Korbei et al., 2013; Sauer and Friml, 2014). Arabidopsis TOLs are further involved in the modulation of ABA signaling and drought stress responses (Moulinier-Anzola et al., 2024). Additionally, TOL mutants or knock-outs led to a number of different phenotypes in Arabidopsis under normal growth conditions (Korbei et al., 2013; Sauer and Friml, 2014).

In Arabidopsis, the ESCRT-I component VPS23A is involved in regulating ABA signaling upon salt stress and it also enhances salt tolerance by strengthening the salt overly sensitive (SOS) signaling module (Lou et al., 2020; Liu et al., 2025). Furthermore, in Arabidopsis seedlings, the AAA-ATPase, VPS4 is up-regulated under salt stress (Gong et al., 2001). Additionally, salt stress sensitivity was observed in transgenic Arabidopsis with lower VPS4 levels (Ho et al., 2010). Multi-omics, microscopy, and physiology studies showed that in salt stressed *Nicotiana tabacum* (tobacco), the ESCRT-III associated protein VPS46.2 was significantly up-regulated and SNF7.1 levels very drastically down-regulated (Garcia de la Garma et al., 2015). However, the function of TOL proteins in abiotic stress remains largely unexplored.

Our recent proteomics studies on barley grains revealed HvESCRT proteins at high levels and differently abundant during various stages of the grain development (Roustan et al., 2020). However, our current understanding lacks the knowledge of how HvESCRTs are involved in salt stress response during barley germination and seedling growth. In this work, we used our DRD-BIBLOX to study salt stress application in barley as near as possible to nature - to promote knowledge transfer from the laboratory to the field (Dermendjiev et al., 2023). Over-expression of Hv*SNF7*.*1* and Hv*VPS4* show distinctly different response to salt stress. Additionally, different combinations of Hv*toldouble* and Hv*tolquadruple* mutants were lethal in barley. The knock-out of At*TOL3* shows salt-induced altered germination and root growth response. Collectively, these results reveal critical roles for single ESCRT proteins in salt stress response and underline the necessity of plant-, tissue-, and time-specific analysis of key-players like ESCRT proteins.

## RESULTS

### Barley seedling growth is sensitive to salt

We have previously identified the effects of high salt concentration on the actin cytoskeleton and vacuolization in aleurone cells during early germination of barley (Dermendjiev et al., 2021). Therefore we further wanted to investigate the impact of salt stress on barley seedling growth. For this purpose, barley grains were germinated and grown under conditions as near as possible to nature, in rhizoboxes, placed in DRD-BIBLOXes under control (TAP) and salt stress (EC30) conditions for 8 and 16 days after sowing (DAS) **(Figure1 A-F)**. Significantly shorter root and shoot length of both stages were observed upon EC30 compared to TAP **(Figure1 G-H, Figure S1 A-D)**. Plant root lengths of 8 and 16 DAS EC30 plants reduced to 22 % and 68 % of TAP plants. Shoot lengths of 8 and 16 DAS EC30 plants reduced to 2 % and 48 % of TAP plants respectively. Additionally, seminal roots and therefore overall root numbers reduced under EC30 **(Figure S1 E)**. Subsequently, wet and dry weight of root and shoot material showed a significant decrease of biomass under EC30. Root mass of 8 and 16 DAS plants decreased to 10 % and 44 % of TAP, and shoot mass decreased to nearly 0 % and 44 % of control, upon EC30 respectively **(Figure S1 F,G)**. Summarizing, barley seedlings subjected to EC30 revealed substantial reductions in root and shoot lengths, and biomass.

**Figure 1.**
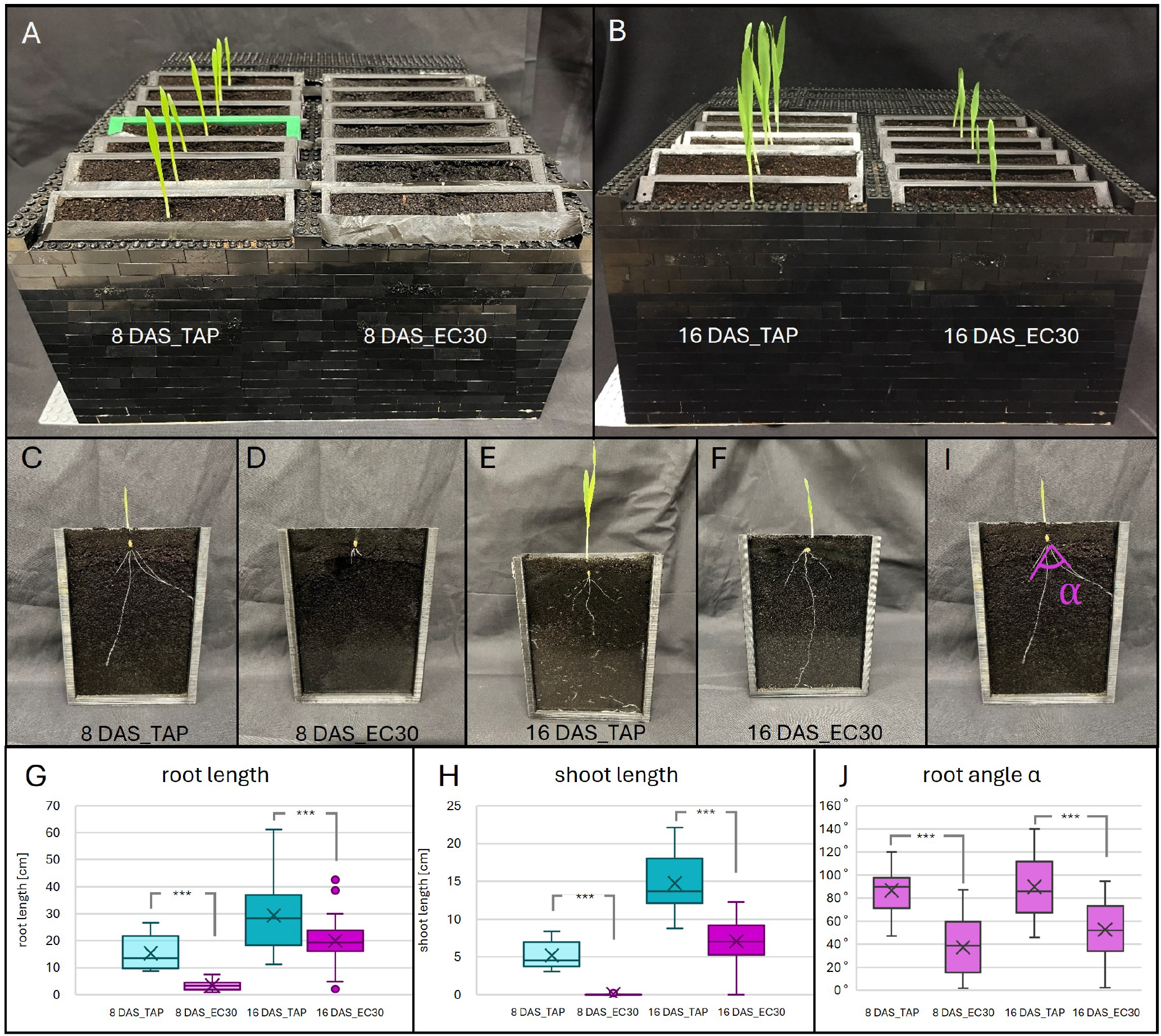
Salt stress affects barley germination and seedling growth. (A) DRD-BIBLOX with rhizoboxes and barley plants 8 days after sowing (DAS). Boxes on the left side were watered with tap water (TAP) (control; 8 DAS-TAP), on the right with salt stress (EC30) (8 DAS-EC30). (B) DRD-BIBLOX with rhizoboxes and barley plants 16 DAS. Boxes on the left show 16 DAS-TAP, on the right 16 DAS-EC30 plants. (C-F) Rhizoboxes with barley plants of 8 or 16 DAS, grown under EC30 or TAP conditions. (G) Plant root length of 8 and 16 DAS under TAP condition, and EC30 conditions. (H) Barley shoot lengths of 8 and 16 DAS at TAP conditions, and EC30 conditions. The bars indicate the medians and the x indicate the means; n=27-51 plants per category; (I) Indicates angle *α*, spanning along the two furthest apart seminal roots. (J) Graphs of root angle *α* of 8 and 16 DAS plants, grown under TAP and EC30 conditions. n=14-28 plants; *** indicating a p-value of ≤ 0.001 upon *t*-tests.

We next analysed the impact of EC30 on barley germination using the DRD-BIBLOX with an infrared camera set-up (Dermendjiev et al., 2023). This device allowed barley root phenotyping and tracking from the point of sowing to the late seedling stage in soil, in dark. The resulting picture time series shows the first appearance of root structures for TAP plants at 2.8 DAS, and for EC30 plants at 6.9 DAS **(Figure S2)**. Besides germination delay, slower root growth and reduced seminal root production were tracked under EC30.

We then analysed the root architecture in more detail. Several studies already showed some effects of salt stress on the root system architecture (RSA) (Farooq et al., 2024; Shelden and Munns, 2023). Using the DRD-BIBLOX-camera system, we tracked a reduction of seminal root numbers under EC30. Furthermore, root angles of primary and seminal roots (*α, β, γ, δ, ϵ*) were measured as depicted in **Figure1 I,J and Figure S3 A-E**. Angles *α* (seminal root growth angle; root spread angle), *β*, and *γ* significantly decreased under EC30, meanwhile the angles *δ* and *ϵ* increased. Additionally, we found roots of EC30 plants to appear thicker **(Figure S3 F)**, since more tiny root hairs would grow along the primary and seminal roots. Fewer root hairs were found on the TAP plants, thus the root hair length were exceeding those of salt stressed plants by far. Generally TAP plants showed a more wide spread and branched root system. Altogether, these results show that salt stress alters barley germination, and seedling root- and shoot development. Especially, the RSA is highly affected.

### Detecting proteome-wide effects in barley root and shoot during development and salt stress

Previous proteomic analyses have identified proteins involved in salt stress response, including ion transporters and channels, and stress-responsive proteins, which are crucial for maintaining ionic balance and facilitating detoxification under saline conditions (van Zelm et al., 2020; Mostek et al., 2015). Further, proteomic studies during barley germination specifically revealed the significant effect of high salt concentration on proteins of the actin cytoskeleton and proteins involved in vacuolization processes in aleurone cells (Dermendjiev et al., 2021). Additionally, ESCRT(- related) proteins have been described to be responsive to salt stress (Liu et al., 2025; Xia et al., 2016). In this study, we applied untargeted shotgun proteomic analysis to investigate first, proteome-wide effects in barley root and shoot during development and salt stress, and second, if ESCRT proteins are affected by salt stress. Protein extracts of TAP and EC30 roots (R) and shoots (S), of 8 DAS and 16 DAS plants, were subjected to liquid chromatography, tandem mass spectrometry analysis, followed by data analysis. **(Figure S4)**.

The principal component analysis (PCA) plot of the whole dataset shows a clear separation of the different sample groups, and clustering of biological replicates **(Figure S5A)**. For the root and shoot material a total of 2141 proteins were identified and 1381 of them were significantly differently regulated upon development or salt stress **(Supplementary file Perseus)**. Z-scores were calculated and hierarchical cluster analysis (HCA) over the whole dataset visually confirmed uniformity of biological replicates, as well as they highlighted differences between sample groups **(Figure S5B,C)**.

According to *t*-tests for 8 DAS roots of TAP versus EC30 (8RTAP_vs_ 8REC30) a total of 224 proteins, for 16RTAP_vs_16REC30 736 proteins, for 16 DAS shoots of TAP versus EC30 (16STAP_vs_16SEC30) 52 proteins, for 8RTAP_vs_16RTAP 26 proteins, and for 8STAP_vs_ 16STAP 662 proteins could be identified as being tissue-specifically and significantly up- or down-regulated upon development or salt stress. Those identified proteins were analysed and categorized according to their gene ontology, including cellular components, molecular functions, and the molecular processes they are associated with **(Figure S5 D-F, Figure S6,7,8, Supplement Data)**. Subsequently, to understand the systemics of development and salt stress on barley on the proteome level, z-scores were calculated, and HCA of above discussed sample groupings allowed a more detailed overview of protein levels during development and under salt stress **(Figure S9 A-E)**.

The HCA for root development shows three clusters, where most of the involved proteins contribute to cluster two and three, showing their up-regulation during development **(Figure S9 A)**. Hereto belong proteins involved in development and growth, as well as cell wall and cell wall biogenesis associated proteins. Regarding shoot development HCA shows 4 clusters **(Figure S9 B)**. In cluster one and two protein levels are up-regulated during development. They comprise proteins of photosynthesis, photo-respiration, and proton transport, as well as proteins involved in glycolytic processes, and defense response. In cluster three protein levels of translation associated proteins, protein folding, ubiquitin associated proteins, as well as vesicle associated proteins are down-regulated during shoot development. While cluster four shows again an up-regulation of levels of photosynthesis related proteins.

Focusing on developing roots, z-scores of label free quantification (LFQ) intensity values over several protein groups were calculated **(Table S1, Figure S10**,**11)**. This revealed, COPI vesicle associated proteins, actin related proteins, v-ATPase related proteins, proteolysis associated proteins, ubiquitin associated proteins, LEA proteins, root development related proteins, auxin catabolism related, and kinase associated proteins were up-regulated during development. Meanwhile annexins, lypoxygenases, and glycolysis associated proteins were down-regulated during early root development.

Comparing 8 DAS TAP to EC30 roots, three protein clusters were found **(Figure S9 C)**. In cluster one and two levels of defense related proteins, stress response proteins, and ABA associated proteins, including the ABA receptor HvPYL2, are up-regulated under salt stress. While in cluster three, protein levels of cytoskeleton and vesicle associated proteins are down-regulated under salt stress. Further comparing 16 DAS TAP to EC30 roots, three clusters formed **(Figure S9 D)**. Cluster one and three show a down-regulation of again cytoskeleton, proteolysis, vesicle, and protein transport associated proteins. And cluster two shows defense response and stress response proteins up-regulated during salt stress. Further comparing 16 DAS TAP to EC30 shoots **(Figure S9 E)**, on the one hand defense response proteins are up-regulated in cluster one, and on the other hand proteolysis and photosynthesis related proteins are down-regulated as part of cluster two, when facing salt stress.

Focusing on how salt affects root proteomes, z-scores of label free quantification (LFQ) intensity values over several protein groups were calculated **(Table1, Figure S10,11)**. This revealed under salt stress, protein abundances of ABA-associated proteins, auxin catabolism proteins, dynamin associated proteins, COPI vesicle associated proteins, actin related proteins, v-ATPase related proteins, annexins, cell wall biogenesis associated proteins, vesicle associated proteins, proteolysis associated proteins, ubiquitin associated proteins, glycolysis associated proteins, photosynthesis associated proteins, root development related proteins, auxin catabolism proteins, and kinase associated proteins were less abundant. Further, defense response associated proteins and LEA proteins were more abundant in roots upon salt stress. Furthermore, identified lipoxygenases in roots decreased in their abundance during 8 DAS EC30, but increased during 16 DAS EC30 compared to TAP.

**Table 1:**
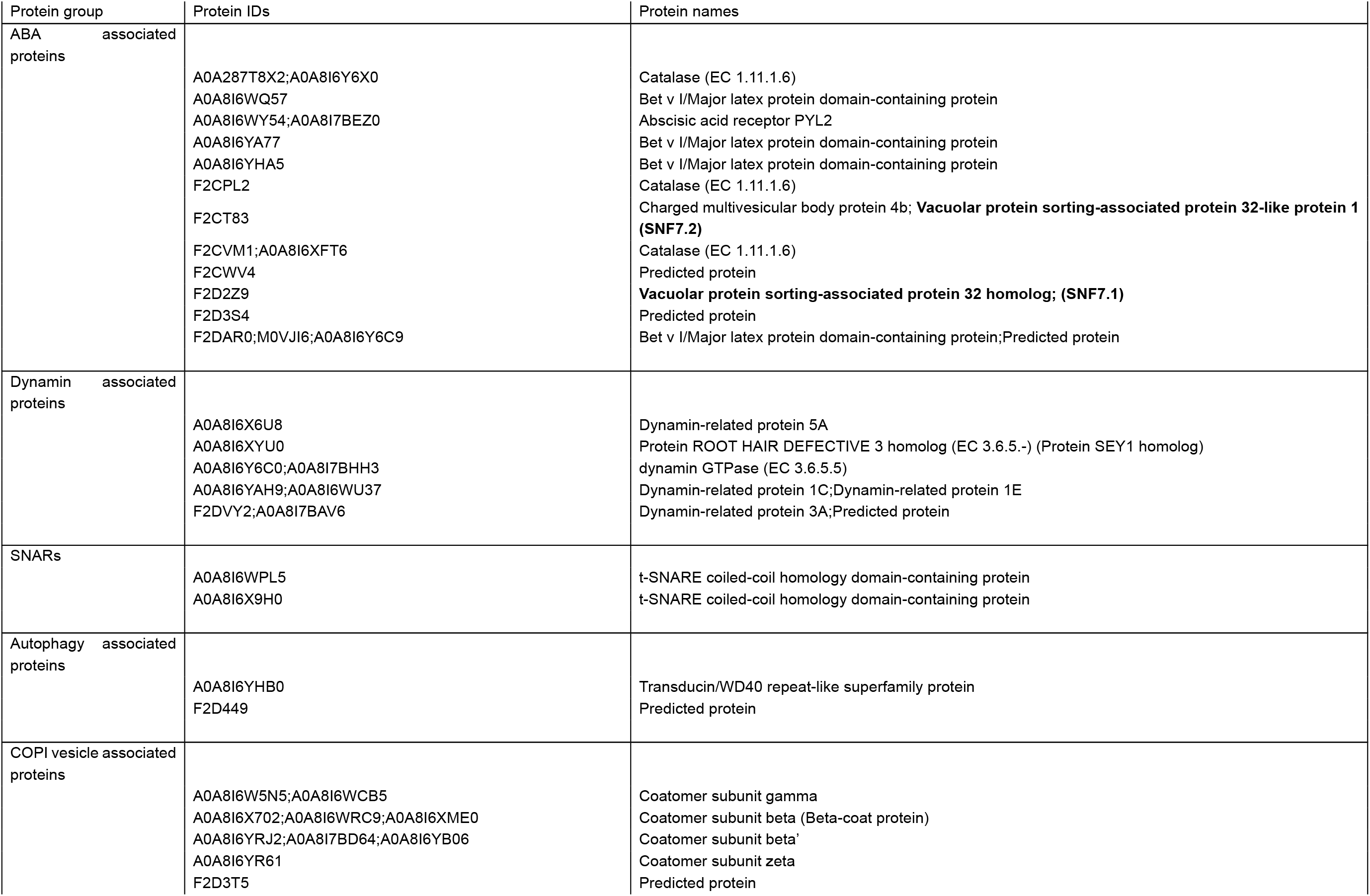
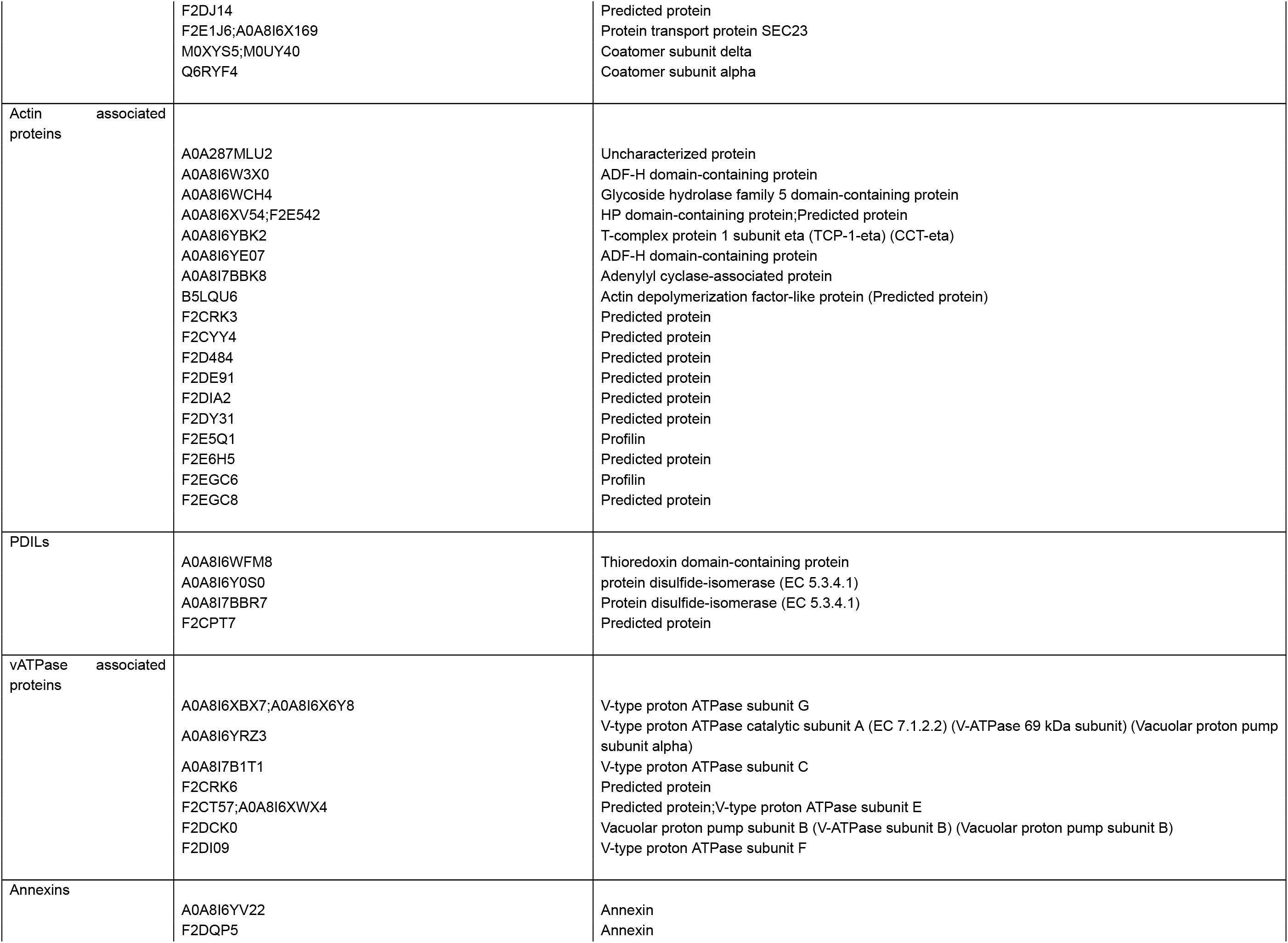
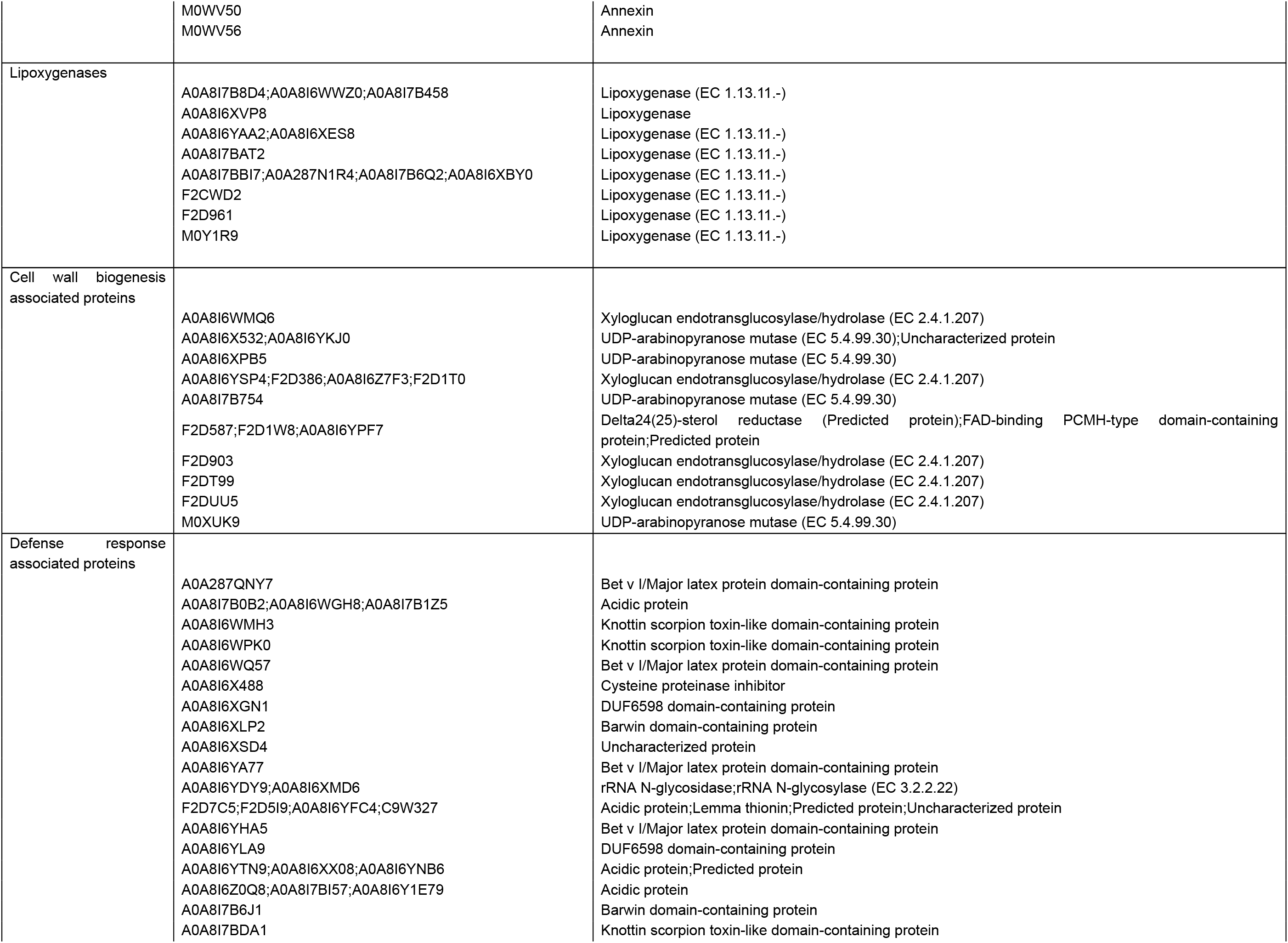
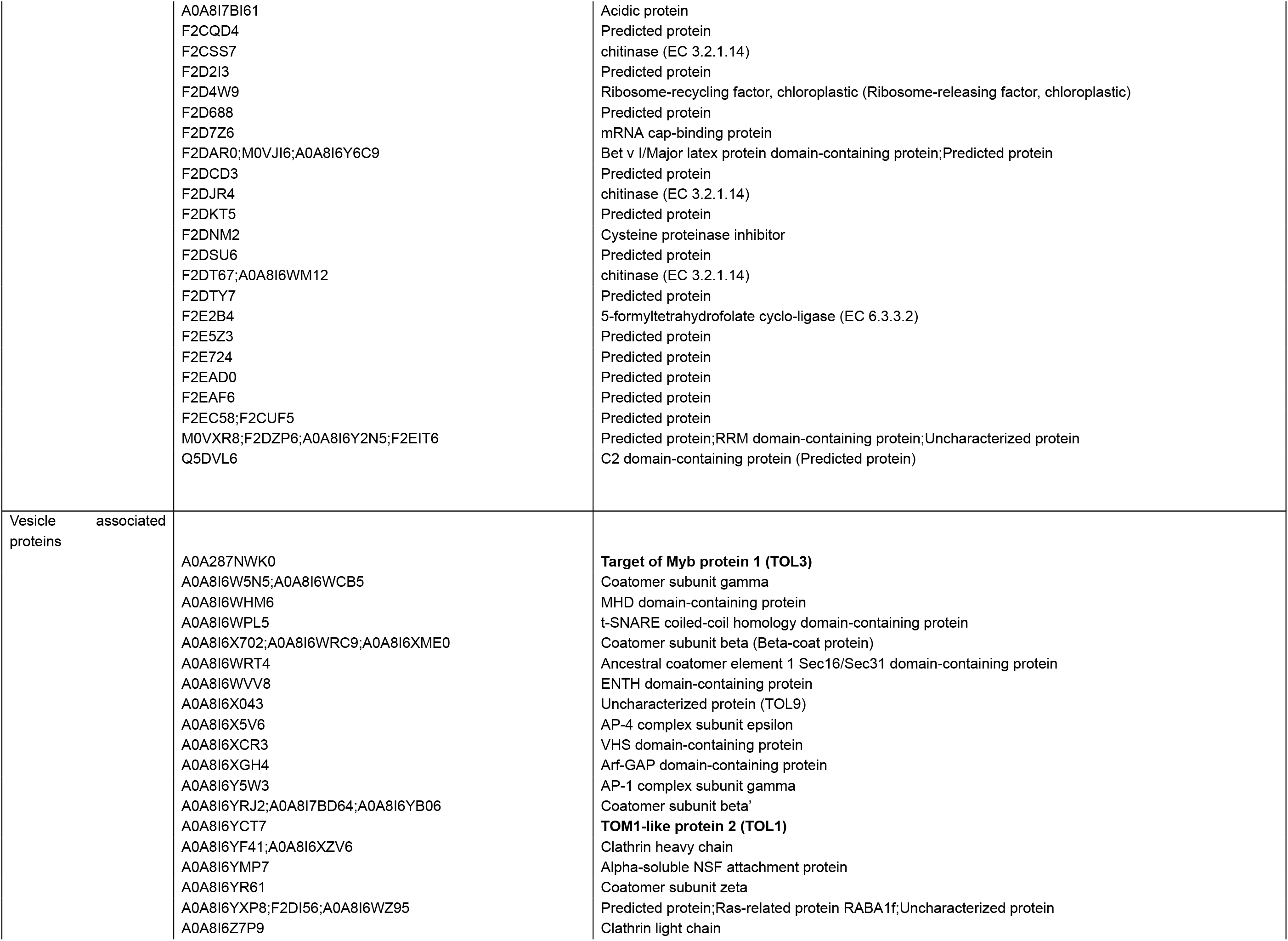
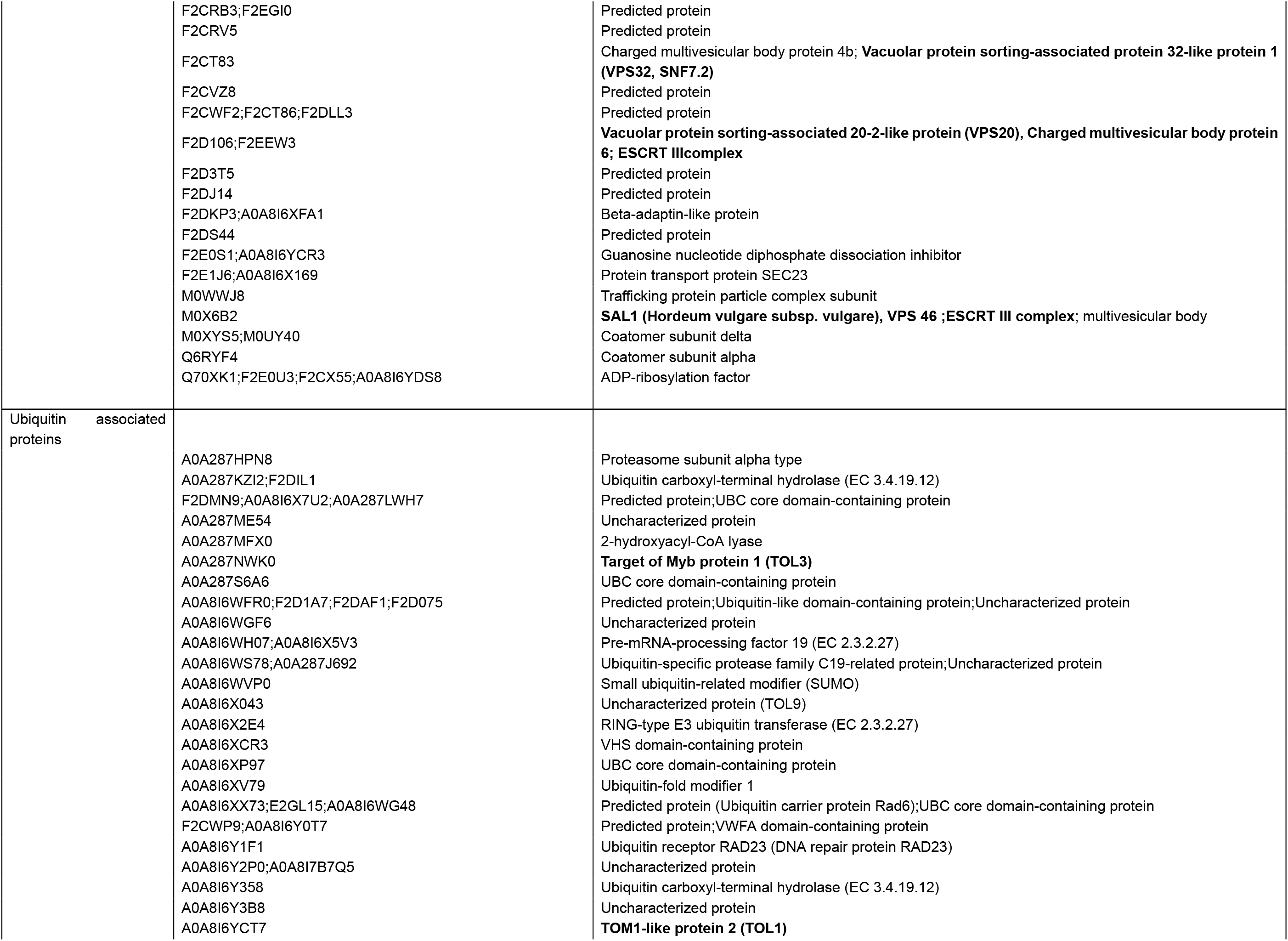
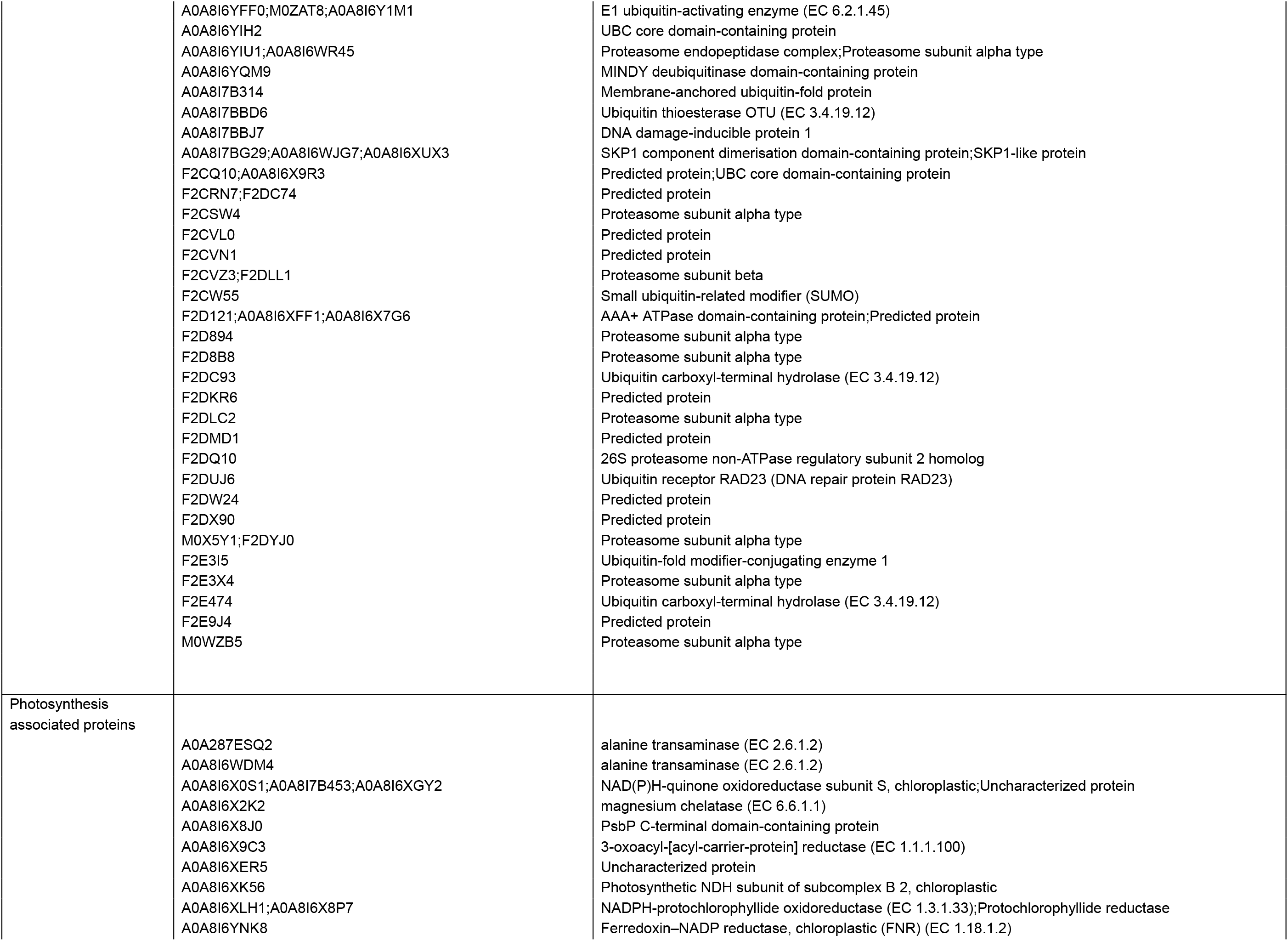
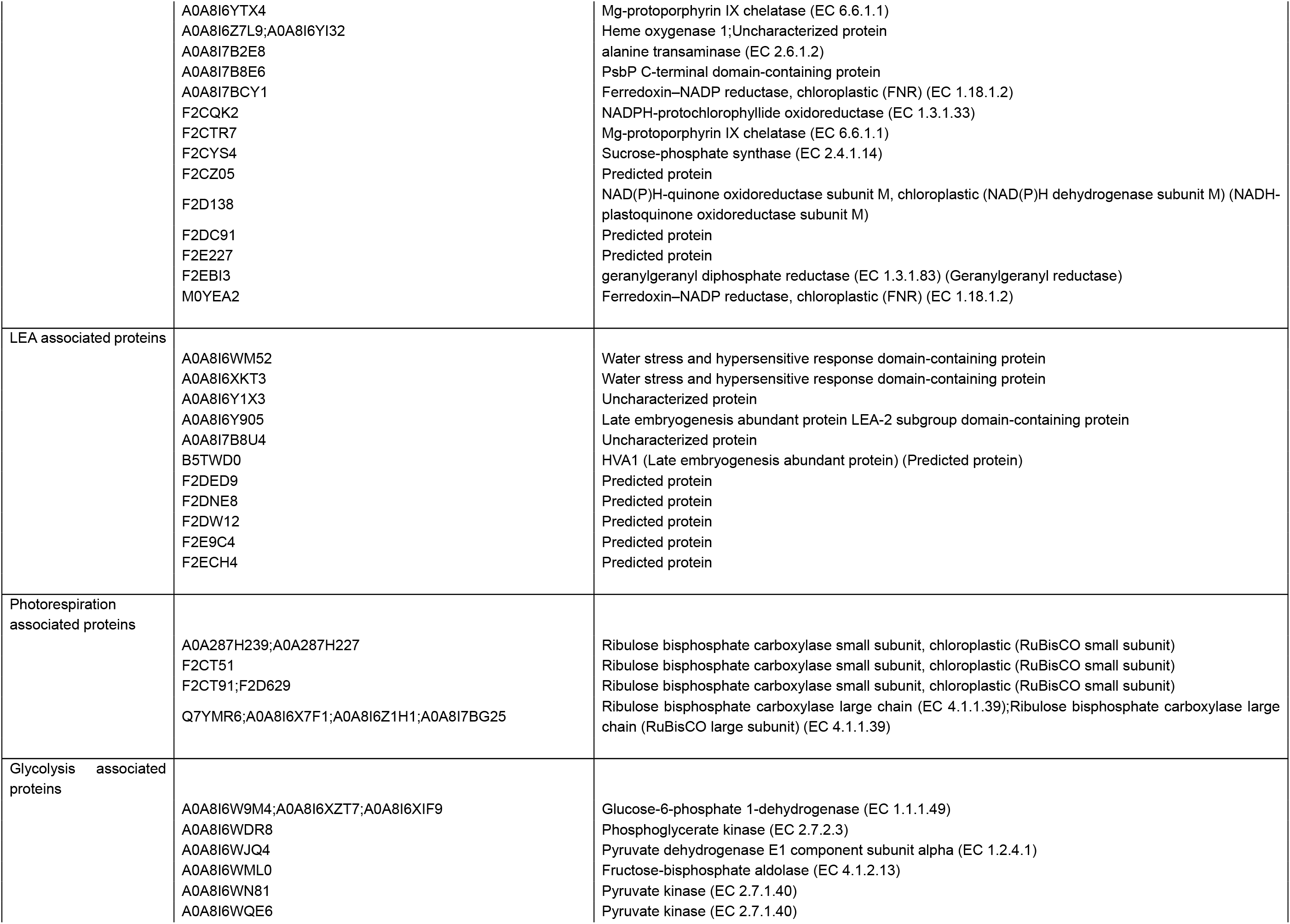
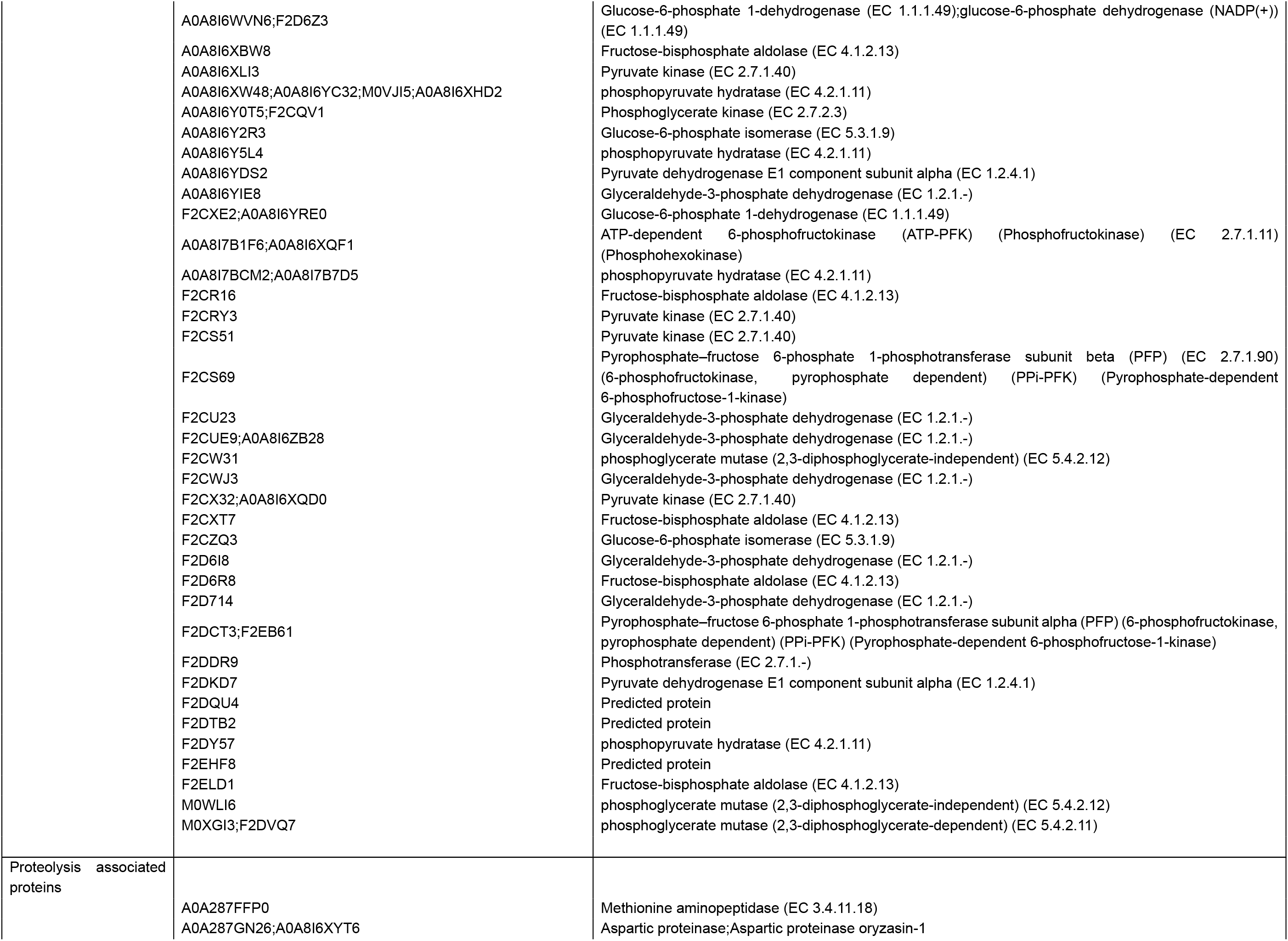
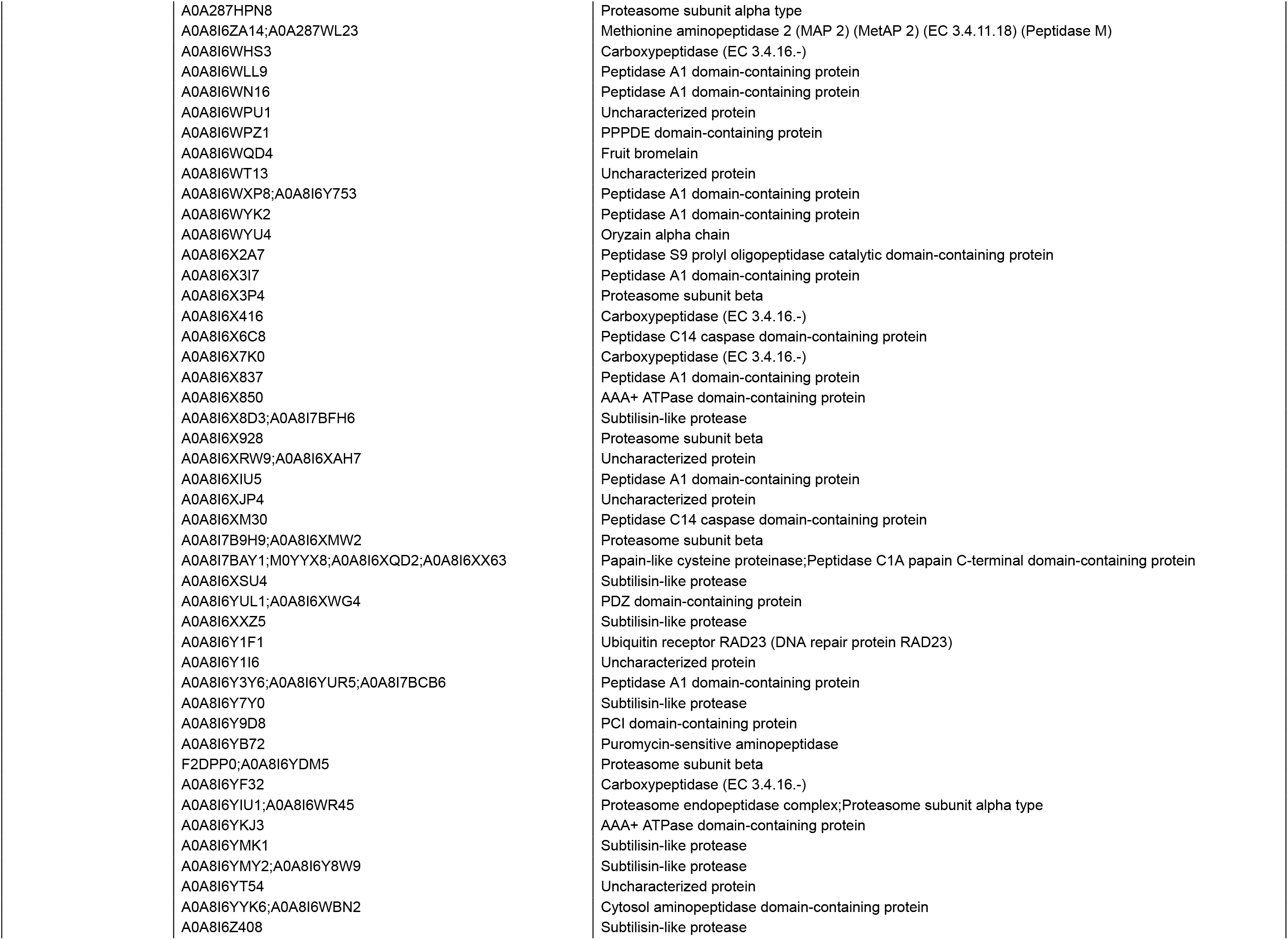
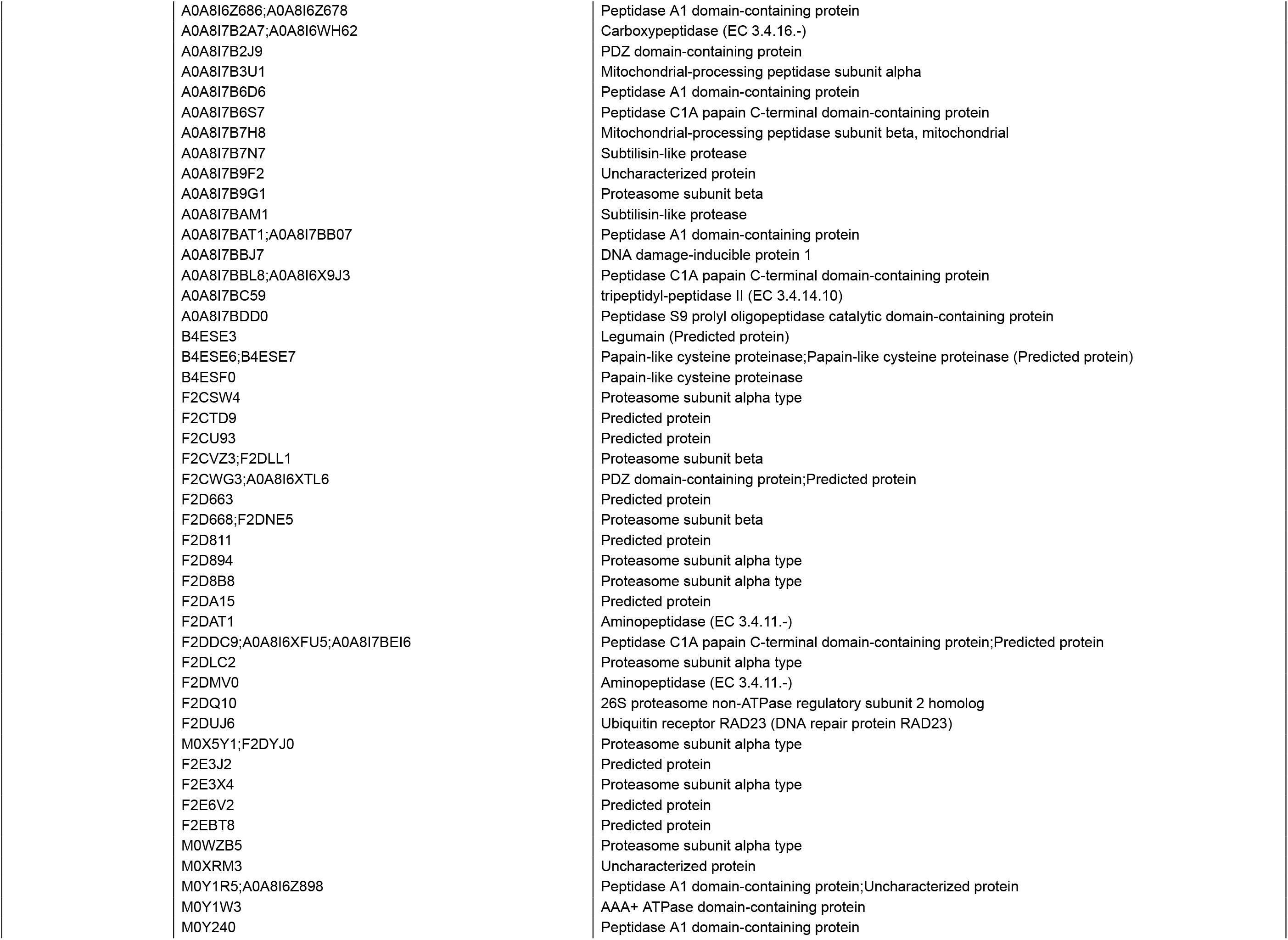
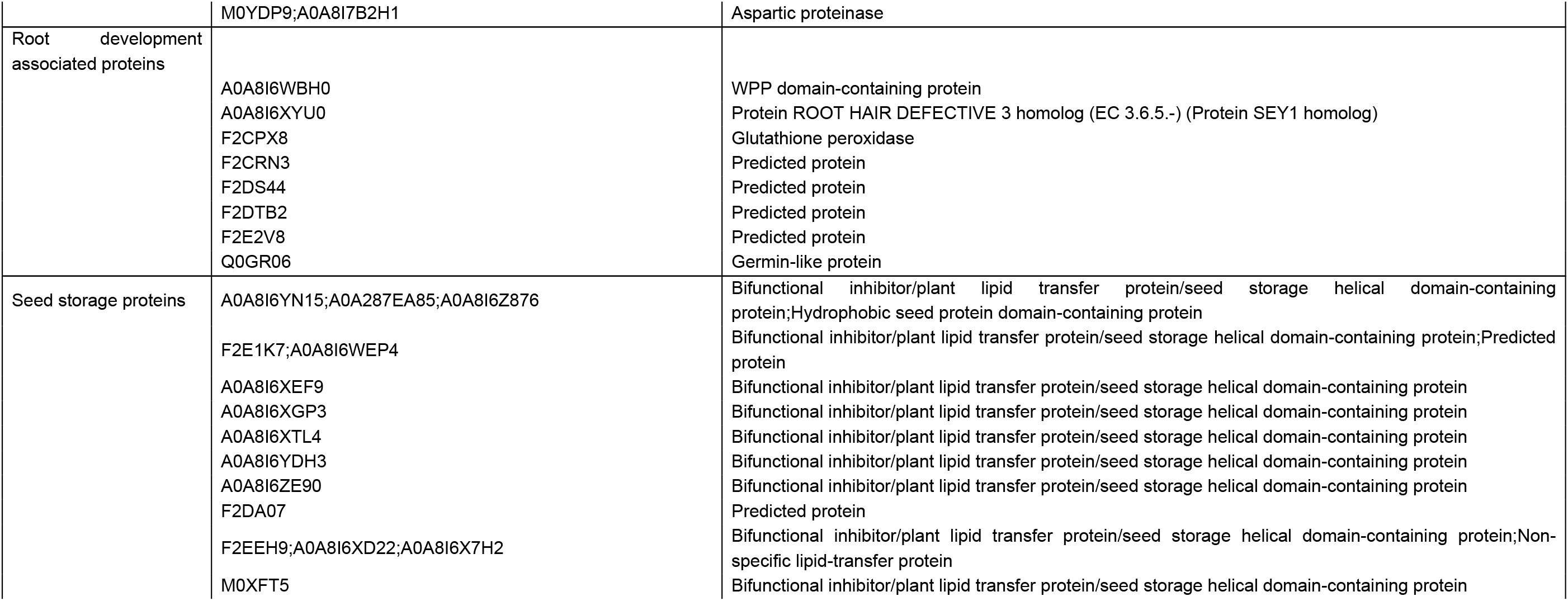
Protein groups identified among salt stress proteomic data

**Table 2.**
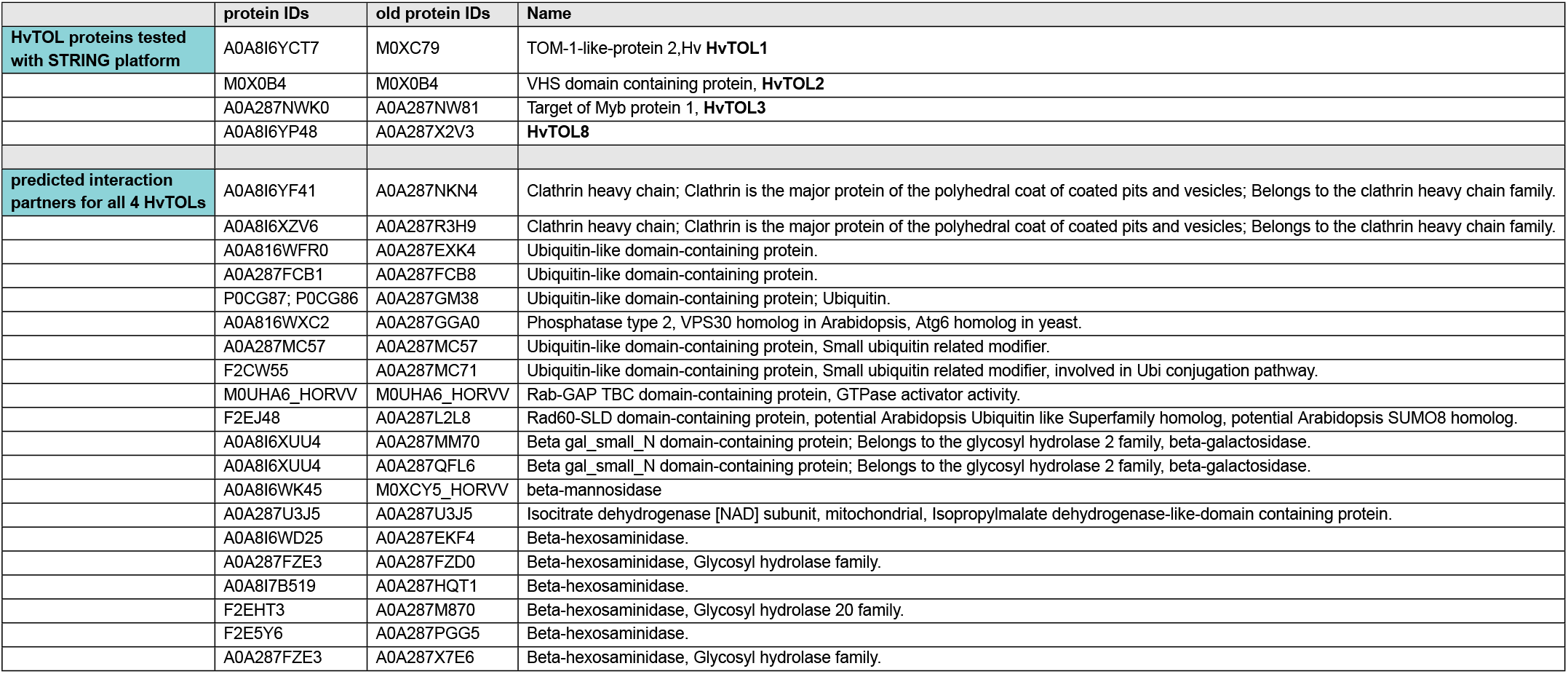
String data HvTOLs and interacting proteins.

**Table 3.**
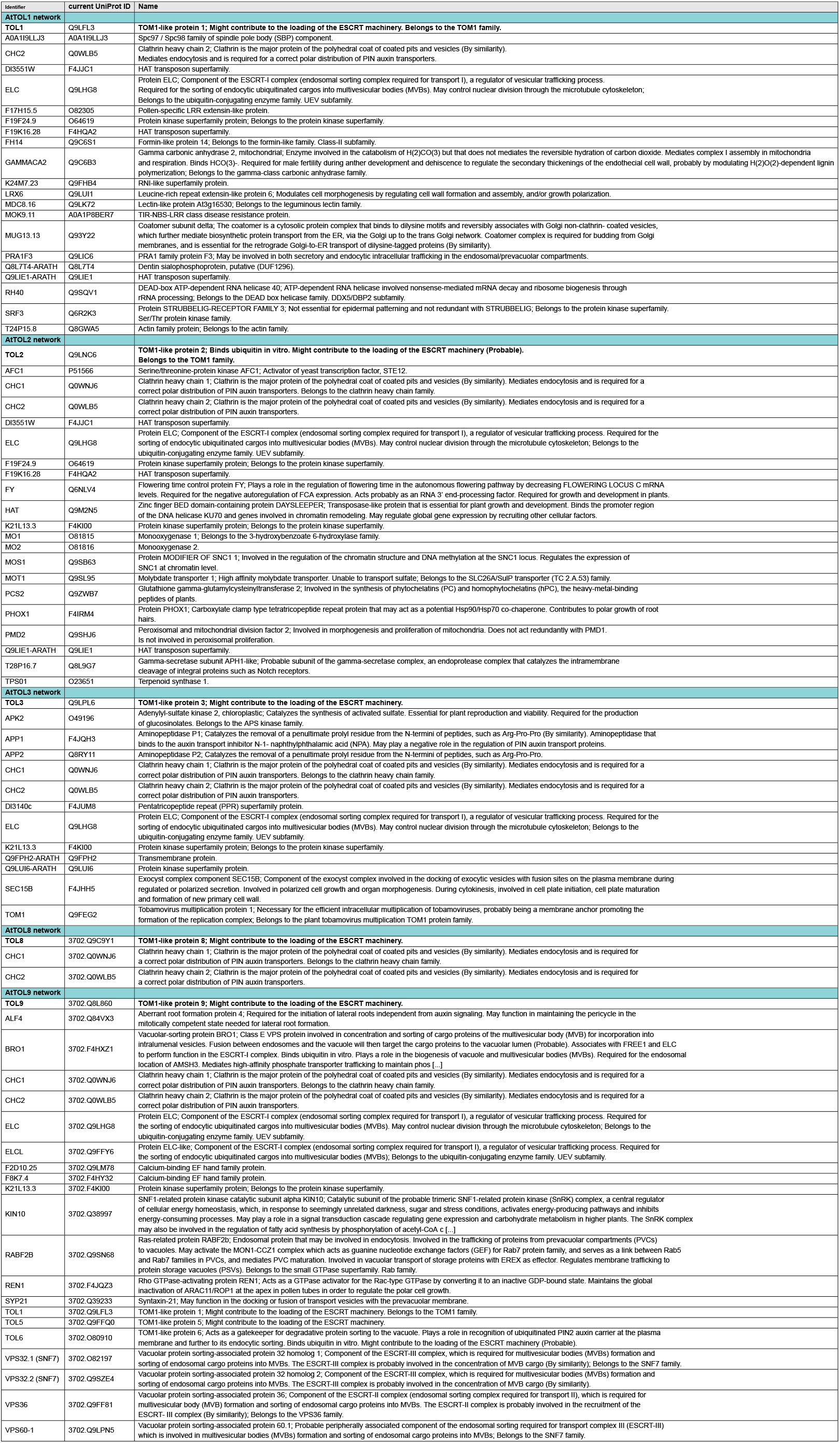
String data AtTOLs and interacting proteins.

These results demonstrate that barley root and shoot proteomes undergo extensive reprogramming during development and under salt stress, with clear tissue- and stage-specific regulation of proteins involved in growth, stress response, and cellular organization. In particular, stress- and defense-related proteins were strongly up-regulated under salt stress, while proteins associated with cytoskeleton, vesicle trafficking, proteolysis, and photosynthesis were largely down-regulated.

### Impact of salt stress on HvESCRT protein abundance

In a second approach we addressed, whether the abundance of the identified ESCRT proteins was altered after salt treatment. In total, we could identify and quantify eight ESCRT proteins (HvTOL1, HvTOL3, HvTOL9, HvVPS20, HvSNF7.1, HvSNF7.2, HvSAL1, and HvVPS4. The heat map of their LFQ intensity values reveals three clades **(Figure2 A)**: clade (a) with HvTOL9; clade (b) including HvVPS4, HvVPS46, and HvTOL3 that show decreasing abundance in roots facing EC30; and clade (c) including HvSNF7.1/2, HvVPS20, and HvTOL1 with a more diverse LFQ intensity value pattern. We next generated a Venn diagram of the significantly differentially regulated proteins identified in the previously discussed sample category comparison **(Figure2 B)**. It reveals HvSNF7.1 is specifically differentially abundant within 8STAP_vs_16STAP, while HvTOL1 and HvTOL3 are specifically differentially abundant within 16RTAP_vs_16REC30, indicating distinct spatio-temporal functional roles.

**Figure 2.**
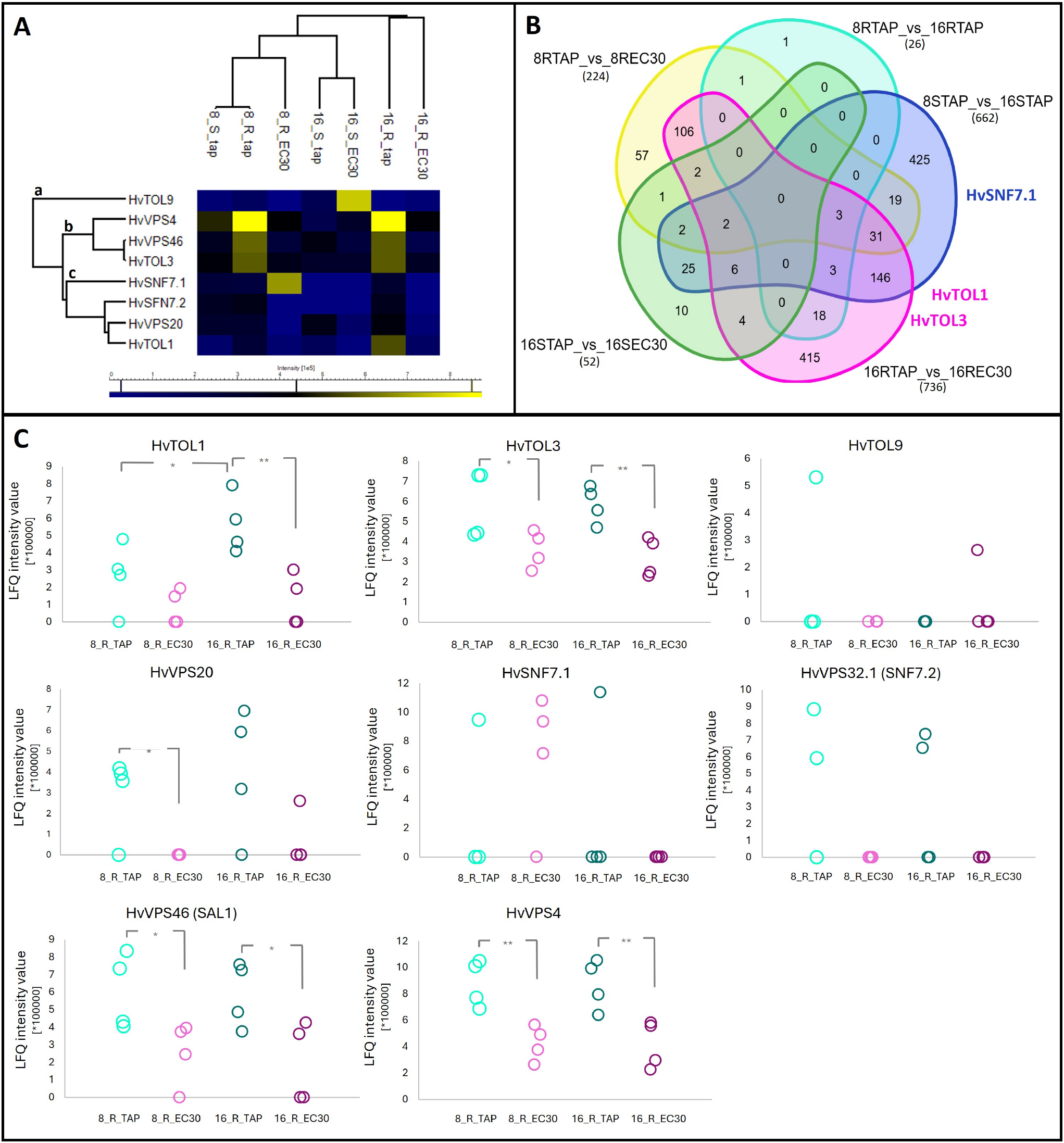
The abundance of ESCRT proteins is significantly regulated during development and salt stress. (A) Hierarchical cluster heat map of the LFQ intensity values of HvESCRT proteins in roots, identifying three clusters a, b, and c. (B) Venn diagram of the significantly altered proteins, upon sample category testing, during development or under salt stress. Different intersection groups are specifically analysed in **Figure S 6-8**. (C) LFQ intensity values of all identified ESCRT proteins in roots HvTOL1, HvTOL3, HvTOL9, HVVPS20, HvSNF7.1, HvSNF7.2, HvVPS46, and HvVPS4 for 8 and 16 DAS root samples of TAP and EC30 treatment. Significance level upon *t*-tests is depicted via connecting line and asterisks. n = 4, * Indicating a p-value of ≤ 0.05, ** Indicating a p-value of ≤ 0.01, *** indicating a p-value of ≤ 0.001 upon *t*-tests.

Next, the protein abundance of the identified HvESCRT-0-like, HvESCRT-III, and HvESCRT-associated proteins in roots was further analysed **(Figure2 C)**: The ESCRT-0-like protein abundance HvTOL1 is significantly up-regulated during root development and especially at 16 DAS significantly down-regulated under EC30. While HvTOL3 shows a rather constant abundance during root development and is as well significantly down-regulated under EC30. The protein levels of ESCRT-III protein HvVPS20 are up-regulated during root development and significantly down-regulated under EC30. Furthermore, the ESCRT-III protein HvSNF7.1 is high abundant at 8 DAS, but less abundant under EC30 at 16 DAS, whereas HvSNF7.2/HvVPS32.1 is constantly abundant during root development, but could not be detected during EC30. The ESCRT-III-associated protein HvVPS46/SAL1, and the ATPase HvVPS4 are constantly abundant during root development, but significantly less abundant during EC30. Altogether, these results suggest that the abundance of HvESCRT proteins is down-regulated by EC30.

### HvTOL quadruple- and double knock-out barley lines are lethal

TOL proteins have not yet been described to be involved in salt stress response. Our previous data show a temporal regulation of the protein abundance of HvTOL1, HvTOL2, HvTOL3, and HvTOL8 during barley grain development (Roustan et al., 2020), but we could not detect TOL proteins in our previous barley germination study during salt stress (Dermendjiev et al., 2021). We first performed bioinformatic analysis of the HvTOL family in barley. We identified 8 out of 9 known Arabidopsis TOL protein orthologs in barley, where HvTOL3 and HvTOL8 cluster in a different clade compared to HvTOL1 and HvTOL2 **(Figure3 A)**. Consistent with the phylogenetic tree, the protein alignments differ between HvTOL1/HvTOL2 and HvTOL3/HvTOL8 **(Figure3 B)**. However, the VHS (VPS-27, Hrs, and STAM) and GAT (GGA and Tom1) domains are conserved in all four HvTOL proteins. To analyse, whether they show a different localization behaviour, we generated HvTOL1-, HvTOL2-, HvTOL3-, and HvTOL8-GFP tagged constructs, regulated by the rice pActin promotor **(Figure3 C)**. Confocal microscopy of infiltrated tobacco leaves showed for HvTOL1, HvTOL2, and HvTOL8 show little signal in the cytosol but stronger putatively at the plasma membrane **(Figure3 C)**. HvTOL3 showed a strong signal in the cytosol and putatively at the plasma membrane **(Figure3 C)**.

**Figure 3.**
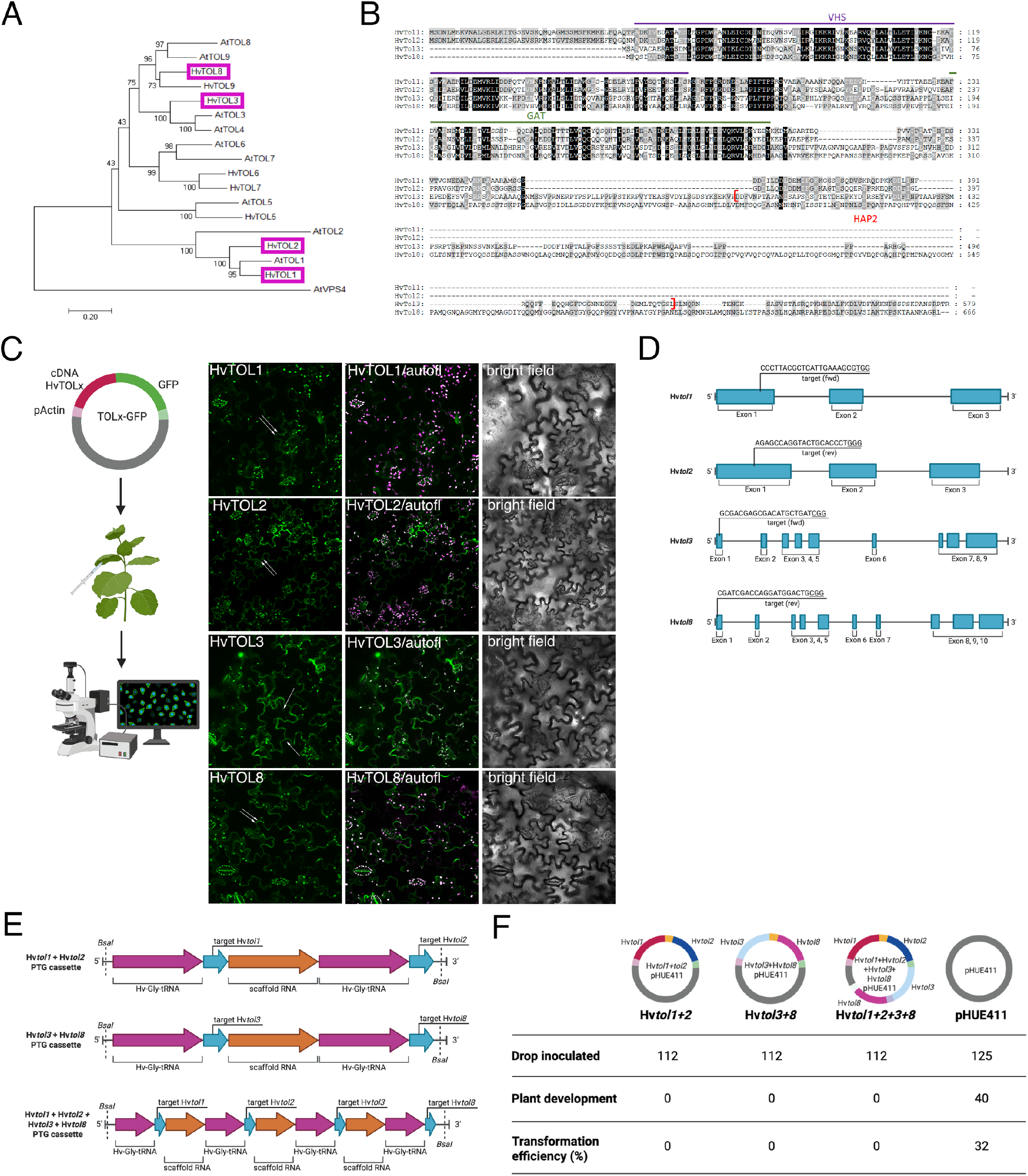
HvTOL1, 2, 3, and 8 localized differently and CRISPR Hvtoldouble and Hvtolqua mutants are lethal. (A) Phylogenetic tree with Arabidopsis and barley TOLs, with HvVPS4 as outgroup, generated by MEGA software, Bootstrap 1000. (B) Protein alignment of barley HvTOL1, HvTOL2, HvTOL3, and HvTOL8. Highly conserved regions in the alignment are in black 100 % and grey 80 %. In all four proteins we found a VHS domain which is important for membrane binding; and a GAD domain which is important for Ubiquitin binding. HvTOL3 additionally has a HAP2 domain. (C) Localization of HvTOL1-GFP, HvTOL2-GFP, HvTOL3-GFP, and HvTOL8-GFP in infiltrated tobacco leaves. Grafic designed with BioRender. Scale = 40 um. (D) Genomic DNA regions of HvTOL sequences with exons (blue) and indicated target sequences for genome editing by CRISPR/Cas9, the Protospacer Adjacent Motiv (PAM) sequences are underlined.(E) Schematic structures of polycistronic tRNA-gRNA gene (PTG) cassettes for multiplex genome editing to create Hvtolwins and Hvtolqua mutants. HvTOL target sequences, encoded gRNA scaffold region and the encoded barley Glycine t-RNA are indicated. BsaI restriction enzyme sites enable Golden Gate cloning into pHUE411 vector, which contains the required downstream gRNA scaffold with terminator.(F) Summary of approach to generate stable barley CRISPR mutants. Constructs used for drop inoculations are indicated. Number of transformed barley embryos are compared to developed plant material, resulting in the transformation efficiency. Hv*toldouble 1/2, 3/8* and Hv*tolqua* mutants are lethal since no plants could be generated, whereas 40 plants were positively screened for the empty pHUE411 plasmid.Table prepared with BioRender.

In Arabidopsis, it required *tol* quintuple knock-outs (*tolQ*) to produce pronounced phenotypes under normal conditions (Korbei et al., 2013). Therefore a barley quadruple CRISPR/Cas9 knock-out construct (Hv*tolqua*) of HvTOL1, 2, 3 and 8 was generated **(Figure3 D,E)**. Additionally double CRISPR/Cas9 knock-out constructs for (Hv*Tol1*) and (Hv*Tol2*) (Hv*toldouble 1/2*), and for (Hv*Tol3*) and (Hv*Tol8*) (*Hvtoldouble 3/8*) were generated **(Figure3 D,E)**.

Agrobacteria-mediated transformation of barley immature embryos (Hinchliffe and Harwood, 2019), for the generation of stable knock-out barley lines was performed. Several experimental rounds of transforming Hv*toldouble 1/2, Hvtoldouble 3/8*, and Hv*tolqua* constructs into barley embryos were performed, all consistently resulted in dying of the calluses. However, the negative control—transgenic barley plants generated with empty vector pHUE411—showed survival, and shoot and root development with a 32 % transformation efficiency, indicating, that Hv*toldouble 1/2, Hvtoldouble 3/8*, and (Hv*tolqua*) are lethal at this stage **(Figure3 F)**.

### HvTOL3 weakly interacts with HvTOL8

Arabidopsis *tolQ* plants, which lack five of the nine At*TOLs* (At*TOL2*, At*TOL3*, At*TOL5*, At*TOL6*, and At*TOL9*), exhibit severe developmental defects, suggesting interaction and coordination between family members (Korbei et al., 2013; Moulinier-Anzola et al., 2020). However, we were not able to generate knock-out lines in barley since generated CRISPR/Cas knock-out Hv*toldouble 1/2* and *3/8* and Hv*tolqua* lines are lethal.

Since protein levels of HvTOL3 were significantly down-regulated in EC30 at both developmental stages, we further focused on the characterization of HvTOL3. HvTOL3 showed a slightly distinctive localization pattern, different compared to the other HvTOLs, with stronger cytosolic signal, but also signal at the plasma membrane **(Figure3 C)**. Additionally, in Arabidopsis, AtTOL3 was shown to be phosphorylated by AtMPK6 (Rayapuram et al., 2018; Hoehenwarter et al., 2013). However, the single knock-out At*tol3* does not show any phenotype under normal growth conditions (Korbei et al., 2013).

To investigate putative interactions between HvTOL3 and further HvTOLs under normal growth conditions, we first conducted a STRING^1^ (Szklarczyk et al., 2023) online protein-protein association network and functional enrichment analysis for HvTOL proteins **(Figure S12, Table S2)**. STRING analysis for HvTOL1, HvTOL2, HvTOL3, and HvTOL8 provided a first selection of a potential interaction network, although it showed no specific HvTOL interaction network with HvTOL3. This was additionally assured by STRING analysis of AtTOL3 **(Figure S13, Table S3)**.

We further conducted bimolecular fluorescence complementation assay (BiFC) and yeast-two hybrid (Y2H) experiments to analyse the HvTOL interaction *in vivo* **(Figure4, Figure S14**,**15)**. Since Arabidopsis AtTOL3 was shown to be phosphorylated by AtMPK6 (Rayapuram et al., 2018; Hoehenwarter et al., 2013), we performed bioinformatic analysis, to reveal putative phosphorylation sites in HvTOL3 **(Figure S16)**. Subsequently, we chose AtMPK6 as potenital interactor, and AP2C1 as negative control for BiFC (Schweighofer et al., 2007). Indeed, a strong positive signal could be observed between HvTOL3 and AtMPK6, but not with the other tested HvTOLs when transiently expressed in tobacco **(Figure4 A)**. In contrast, no signal was observed between HvTOL3 and AP2C1, indicating the reliability of the BiFC analysis. Potential homodimerisation of each HvTOL2, 3, and 8 was found **(Figure14)**. However, only HvTOL1 and HvTOL3 showed putative heteromeric interaction with HvTOL8. Our Y2H analyses confirmed strongly the controls and indicated possible additional interaction of HvTOL3 and HvTOL8, thus no further interactions with other ESCRT-0-like proteins **(Figure4 B)**. Together the interaction studies provided us with a first glimpse of HvTOL interactions, indicating weak interactions of HvTOL3 with HvTOL8 and HvTOL8 with HvTOL1 under normal growth conditions **(Figure4 C)**.

**Figure 4.**
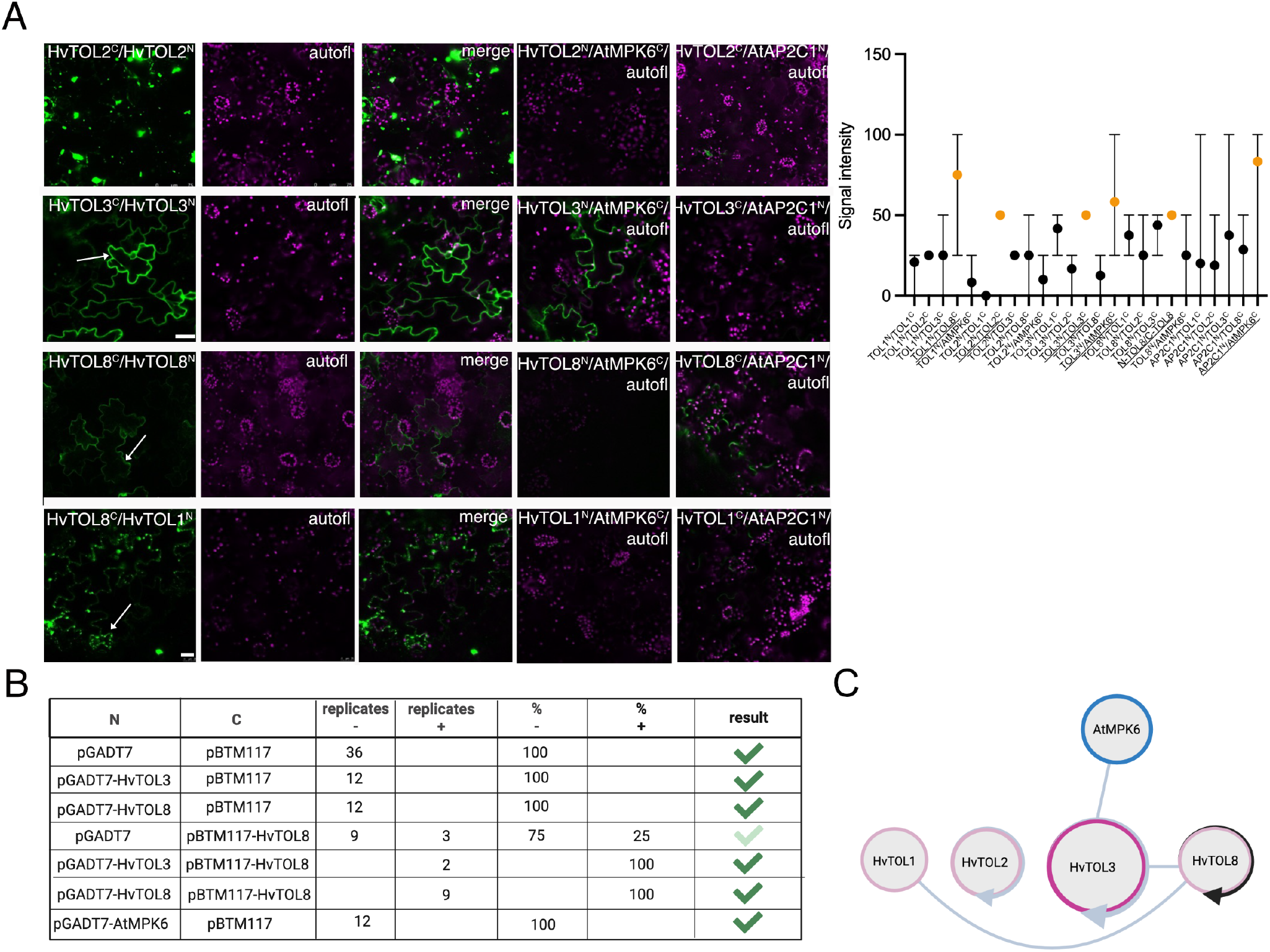
HvTOL3 weakly interacts with HvTOL8. (A) Different interaction results for BiFC in tobacco plants for HvTOL 1/3/8 upon split YFP combination on N(SPYNE) and C(SPYCE) term. Note the negative controls of HvTOLs N- or C-terminal combined with proteins AtMPK6 and AP2C1. 3 - 6 biological replicates (except HvTOL3^*N*^ -HvTOL3^*C*^ ; n is 1) were performed and the signal intensities were classified into no signal (y=0), low signal (y=25), middle (y=50), and strong (y=100). Positive interaction was assumed when mean was higher, or the same as 50 (labeled in orange). (B) Y2H interaction table. Interaction tests of different N and C terminal combinations (column one and two), column three and four show the amount of biological replicates, evaluated as negative or positive Y2H interaction. Column five and six represent the percentage of positively and negatively tested interactions. (C) TOL protein interaction schema. Interaction proven by both (BiFC and Y2H) experiments are shown with black lines, grey lines show the result from one experiment. Circle with arrow indicates homodimerization. Prepared with BioRender.

### At*tol3* mutants are sensitive to salt

Our generated CRISPR/Cas knock-out Hv*toldouble 1/2* and *3/8* and Hv*tolqua* lines are lethal. On the other hand, the single knock-out At*tol3* shows no phenotype under normal growth conditions. However, nothing is known about the role of TOL3 in abiotic stress response. Subsequently, we wanted to know if At*tol3* shows an altered growth behavior when subjected to salt stress. An ePLANT^2^ search indicated gene expression of At*TOL3* in Arabidopsis seed and root, but transcriptionally no up- or downregulation during salt stress (150 mM NaCl) in roots at seedling stage. Nevertheless, we subjected At*tol3* plants to 150 mM NaCl and analysed the germination rate, the root growth, and the degree of root curling (DC) (according to Wu et al. (2015)). We identified a delay in the germination rate of At*tol3* when germinated on ½ MS including 150 mM NaCl, analysing them at 5, 7, and 12 DAG (days after germination) **(Figure 5A)**. Next, we analysed the sensitivity of At*tol3* to 150 mM NaCl. We therefore subjected WT and At*tol3* plants, that grew 7 DAG on ½ MS for 7 DAG to salt stress and analysed the root growth rate per day and the DC. At*tol3* was sensitive to 150 mM NaCl since the root growth rate decreased and the DC ratio increased compared to WT **(Figure 5B-D)**. Taken together, these data indicate a NaCl-sensitive phenotype of the single knock-out At*tol3*.

**Figure 5.**
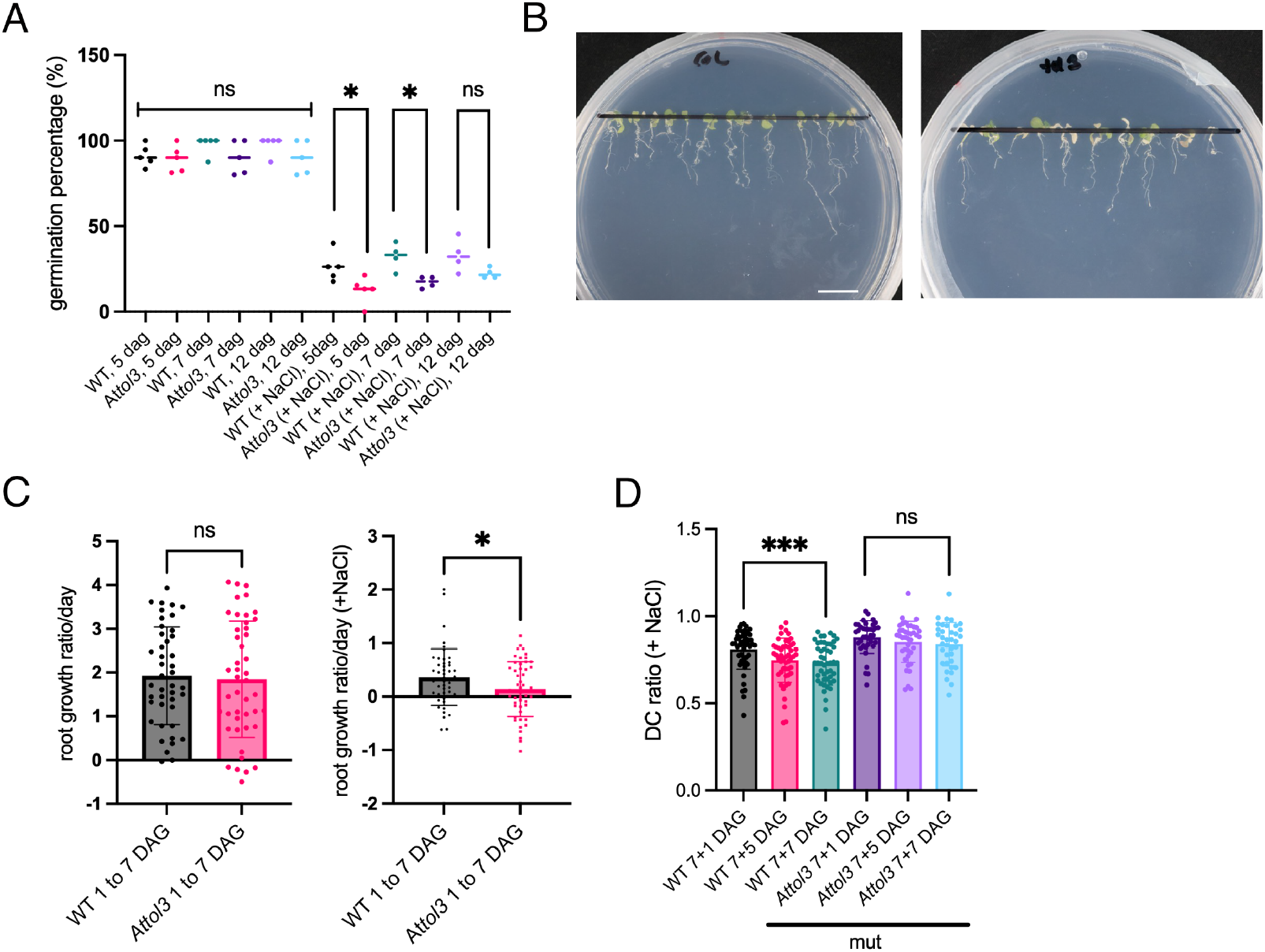
At*tol3* is sensitive to salt stress. (A) Germination rate of WT (Columbia) and At*tol3*, plated on ½ MS versus on ½ MS + NaCl. n = 75– 92, 4 biological replicates. (B) Representative picture of WT (Col) and At*tol3* at ½ MS + NaCl, 7+7 DAG. (C) Root growth ratio per day after growing 7 days on ½ MS, and replatet on ½ MS + NaCl, 1 to 7 DAG. n = 37– 50, 3-4 biological replicates. (D) Degree of root curling (DC) ratio, measured at three time points after replating: 7+1, 7+5, and 7+7 DAG. n = 37 – 50, 3-4 biological replicates. Significance level upon *t*-tests between certain groups is depicted via connecting line and asterisks. * Indicating a p-value of ≤ 0.05, ** Indicating a p-value of ≤ 0.01, *** indicating a p-value of ≤ 0.001 upon *t*-tests.

### Over-expression of Hv*SNF7*.*1* leads to hypersensitivity to EC30, and Hv*VPS4* is a positive regulator

SNF7.1 is drastically down-regulated in salt stressed Nicotiana tabacum (Garcia de la Garma et al., 2015), confirming partially our time-scale data, where HvSNF7.1 is up-regulated at 8 DAS, but down-regulated at 16 DAS EC30, and HvSNF7.2 are predominantly less or non abundant during EC30. We therefore investigated whether the over-expression line HvSNF7.1-mEosFP (HvSNF7.1 OE) (Roustan et al., 2020) shows altered response to EC15 **(Figure 6A, Figure S17A,B)**. Hv*SNF7*.*1* OE shows strong effect of EC15 on germination, since no grains germinated compared to the negative control, even though germination in TAP is not affected **(Figure S17C)**. This result indicates that HvSNF7.1 is a negative regulator in salt stress response. Zm*SKD1/VPS4* over-expression in tobacco plants shows enhanced tolerance to salt stress (Xia et al., 2013). Thus, if HvVPS4 is involved in salt stress response in barley, over-expression of Hv*VPS4* arguably would show altered response to salt stress treatment. We therefore generated a barley actin::HvmEosFP-VPS4 over-expression line (HvVPS4 OE). Positive T3 plants were subjected to EC15, subsequently root / shoot growth, and root angles were measured **(Figure 6A,B, Figure S17A,B)**. HvVPS4 OE showed a significant better shoot development compared to the negative control Hvactin::mEosFP (mEosFP-control) in EC15, however no better root development or significantly altered root angle were observed. Therefore, we propose HvVPS4 as a weak positive regulator to cope with salt stress. Altogether, these results suggest different, tissue specific response of HvSNF7.1 and HvVPS4 to changes in salinity.

**Figure 6.**
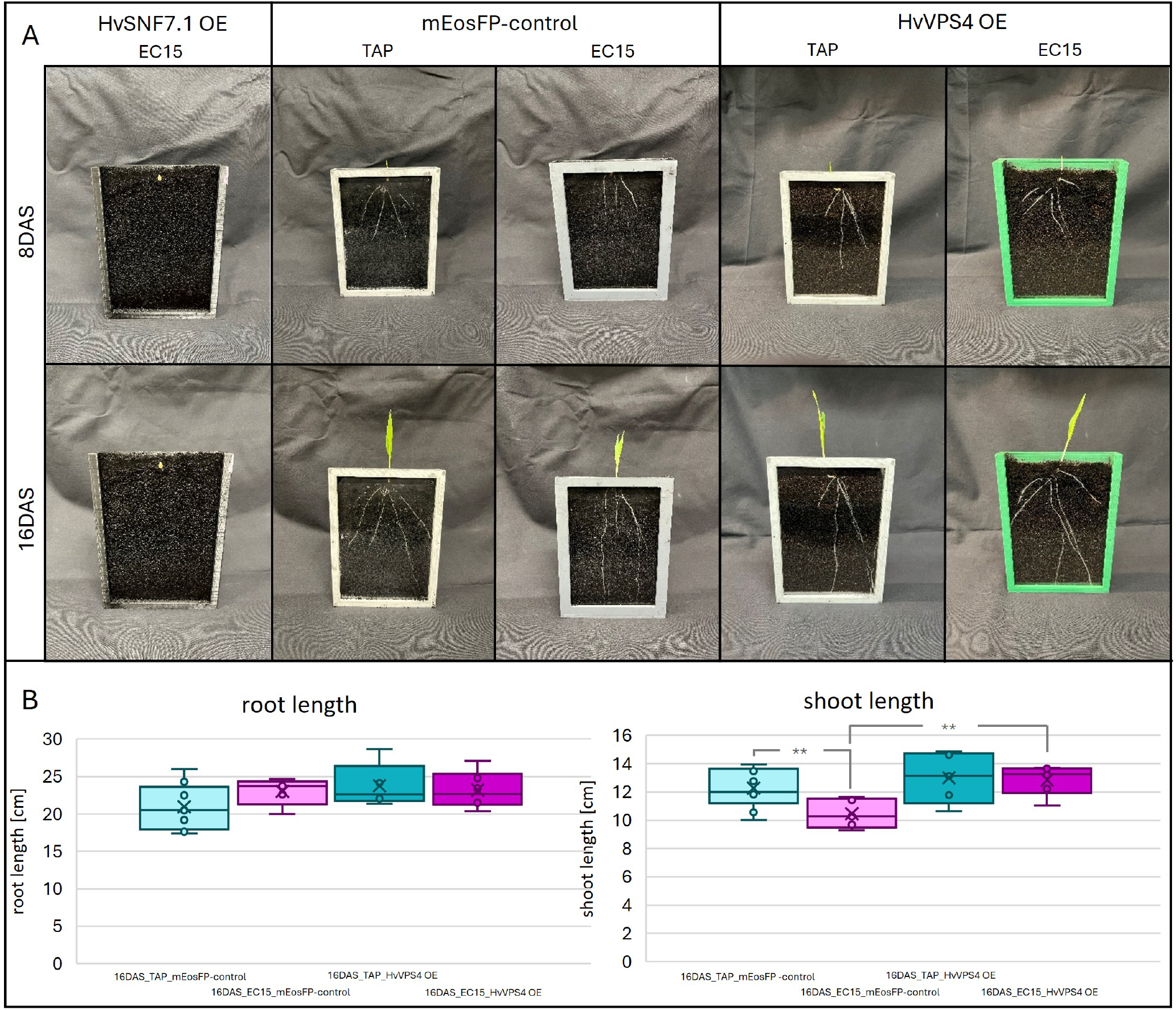
HvSNF7.1 and HvVPS4 are involved in salt stress response. (A) mEosFP (mEosFP-control), mEosFP-HvVPS4 (HvVPS4 OE), HvSNF7.1-mEosFP (HvSNF7.1 OE) plants grown for 8 DAS and 16 DAS in TAP or EC15 in rhizoboxes and DRD-BIBLOXes. (B) Boxplots of root and shoot length in cm of 16DAS mEosFP-control and HvVPS4 OE plants of TAP and EC15, ** indicating a p-value of ≤ 0.01 upon *t*-tests. n = 5-9.

## DISCUSSION

Increasing evidence shows that ESCRT proteins respond to abiotic stress (Liu et al., 2025; Zeng et al., 2023; Gao et al., 2017). In the present study, we identified ESCRT-0-like TOL3, ESCRT-III SNF7.1, and ESCRT-associated VPS4 proteins as key players in plant salt stress response. Although these proteins share the common feature of being down-regulated in response to salt stress, our data suggest a functional divergence in their biological roles. At*tol3* displayed delayed germination and altered root development under salt stress, demonstrating the necessity of a single TOL in salt stress response. Further, HvSNF7.1 seems to act as negative regulator, whereas HvVPS4 acts as positive modulator during germination and seedling development in salt stress response **(Figure 7)**.

**Figure 7.**
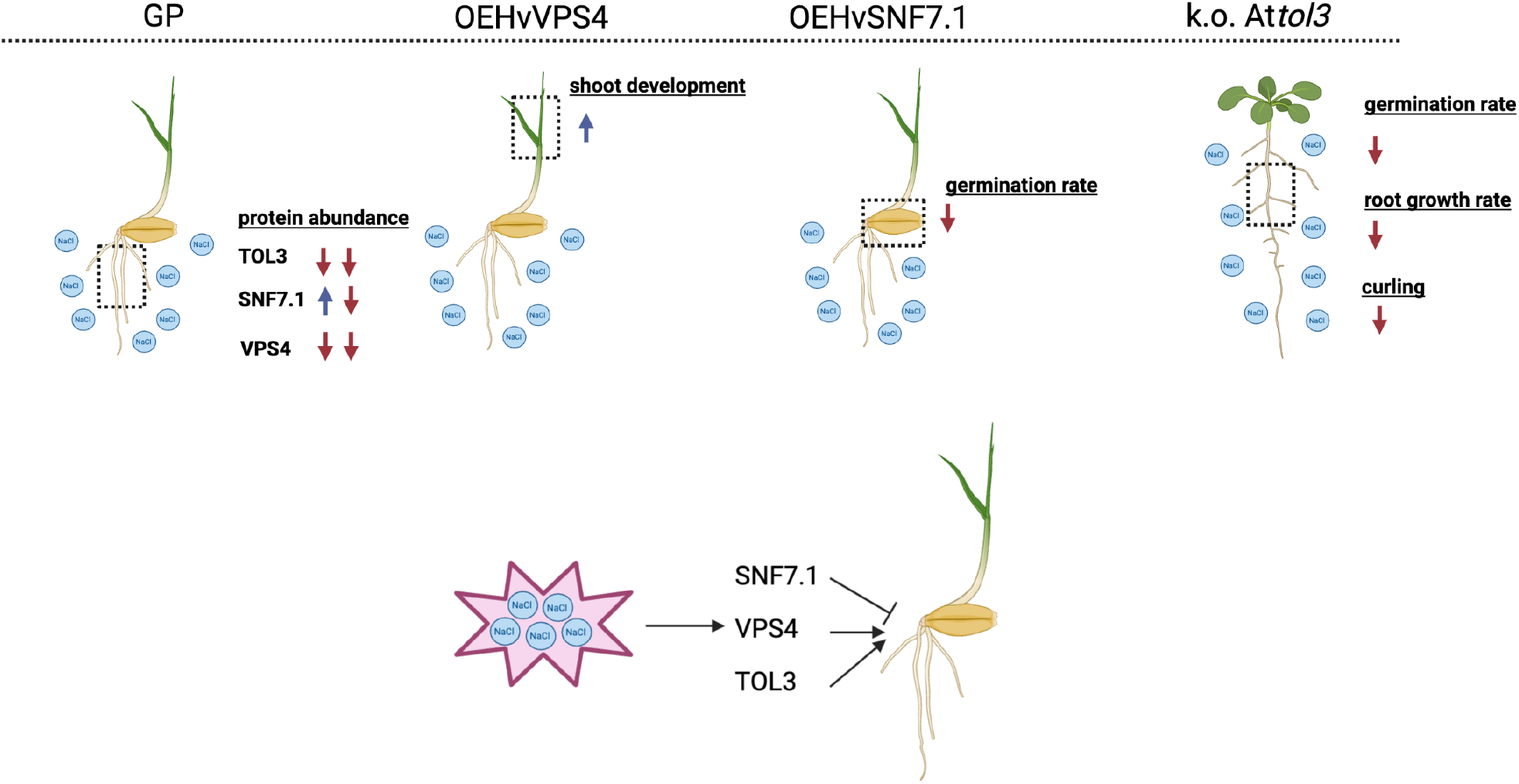
ESCRT proteins involved in salt stress response. Graphic display of how different ESCRT proteins (HvTOL3, HvSNF7.1, HvVPS4) are affected in their protein abundance during stress in the barley root (left). Display of increased shoot growth in HvVPS4 OE and reduction of germination rate in HvSNF7.1 OE under salt stress (middle two). Grafic of Arabidopsis At*tol3* show reduced germination rate, root growth and root curling under salt stress (right). SNF7 is illustrated as negative, VPS4 and TOL3 as positive regulators during salt stress (bottom). Prepared with BioRender.

Here, we applied high soil salt concentration to barley using our DRD-BIBLOX system, simulating near-natural soil conditions, and salt stress conditions. While being aware of its limitations, for this first experimental approach we solely used NaCl for stress experiments and we also included stress experiments on MS media. However, we herewith did first steps to perform stress application in conditions as near as possible to nature, to enhance the knowledge transfer to the field.

Although barley is a halophyte organism, salt stress disrupts its ion homeostasis and the metabolic processes essential for germination and seedling establishment (Munns and Tester, 2008; Dermendjiev et al., 2021; Xiong et al., 2024). Consistent with previous studies (van Zelm et al., 2020; Kumari et al., 2021; Fricke et al., 2006; Mostek et al., 2015), we observed that high soil salinity under near natural conditions led to delayed germination and reduced seedling growth in barley—likely consequences of osmotic and ionic stress, reduced root auxin levels, by limiting water uptake and enzyme activation, and therefore inhibited cell expansion (Liu et al., 2015; West et al., 2004; Munns and Tester, 2008). Further, high soil salinity altered the barley RSA. Specifically, we observed reduction in the seminal root growth angle *α* down to 50%, less seminal roots, and increased root hair density following salt stress. Those represent significant traits for salt stress adaptation and potentially help with deeper root growth and the plants access to less saline water in monocot plants, including modifying water and nutrient uptake zones - previously identified in wheat, maize, and rice (Alrajhi et al., 2024; Shelden and Munns, 2023; McCormack et al., 2015; Li et al., 2021). Proteomic analyses have identified proteins involved in the salt stress response, including ion transporters and channels, and stress-responsive proteins, which are crucial for maintaining ionic balance and facilitating detoxification under saline conditions (van Zelm et al., 2020; Mostek et al., 2015).

In this study, proteomic analysis of barley seedling revealed drastic, tissue-specific, salt stress-responsive protein dynamics. During root development, proteins linked to growth, cell wall biosynthesis, and vesicle trafficking remained abundant, as expected for the active build-up required for root maturation (Siqueira et al., 2022). Shoots showed energy allocation by temporal regulation of translation and photosynthesis-related proteins, with late-stage down-regulation possibly shifting resources to defense (Bilgin et al., 2010). We found distinct proteomic changes, especially in roots, including reduced levels of defense- and stress-related proteins, as well as ABA- and auxin-related proteins, such as the ABA receptor HvPYL2. Vesicle-associated proteins, particularly clathrin-independent components, were mostly down-regulated, while some vacuolar transport proteins increased in their abundance. These changes, alongside decreased actin and tubulin levels, suggest impaired trafficking and structural adaptation under salt stress (Garcia de la Garma et al., 2015; Dermendjiev et al., 2021). Concurrent up-regulation of cellulose synthases and sugar transporters, together with thicker roots, point to stress-induced reinforcement of cell walls (Byrt et al., 2018).

Under normal conditions, ESCRT proteins have previously been described to be involved in root growth regulations, development, and architecture, as mutation of the ESCRT-III component AtVPS2.2, localized in the root meristem, cause a strong root growth phenotype in Arabidopsis (Ibl et al., 2012).

In addition, several ESCRT proteins have been described to be involved in plant stress response (Gao et al., 2017), where ESCRT proteins regulate salt stress response via several routes. First, they facilitate targeted degradation or recycling of membrane proteins (such as ion transporters) (Jou et al., 2013; Babst and Odorizzi, 2013). Second, they modulate key signaling pathways for salt stress tolerance, e.g., ESCRT-I component VPS23 and ESCRT-III component FYVE are involved in positively regulating salt resistance by enhancing the interactions in the SOS pathway (Liu et al., 2025). Third, ESCRT components (ESCRT-I, II, III) vacuolar protein sorting-associated proteins (VPSs), and SNARE proteins, are upregulated in response to salt stress, indicating a conserved role for ESCRT-mediated vesicular trafficking in salt adaptation across photosynthetic organisms. This involves regulating membrane protein turnover and stress signaling receptors (Zhang et al., 2023). Fourth, ESCRTs are involved in autophagy-related processes upon stress (Mosesso et al., 2024). The plant-unique ESCRT component FREE1 (FYVE DOMAIN PROTEIN REQUIRED FOR ENDOSOMAL SORTING 1), is involved in endosomal sorting and vacuolar biogenesis. However, *free1* mutants accumulate autophagosomal structures in roots, suggesting ESCRT-related membrane trafficking influences root cell homeostasis and potentially responses to environmental cues influencing root system architecture (Zeng et al., 2023; Gao et al., 2015).

Here, we provide evidence that TOL3, SNF7.1, and VPS4, are key players in salt stress response. But what are their biological roles within this process? Although direct studies on ESCRT proteins specifically controlling root angle formation, halotropism (salt-avoidance tropism), or gravitropism are limited, the involvement of ESCRT proteins in hormone signaling, vesicle trafficking, and membrane remodeling hints at an indirect, but critical role in these processes. For example, ESCRT machinery impacts the trafficking of plasma membrane proteins and receptors, that perceive environmental and hormonal signals, governing root directional growth (Hsu and Jauh, 2017). The ESCRT-III complex is involved in membrane deformation and scission events, critical for vesicle-mediated transport and signaling regulation processes, essential for cellular polarity and directional growth responses, such as gravitropism and halotropism (Otegui and Spitzer, 2008; Henne et al., 2011; Cai et al., 2014).

The predicted HvTOL interactomes suggest divergent functions from AtTOLs, but support involvement in endocytosis, auxin modulation, and stress signaling via clathrin, Rab-GAPs, and MAP kinases like AtMPK6. STRING^1^ analysis indicates that AtTOL9 interacts with AtTOL6 and AtTOL5, suggesting functional redundancy rather than a direct interaction network. HvTOL3 may interact with HvTOL8 under non-stress conditions. While TOL8 expression in Arabidopsis is limited to siliques and flowers (Moulinier-Anzola et al., 2014), its barley ortholog is detected in developing grains but not in seedlings^3^. Double mutants involving At*tol8* and either At*tol5* or At*tol7* exhibit aborted seed development, suggesting a tissue-specific function for TOL8 (Korbei et al., 2013). Further, transient expression studies showed HvTOLs localizing to cytoplasm, plasma membrane, and vesicles, supporting roles in endocytosis and multivesicular body formation. We state the question if TOL3 is a distinct key player in root salt stress response and / or halotropism. Halotropism, a directional root growth response away from high salt concentrations, driven by asymmetric auxin distribution via PIN proteins, contrasts with the salt stress response, which involves physiological adaptations like ion homeostasis, osmoprotectant synthesis, and antioxidant defense to tolerate saline conditions (Galvan-Ampudia et al., 2013; Ma et al., 2022). In recent publications, no single knock out plant of the nine AtTOL family members shows an obvious phenotype (Korbei et al., 2013). While in Arabidopsis, knock-outs of multiple *TOL* genes *(tol2/3/5/6/9)* display strong phenotypes under standard conditions (Moulinier-Anzola et al., 2020; Korbei et al., 2013). Interestingly, quadruple mutants *(tol2/3/5/6)* lack obvious phenotypes under normal conditions but show increased drought tolerance and ABA sensitivity (Moulinier-Anzola et al., 2024). In this study, HvTOL1 and HvTOL3 decreased significantly in roots under salt stress **(Figure7)**.

Gravitropism, regulated by columella cells in the root tip, is disrupted by high salinity (Sun et al., 2008; Su et al., 2017). Negative halotropism— roots growing away from salt—is common in salt-sensitive species, while halophytes like *Bassia indica* and *Limonium bicolor* display positive halotropism (Shelef et al., 2010; Leng et al., 2019). In crops, it is still unclear whether true halotropic responses exist, or whether root behaviour under high salinity reflects limited water availability (Munns and Gilliham, 2015). In Arabidopsis, negative halotropism is salt-specific and involves auxin redistribution regulated by PIN2, not triggered by equivalent osmotic stress (Galvan-Ampudia et al., 2013). This directional auxin flow involves changes in PIN2 and AUX1 localization in the root tip (van den Berg et al., 2016). Over-expression of SOS1 reduces halotropic response, while sos1 and sos2 mutants are hypersensitive to NaCl (Galvan-Ampudia et al., 2013), suggesting that intracellular Na^+^ may serve as a signal—though not yet proven. In wheat, genotype-specific differences in root Na^+^ sequestration correlate with salt tolerance (Wu et al., 2018), but whether Na^+^ levels directly influence halotropism is unknown.

Here, we monitored the salt sensitivity of At*tol3*. Interestingly, At*tol3* showed altered root growth and DC **(Figure 7)**. Known to bind ubiquitinated cargo and regulate auxin/ABA signaling, HvTOLs likely contribute to root adaptation mechanisms. However, the role of HvTOL3—putatively together with HvTOL8—in salt stress response and / or halotrophic signaling in barley, or other crops, remains to be investigated, and single barley knock-outs will be critical for further insights. We could detect an up-regulation of HvSNF7.1 at 8 DAS in EC30 roots, but a down-regulation at 16 DAP, indicating that salt stress has a stage-specific effect on the protein abundance of HvSNF7. In a previous study, we showed that HvSNF7.1 is highly abundant in early stages during barley grain development, but decreases at later stages, and has not yet been identified in mature grains, with a putative role in seed storage protein transport (Roustan et al., 2020). So far, SNF7.1 has been described as key player in plants during autophagosome closure and MVB body formation (Zeng et al., 2023; Buono et al., 2017). Meanwhile, we found Hv*SNF7*.*1* over-expression in the grain impaired germination under salt stress, but not under normal conditions. Since germination relies on autophagic processes (Yang et al., 2016) and autophagy is highly induced by salt stress (Luo et al., 2017), over-expressed *SNF7* may alter its autophagosomal functions and therefore impact germination process. Thus, we propose that Hv*SNF7*.*1* over-expression acts as a stress-induced negative regulator. **(Figure 7)**

Conversely, VPS4 is acting very diverse in abiotic stress response. The expression of *VPS4* is up-regulated after salt stress in the halopyhte ice plant *M. crystallinum* (Mc) (Jou et al., 2013). Additionally, *M. crystallinum* copin1 (McCPN1), a Ring-type ubiquitin ligase, is interacting with the phosphorylated (by *M. crystallinum* sucrose non-fermenting 1-related protein kinase, McSNRK1) McVPS4, therefore McVPS4 changes its location from the cytosol to the plasma membrane, forming a complex network, which putatively leads to modulation of ATPase activity. Further, over-expression of the dominant-negative AtSKD1^*E*232*Q*^ (VPS4) results in plant lethality (Haas et al., 2007), and down-regulated AtSKD1/VPS4 shows reduced salinity response and altered root development with a loss in Na^+^/K^+^ homeostasis (Ho et al., 2010). Furthermore, over-expression of *ZmVPS4* in tobacco plants showed less chlorosis and faster root growth when subjected to salt stress (Xia et al., 2013). Additionally, *ZmVPS4* was suggested as regulator of the reactive oxygen species (ROS) signaling network under salt stress. Lyst-Interacting Protein 5 (LIP5), involved in late endosome to vacuole transport via MVB sorting pathway, seems to play a critical role in stress response as positive regulator of VPS4 (Haas et al., 2007). However, in our data, HvVPS4 is down-regulated in salt stressed barley roots **(Figure 7)**. It is worth to mention, that we did not detect the barley ortholog (according to Bar Browser^3^ AK363748, Uniprot F2DAX9) of AtLIP5 (At4g25750/Uniprot Q9SZ15), but an other Pistil-specific extensin-like protein UniProt F2E896, as well as we detected barley orthologs of SNRK1 (A0A8I6XRU5), which was down-regulated during salt stress, and CPN1 (F2DMD1), which was up-regulated during salt stress. Conversely, the abundance of HvVPS4 increased during grain development (Roustan et al., 2020). Consistent with that observations, Hv*VPS4* over-expression improved germination and growth under salt stress, despite endogenous down-regulation, suggesting a positive regulatory role of VPS4. It remains to be investigated whether HvVPS4 is interacting in a complex together with HvLIP5, HvSNRK1, and HvCPN1 in barley under salt stress conditions.

Altogether, our results show that TOL3, SNF7.1, and VPS4 exhibit non-redundant functions in salt stress response, indicating a spatio-temporal functional divergence role during salt stress response.

## MATERIAL and METHODS

### Plant material and growth conditions

The spring barley (*Hordeum vulgare*, L.) variety Golden Promise (GP) was used for all barley experiments. Plants were grown in a Conviron Adaptis A1000 at 14°C/12°C day(12h)/night(12h), with light intensities of 130–220 µmol m^−2^ s^−1^ and 70% humidity. Barley seeds were germinated in rhizoboxes covered by DRD-BIBLOXes (Dermendjiev et al., 2023). Rhizoboxes were filled with substrate that contained Cocopeat (3 mm sieved) mixed with activated charcoal powder at 3:1 with H_2_ O for control (TAP), or EC30 solution [340 mM NaCl in H_2_ O, electric conductivity of 30 mS/cm]. The soil moisture was set at a pF-value of 2-3 during the first 16 days of grain germination and plant growth according to Dermendjiev et al. (2023).

For salt treatment in Arabidopis during germination, sterilized seeds were plated directly on 1/2 MS, 1% sucrose supplemented with 150 mM NaCl, stratified for two days at 4 °C in darkness, and grown at 22 °C under long-day conditions with a photoperiod of 12 h light / 8 h dark. For salt sensitive analysis, sterilized seeds were plated on 1/2 MS 1% sucrose, and stratified for two days at 4 °C in darkness. After 7 days cultivation at at 22 °C under long-day conditions with a photoperiod of 12 h light / 8 h dark, plants were transferred to 1/2 MS 1% sucrose supplement with 150 mM NaCl and grown for further seven days.

### Phenotyping

Grains were germinated and seedlings were grown for 8 or 16 days after sowing (DAS). After 8 or 16 DAS rhizoboxes were photographed in a standardized way. Pictures allowed phenotyping and analysis of RSA. Further the number of visible roots as well as root angles (*α, β, γ, δ*, and *ϵ*) were recorded and measured. Additionally pictures of the root development from 0 DAS to 16 DAS were taken every 4 min, in the dark, with IR light, using a raspberry pi-DRD-BIBLOX camera setup as described in (Dermendjiev et al., 2023). Further plants were harvested at 8 and 16 DAS, soil was removed, pictures of seedlings were taken, actual root number and the total root and shoot lengths, and root diameters were measured. Subsequently the root and shoot material was separated and wet and dry weight was measured, after 70 h of drying at 60 °C. Furthermore, germination assays were conducted in in dark at 14 and 21 °C, in Petri dishes with filter paper in TAP or EC30 conditions. Phenotyping data and images were analysed using ImageJ and Microsoft Excel.

### Sample preparation and protein extraction

8 or 16 DAS shoots (S) were cut under regular light conditions and immediately frozen in LN_2_. Roots (R) were harvested in dark with dimmed red light, soil was removed and material was immediately frozen in LN_2_. Root or shoot material of 14 plants for 8 DAS and seven plants for 16 DAS were pooled to comprise one biological replicate each. Three to four biological replicates were prepared per biological sample category (8 DAS_R_TAP, 8 DAS_R_EC30, 8 DAS_S_TAP, 8 DAS_S_EC30, 16 DAS_R_TAP, 16 DAS_R_EC30, 16 DAS_S_TAP, 16 DAS_S_EC30). Plant material was homogenized with mortar, pestle, and LN_2_. 350 mg of fresh plant powder was used per sample to extract proteins. Protein extraction, digestion and desalting protocols were followed according to (Dermendjiev et al., 2023).

### LC-MS/MS analysis

Liquid chromatography and tandem mass spectrometry analysis was performed on a nano-LC-system (Ultimate 3000 RSLC; Thermo Fisher Scientific), coupled to an Impact II high-resolution quadrupole time-of-flight (Bruker), using a Captive Spray nano-electrospray ionization source (Bruker Daltonics) according to Dermendjiev et al. (2023).

### Data analysis and visualization

#### Maxquant processing

After LC-MS/MS, raw files were processed using the MaxQuant software (version 2.4.2.0) (Cox and Mann, 2008). Peak lists were compared against the barley reference proteome (*Hordeum vulgare*, L. subsp. vulgare (domesticated barley), cv. Morex, UniProt, UP000011116, version July 2023) using the built-in Andromeda search engine (Cox et al., 2011).

Settings for the MaxQuant processing included trypsin digestion as enzyme specificity and up to two missing cleavages were allowed. N-terminal acetylation and methionine oxidation were picked as variable modifications and cysteine carbamidomethylation was set as fixed modification. 248 common contaminants and decoy sequences were automatically added during processing. The false discovery rate (FDR) at peptide and protein level was set to 1%. Proteins were quantified across samples enabling the label-free quantification (LFQ) algorithm (Cox et al., 2014), and the match-between-runs option. Uncharacterized proteins were identified and manually annotated, post processing using UniProt BLAST application.

#### Data analysis and visualization using Perseus software and Microsoft Excel

Data previously processed with MaxQuant were subsequently analysed and visualized using Perseus (v.2.0.10.0) (Tyanova et al., 2016) and Microsoft Excel **(Supplement Data)**. LFQ intensity values were used for analysis. Raw data preparation prior to analysis included removing proteins with LFQ reads of 0 for all biological samples as well as proteins where three out of four or two out of three values of biological replicates of one sample group were missing. Rows where proteins were only identified by site were removed and reverse hits were excluded too. Further contaminants were excluded. Subsequently the dataset was transformed using log2(x) function. And missing values were replaced by random numbers drawn from a normal distribution of 1.8 standard deviation down shift and with a width of 0.3 of each sample. Next the data was transformed into a normalized expression matrix via substraction function. At this stage a PCA helped to visualize the whole dataset.

Additionally multiple sample ANOVA was performed with FDR of 1% and z-scores of significant data and overall data were used to map hierarchical cluster heat maps. Further *t*-tests helped to find significant differences regarding LFQ values between mean values of different sample groups (8RTAP_vs_16RTAP; 8STAP_vs_16STAP; 8RTAP_vs_8REC30; 16RTAP_vs_16REC30; 16STAP_vs_16SEC30), with two-tailed distribution and FDR of 0.05. Those were again visualized via hierarchical cluster heat maps.

Proteins that significantly differed in their abundance when comparing the mentioned sample groups were classified according to their cellular component, biological process and molecular function affiliations. Additionally protein groups of special interest, independent of significance, were picked to investigate further. All identified proteins among the groups were visualized regarding their LFQ trend in developing root tissue and under salt stress. Hereto means of z-scores over all biological replicates were used. **(Supplementary Data)**

### Cloning strategies

HvTOL-GFP constructs were designed with the long term aim to produce GFP-tagged barley TOL proteins in different organisms (*E*.*coli, Agrobacterium tumefaciens; A. tumefaciens, Saccharomyces cerevisiae strains*; Sc., and plants). In silico cloning was performed using SnapGene® and VectorNTI software. Coding sequences for *HvTOL1* (AK251767.1; XM_045095615.1), *HvTOL2* (AK250440.1; XM_045115633.1), *HvTOL3* (AK368118; XM_045124630.1; XM_045124629.1) and *HvTOL8* (AK252335.1; XM_045100432.1), were chemically synthesized^4^ and cloned into pGEX-4T-1 vector with codon optimization and removal of common internal restriction enzyme sites by silent mutations. For yeast two hybrid experiments the HvTOL sequences were subcloned into pGADT7, or pBTM116/7 vectors, sequenced, and transformed into Sc strains L40 (Vojtek et al., 1993). For BiFC experiments in *Nicotiana tabacum; N. tabacum* the HvTOL sequences were subcloned into pGreenII0029 based vectors (Roustan et al., 2020; Schweighofer et al., 2014; Hellens et al., 2000), sequenced, and transformed into *A. tumefaciens* strain GV3101 via electroporation. For expression in barley (*H. vulgare*) the HvTOL sequences were subcloned into pSB227-GFP vector (Kapusi et al., 2007), sequenced, and transformed into *A. tumefaciens* strain GV3101 by electroporation. For cloning of mEosFP-HvVPS4 the coding sequence of HvVPS4 was chemically synthesized based on *H. vulgare* Haruna nijo AK359160.1, and cloned into pBluescript II SK(+) by GeneCust^4^. Subcloning into pSB227-mEosFP (Kapusi et al., 2007; Hilscher et al., 2016), including sequencing, was used to obtain a mEosFP-HvVPS4 binary plant expression vector.

For the CRISPR/Cas9 construct design, three different polycistronic tRNA-sgRNAs (PTGs) were designed accodring to Gasparis and Przyborowski (2020), to aim double or quadruple HvTOLs gene target combinations *HvTOL1*/*2, HvTOL3*/*8*, or *HvTOL1*/*2*/*3*/*8*. CRISPR/Cas9 strategy aimed for non homologous end joining, a subsequent frame shift and knock-out. PTGs guide Cas9 to cut in first exons of the genes. For target sequence selection and designing of sgRNAs following online resources were used: chopchop^5^ (Labun et al., 2019) (barley reference genome Morex V2); crispor^6^ (Concordet and Haeussler, 2018) (barley reference genome GPv1); RNAWebSuite^7^ for RNAfold gRNA secondary structure predictions. IPK Barley BLAST Server^8^; gmapper^9^ GP genome blast server.

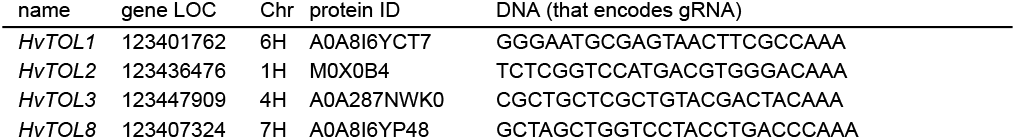

The corresponding PTG cassettes (flanked by BsaI restriction sites) were chemically synthesized and cloned into pCDFDuet-1 vector^4^. PTGs were subcloned into the pHUE411 vector (a gift from Qi-Jun Chen (Addgene plasmid #62203^10^; RRID:Addgene_62203)) (Xing et al., 2014) via BsaI/BsaI restriction enzyme sites and Golden Gate cut/ligation reaction. All constructs were screened and confirmed by sequencing. Constructs were transformed into an Agrobacteria tumefaciens strain GV3101 via electroporation and transformed into barley embryos (see Material and Methods, Agrobacteria Drop inoculation).

### Transient transformation by agroinfiltraton into *N. tabacum*

*N. tabacum* plants were grown for 4 weeks on soil, at 26/19 °C day/night, 12 h light cycles [4000 lx] with a humidity of 50-60%. Over night *Agrobacterium tumefaciens* cultures GV3101-pMP90-pSoup (Roustan et al., 2020; Hellens et al., 2005), transformed with the respective constructs were grown in LB with Tet, Rif, Gent, Spect/Kan at 28 °C to an OD_600_ of 1.0-1.2. Cultures were spinned down, the pellets were resuspended in infiltration media [100 mM MES, 10 mM MgSO_4_, 150 *µ*M Acetosyringone, pH5.6 (NaOH)], set to OD_600_ of 0.4, incubated at RT, 190 rpm for 3 h. *Nicotiana tabacum* leaves were infiltrated with the *Agrobacterium tumefaciens* solution using syringes without needles. Co-infiltration with HCpro increased leaf protein product. (Tyurin et al., 2017) Infiltrated plants were grown further for 3-4 days.

### Microcopy

Infiltrated *Nicotiana tabacum* leaves were analysed via live cell imaging using a Confocal Leica SP8 microscope. Leaf sections were mounted in tap water and imaged using an Argon laser and emission filter settings of 497 nm - 537 nm for the GFP signals and 661 nm - 747 nm for autoflourescence detection. Fluorescence signals of leaves infiltrated with either GFP-tagged HvTOLs were compaired against negative controls of not infiltrated leaves and leaves that were infiltrated with only infiltration media. Additionally, pSB227-*GFP* (Kapusi et al., 2007) was used as a control construct.

For BiFC analysis, YFP signal upon protein interaction was checked using a Confocal Leica SP8 microscope with an argon laser and emission filter settings of 530 nm - 587 nm for the YFP signals and 608 nm - 695 nm for auto-fluorescence detection. 3 - 6 biological replicates (except HvTOL3^*N*^ - HVTOL3^*C*^ ; n=1) were performed and the signal intensities were classified into no signal (0), low signal (25), medium (50), and strong 100. Positive interaction was assumed when the mean was higher or the same as 50.

### Generation of transgenic barley plants

Transgenic mEosFP-HvVPS4 plants were generated as described in Roustan et al. (2020). T1 plants surviving hygromycin selection were genotyped using primer pairs (TTGCCTCACAGATTCTCCCAC, CGATCTAGTAACATAGATGA CACC). T3 plants were used for root growth analysis.

For the generation of CRISPR plants, *A. tumefaciens* drop inoculation was used to transform isolated embryos with positively screened colonies for each construct according to Hinchliffe and Harwood (2019).

### Phylogeny assessment

Barley TOL proteins were identified using NCBI^11^ and UniProt^12^ databases. The phylogentic tree and alignment was done using MEGA software^13^ (version MEGA 12). Bootstrap 1000. HvVPS4 was used as outgroup.

### STRING Database analysis

Potential interaction partners of barley and Arabidopsis TOL1, TOL2, TOL3, and TOL8 were predicted using STRING, an online protein data base and platform^1^. Settings included evidence based network edges, a minimum interaction requirement score of 0.4, and an output of no more than 20 direct interactors. The protein IDs were then manually updated using UniProt and the Blast function.

### Yeast transformation and Y2H

pGADT7- and pBTM116/7 vectors harboring indicated sequences, were co-transformed into yeast strain L40 (Vojtek et al., 1993) and positive transformants were selected by auxotrophy on SD plates, lacking Leucin (Leu) and Tryptophan (Trp). Different HvTOL protein interactions were tested, pBTM116-AtMPK6 and pGADT7-AP2C1 combinations were used as positive control. For reporter gene assay transformed cells were spread onto SD-plates lacking Leu, Trp and His, including 3-Amino-1,2,4-triazole with indicated concentration (8mM, 10mM).

## ACKNOWLEDGEMENTS

We thank the master gardeners Thomas Joch and Andreas Schröfl and their team for the continuous care and maintenance of our plants and for taking care of steps around substrate preparation and rhizobox filling. Microscopy was performed at the Core Facility Cell Imaging and Ultrastructure Research, University of Vienna—a member of the Vienna Life-Science Instruments (VLSI). We thank Stefan Plott for critically reading the manuscript.

## AUTHOR CONTRIBUTIONS

VI conceived the project, VI, AS, JH, and MS designed and performed the experiments. TN, GM, and DL performed the LC-MS/MS. MS performed the data analysis. PF, SD, LK, and LS assisted with the cloning experiment. SD and LS assisted with Y2H experiments. PF and LK assisted with BiFC experiments. LK and MS generated and characterized HvTOL CRISPR lines. JH designed and performed the cloning of the actin::mEosFP-VPS4 construct and EK created transgenic barley over-expression lines. BK provided Arabidopsis knock-out seeds and provided valuable experimental advice. AB assisted with the Arabidopsis salt stress experiments. MS and VI wrote the manuscript, AS revised and finalized the manuscript. All authors contributed to the article and approved the submitted version.

## SUPPLEMENTARY MATERIAL

**Figure S 1.**
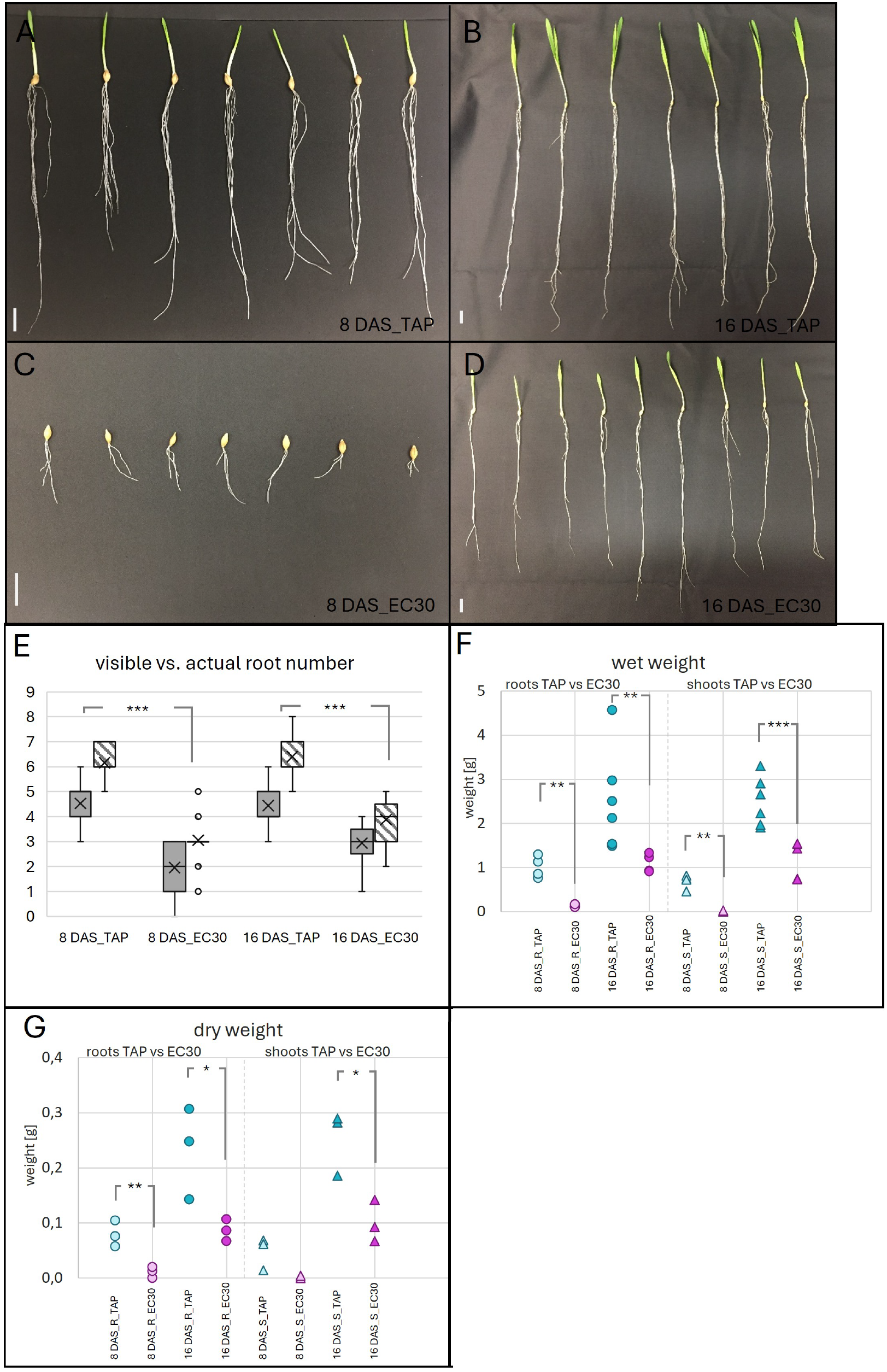
Phenotyping of barley plants under salt stress and control conditions. (A-D) Barley plants harvested after 8 DAS or 16 DAS, after growing in soil, in rhizoboxes, under salt stress (EC30) or control (TAP) conditions. Side bars indicate size differences. (E) Indicates the number of visible roots through the glass front of a rhizobox versus the actual number of roots a barley plant shows, after growing for 8 and 16 days under salt stress and control conditions. n= 19-33 plants. (F) Graph of wet weight (y-axis, in cm) of roots (left part) and shoots (right part) of barley plants grown for 8 DAS and 16 DAS under salt stress and control conditions. n= 3-6, whereas one replicate equals the mean weight (of root/shoot) of 10 plants each. (G) Graph of dry weight (y-axis, in cm) of roots (left part) and shoots (right part) of barley plants grown for 8 and 16 DAS under salt stress and control conditions. n= 3, whereas one replicate equals the mean weight (of root/shoot) of 10 plants each. * Indicating a p-value of ≤ 0.05, ** Indicating a p-value of ≤ 0.01, *** indicating a p-value of ≤ 0.001 upon *t*-tests.

**Figure S 2.**
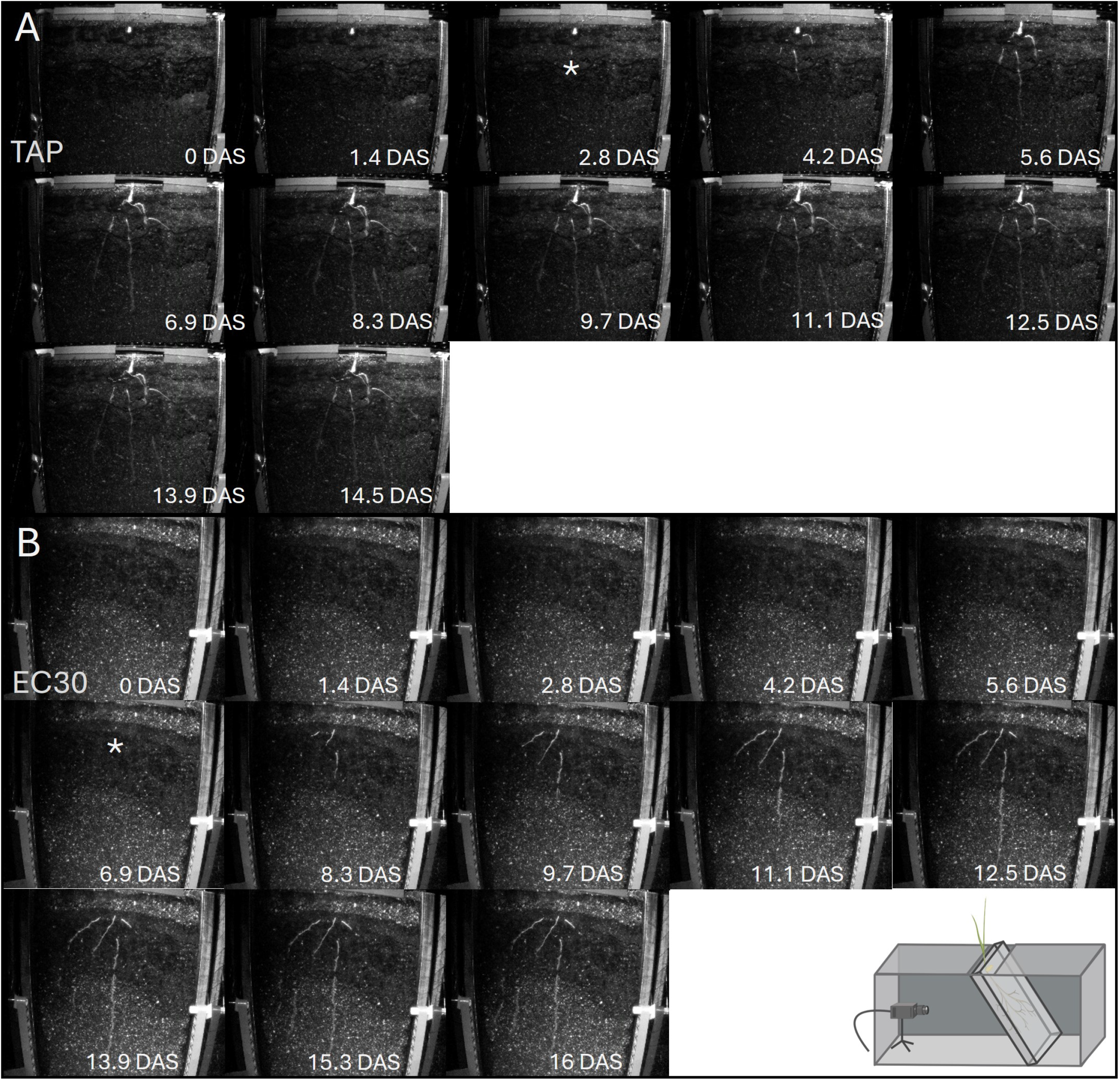
Pictures of growing root through time, 0 to 16 DAS. Pictures taken in DRD-BIBLOXes with camera system and IR light flashes. Roots are growing in soil, in rhizoboxes (A) for 14.5 DAS under control conditions or (B) for 16 DAS under salt stress conditions. The asterixis (*) indicate the respective first appearance of the roots in the recording of root development, for (A) control at 2.8 DAS and for (B) salt stress at 6.9 DAS.

**Figure S 3.**
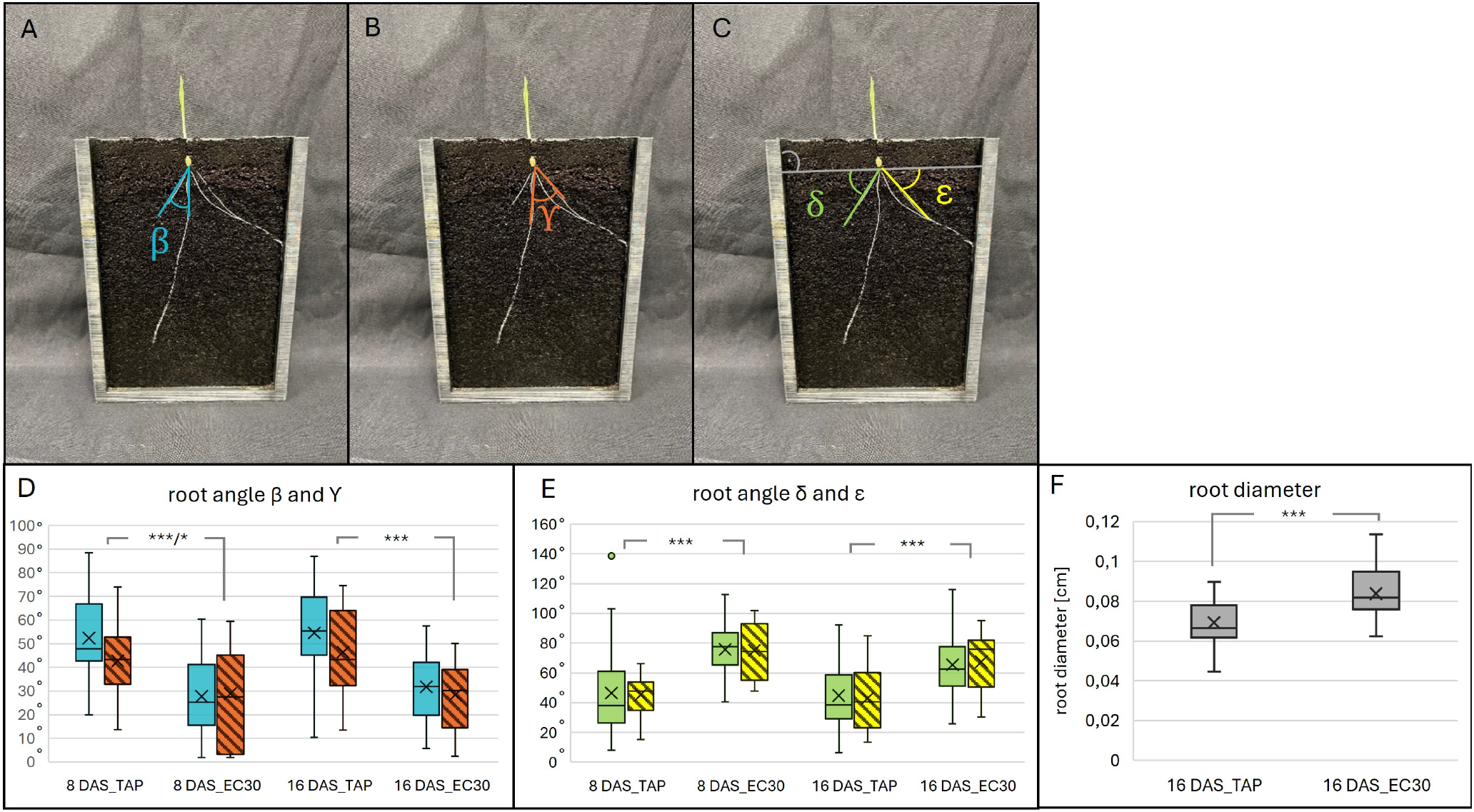
Root angle measurements of barley roots in soil under salt stress. Pictures of a barley plant growing in a rhizobox, labeled to indicate root angle measurement. (A) *β* represents the angle between the (furthest) left seminal root and the primary root. (B) *γ* represents the angle between the (furthest) right seminal root and the primary root. (C) *δ* and *ϵ* represent the angles from the primary root to the respective right and left edge, perpendicular to the rhizobox. (D, E) Graphs of previous described root angles of 8 and 16 DAS plants, grown under salt stress and control conditions. *β* (blue), *γ* (orange striped), *δ* (green), *ϵ* (yellow striped); n=14-33 plants per category. (F) Root diameter measured wtih with ImageJ, from 16 DAS_TAP and 16 DAS_EC30 roots. n = 18-23; *** indicating a p-value of ≤ 0.001 and * indicating a p-value of ≤ 0.05 upon *t*-tests.

**Figure S 4.**
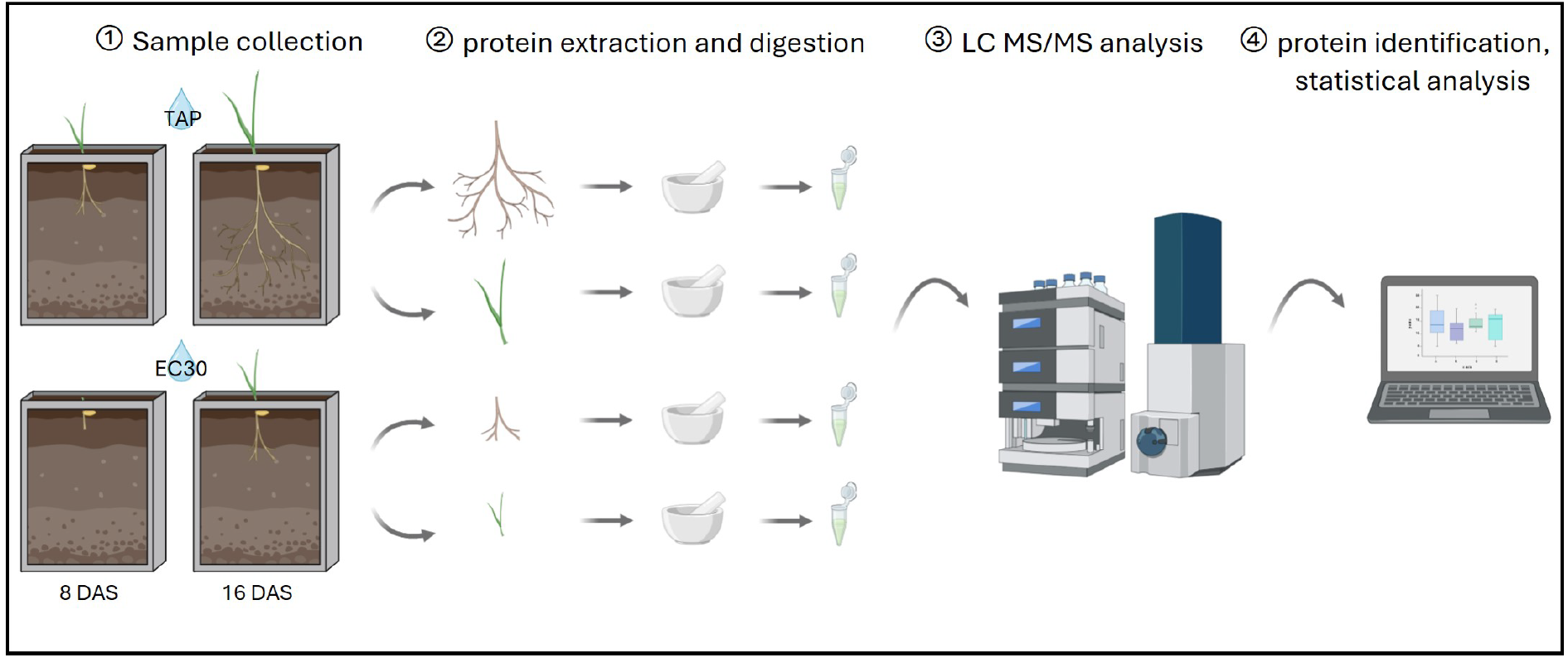
Graphical overview of proteomics experiment. Showing sampling process of plants of 8 DAS and 16 DAS, grown under control (TAP) and salt stress (EC30) conditions. Followed by protein extraction and digestion to peptides and subsequent liquid chromatography and tandem mass spectrometry as well as data analysis.

**Figure S 5.**
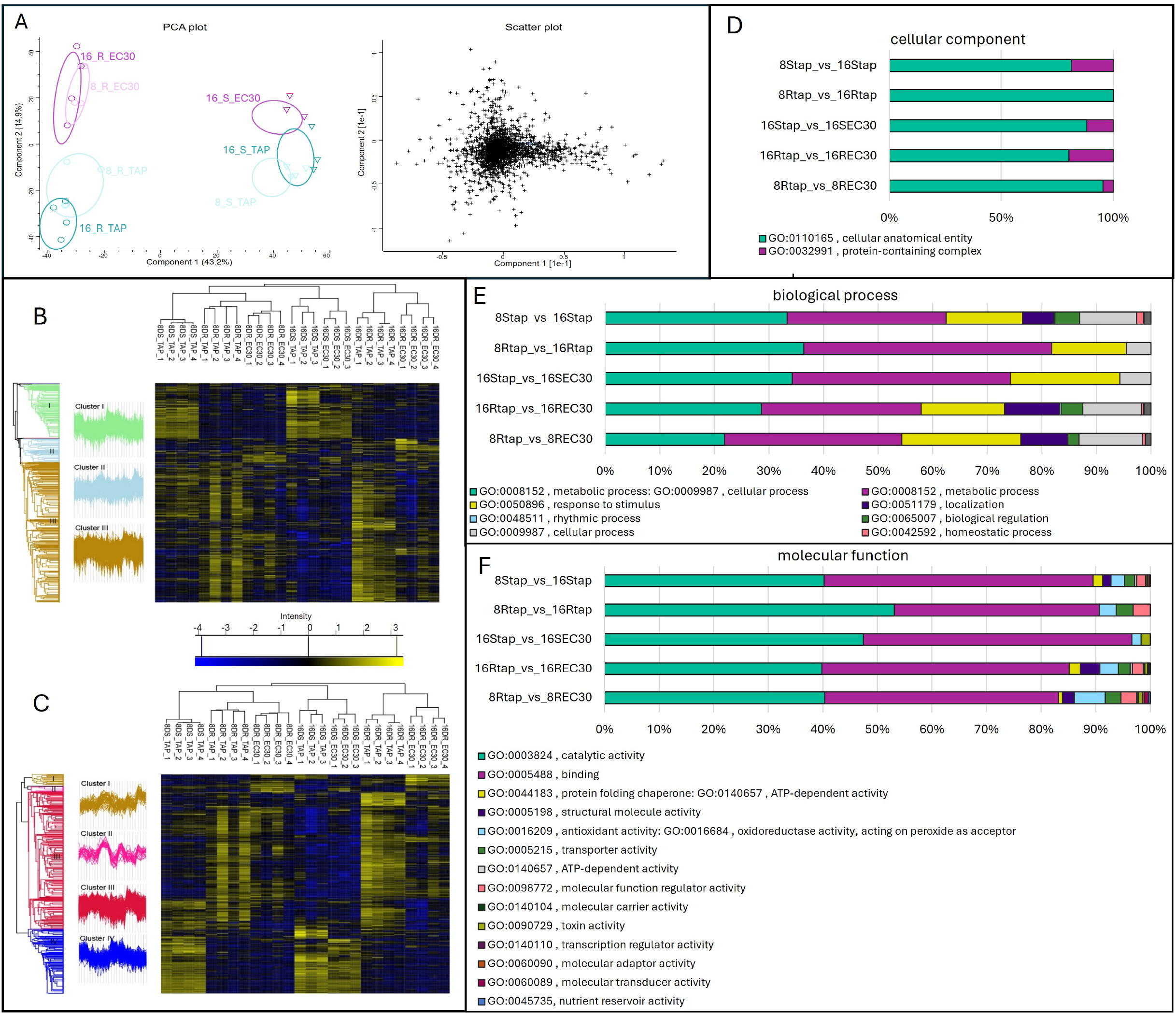
PCA, Scatter plot over all data, Hierarchical cluster analysis, and Gene ontology analysis of significantly up/down regulated proteins. (A) Principal component analysis and scatter plot over all data. Principal component 1 (PC1) separates the proteins regarding root and shoot specificity. PC2 separates the proteins according to control and stressed treatment. Symbols: roots (circle) and shoots (triangle) and control (cyan), salt stress (pink), early development 8DAS (lighter colour), later development 16DAS (darker colour); (B-C) Hierarchical cluster analysis heat maps of (B) the whole data set and (C) only the proteins identified as significant. Z-scores were calculated and are represented by the heat map coloring scheme. Protein clusters trends are included. color code: low abundant protein (blue) to high abundant protein (yellow); (D-F) Protein levels of plant material of root (R) and shoot (S) of 8 DAS (8) and 16 DAS (16) plants under control (TAP) and salt stress (EC30) treatment are analysed. Categorical sample groupings for protein level comparisons were chosen with respect to tissue specific salt stress response and tissue specific development, respectively. Categories for comparison upon *t*-tests were paired as following: 8 DAS roots of TAP treatment versus 16 DAS roots of TAP treatment (8RTAP_vs_16RTAP) and 8 DAS shoots of TAP treatment versus 16 DAS shoots of TAP treatment (8STAP_vs_16STAP) groupings were chosen regarding analysing tissue specific developmental processes. And 8 DAS roots of TAP treatment versus 8 DAS roots of EC30 treatment (8RTAP_vs_8REC30), 16 DAS roots of TAP treatment versus 16 DAS roots of EC30 treatment (16RTAP_vs_16REC30) and 16 DAS shoots of TAP treatment versus 16 DAS shoots of EC30 treatment (16STAP_vs_16SEC30) groupings were chosen to analyse tissue specific salt stress response **(Supplementary file Perseus)**.(D) Analysis of cellular components (GO annotation) —(E) Analysis of biological process (GO annotation) —(F) Analysis of molecular function (GO annotation) of significantly differently regulated proteins when comparing different categories during development or under salt stress.

**Figure S 6.**
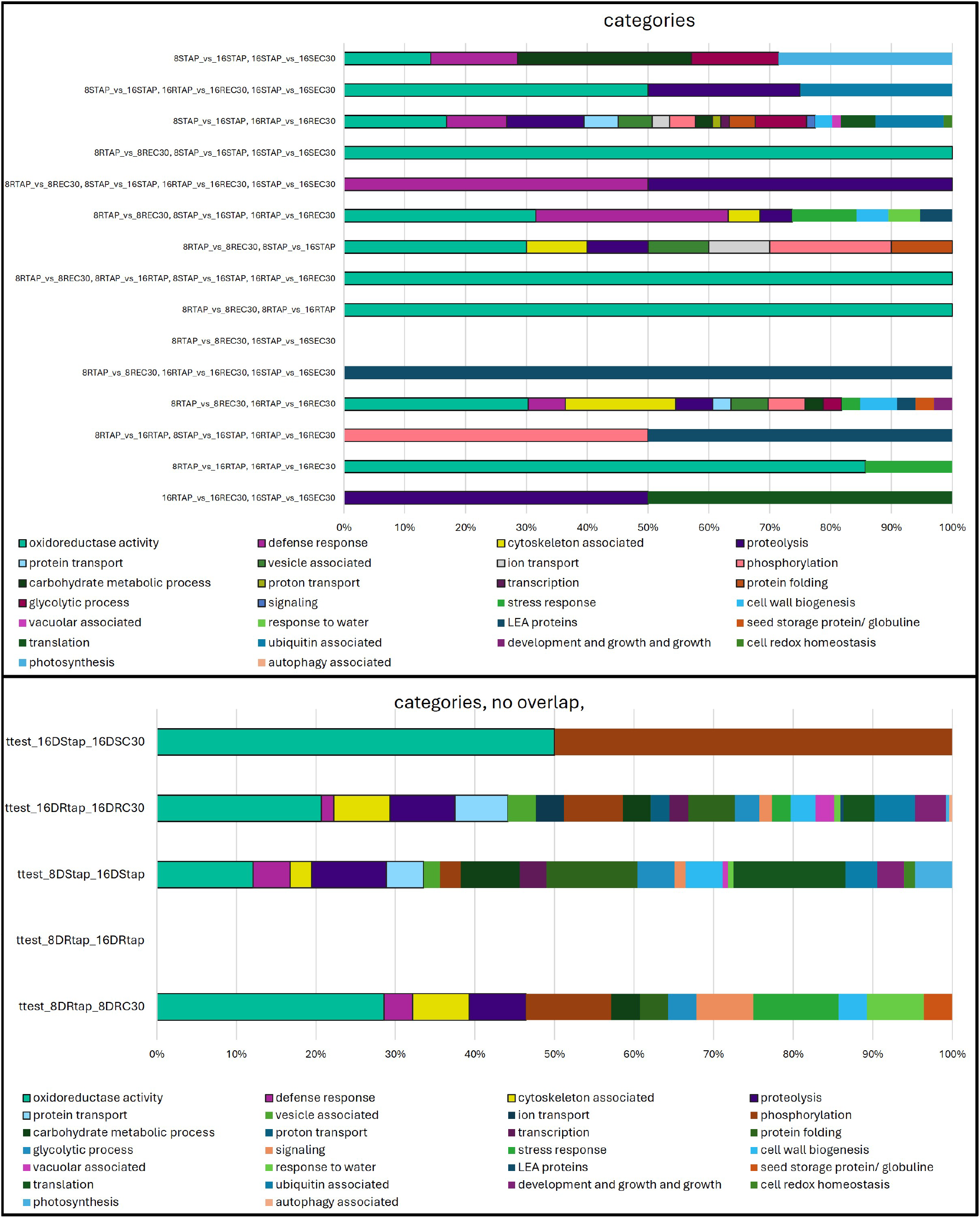
Gene ontology analysis of significantly up/down regulated proteins - close up into Venn Data analysis of Figure2 B. part1. Venn data combinations of significantly differently regulated proteins when comparing different preselected annotation categories during development or under salt stress. (top) Overview of all tested Venn Data points of intersection. (bottom) Overview of all tested Venn Data points without any intersection.

**Figure S 7.**
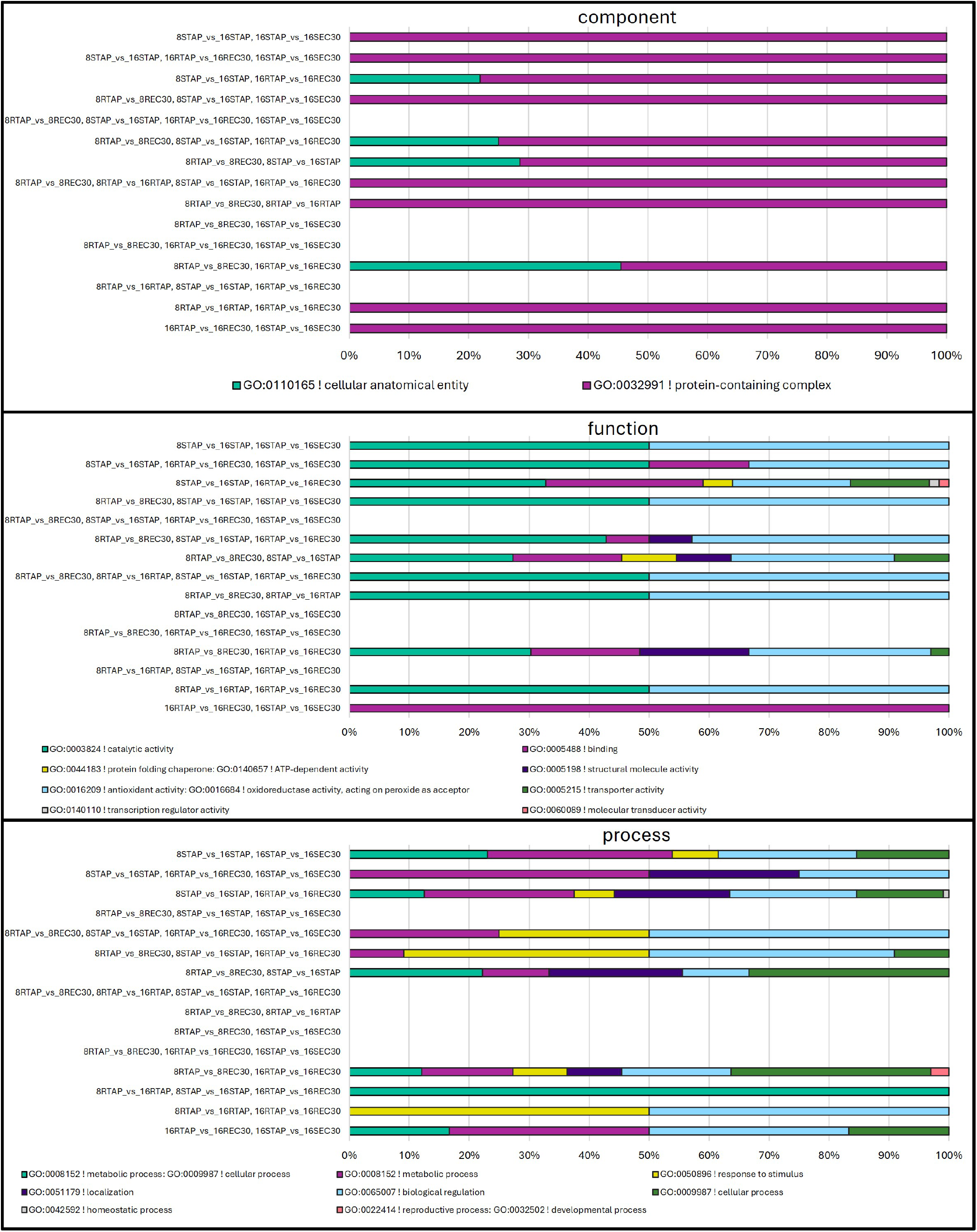
Gene ontology analysis of significantly up/down regulated proteins - close up into Venn Data analysis of Figure2 B. part2. Protein groups of Venn Data points of intersection of significantly differently regulated proteins when comparing GO annotation categories during development or under salt stress. (top) Analysis of cellular components (GO annotation). (middle) Analysis of biological process (GO annotation). (bottom) Analysis of molecular function (GO annotation).

**Figure S 8.**
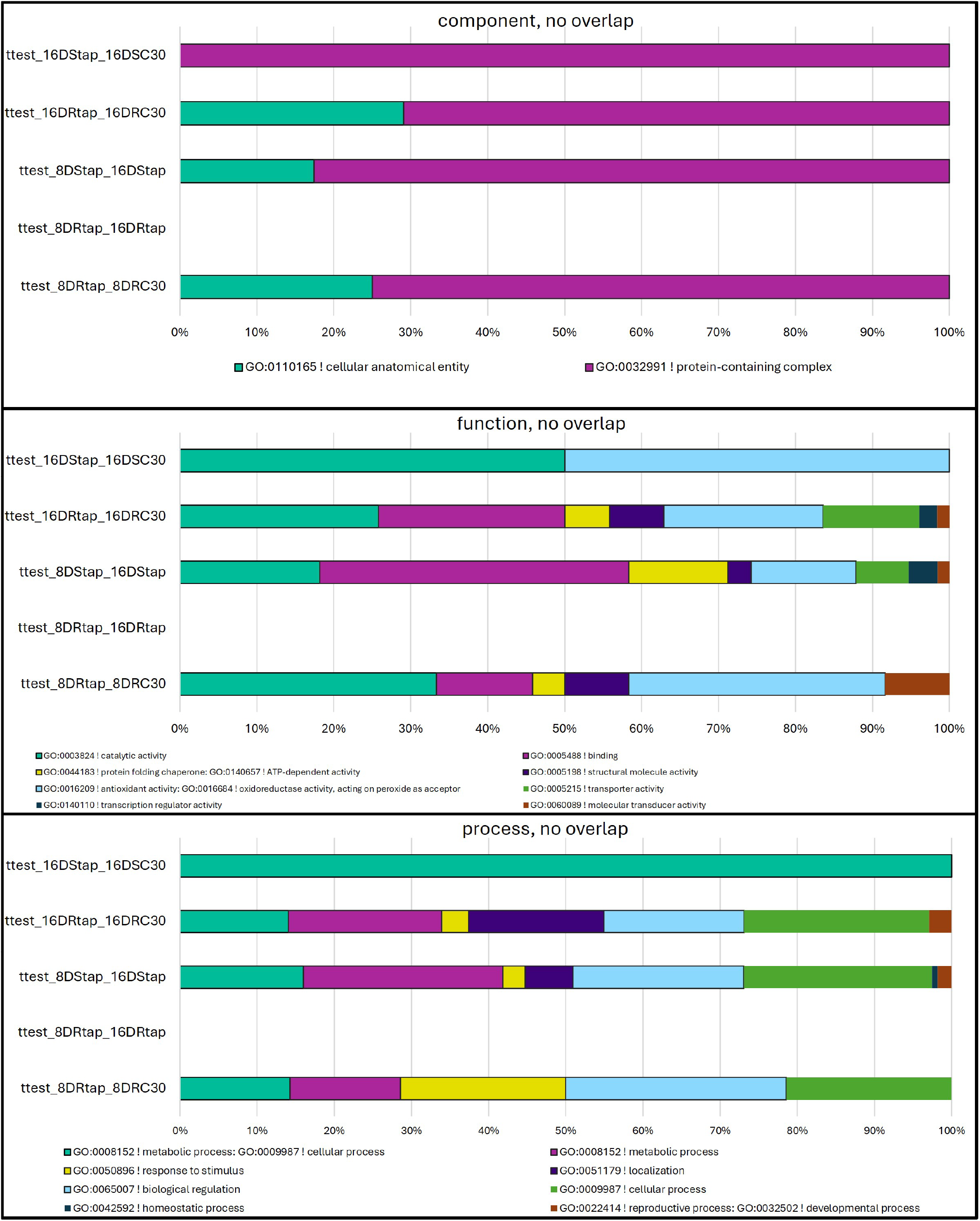
Gene ontology analysis of significantly up/down regulated proteins - close up into Venn Data analysis of Figure2 B. part3. Protein groups of Venn Data points without any intersection of significantly differently regulated proteins when comparing GO annotation categories during development or under salt stress. (top) Analysis of cellular components (GO annotation). (middle) Analysis of biological process (GO annotation). (bottom) Analysis of molecular function (GO annotation).

**Figure S 9.**
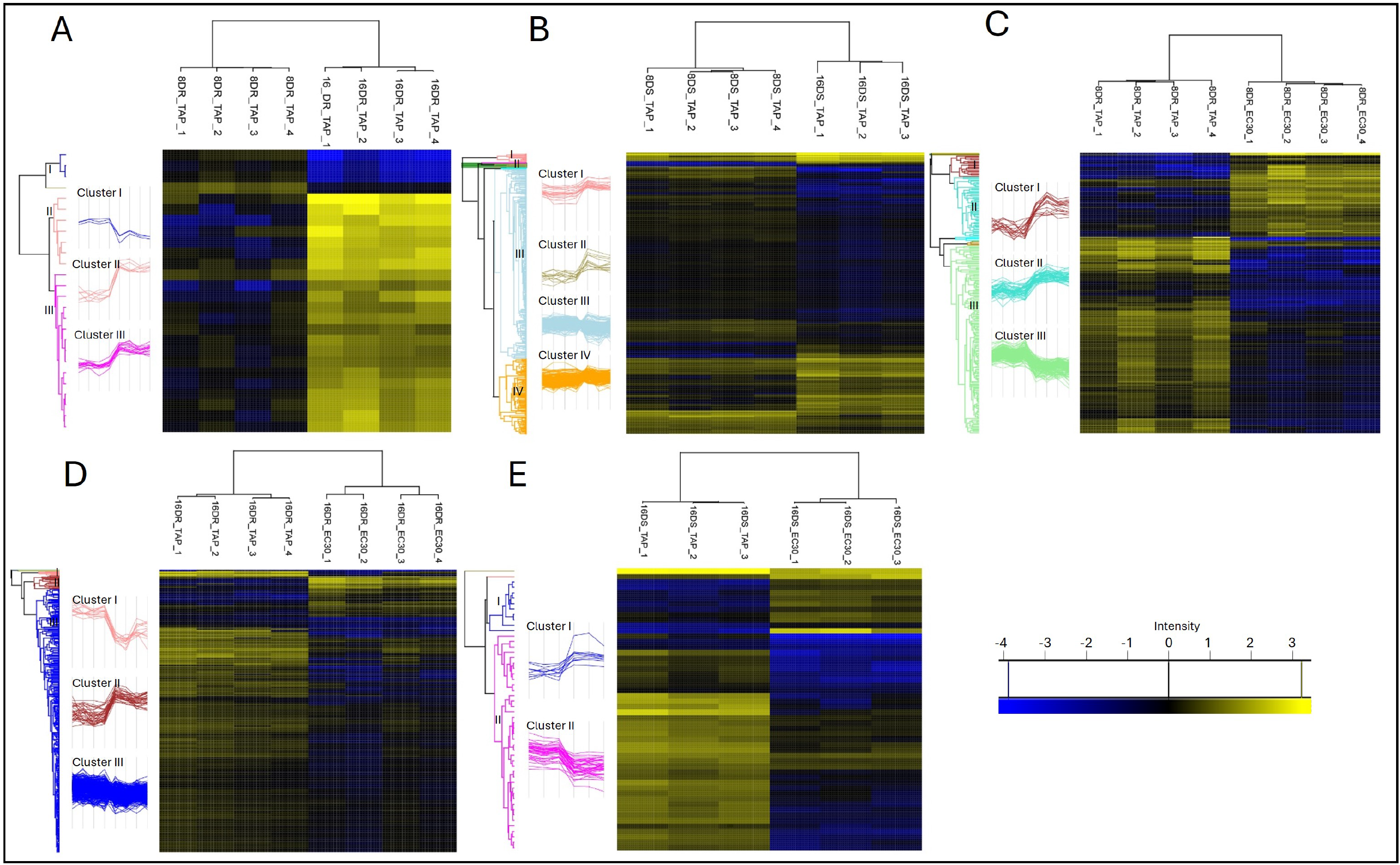
Hierarchical cluster analysis heat maps of significant different abundant proteins upon *t*-tests. Heat maps and protein clusters after comparing (A,B) proteins levels of roots and shoots during development, (C, D) protein levels of roots under salt stress and (E) protein levels of shoots under salt stress. Clusters of similar trending proteins are depicted in color left to heat maps, every line represents protein levels of a single protein. color code: low abundant protein (blue) to high abundant protein (yellow)

**Figure S 10.**
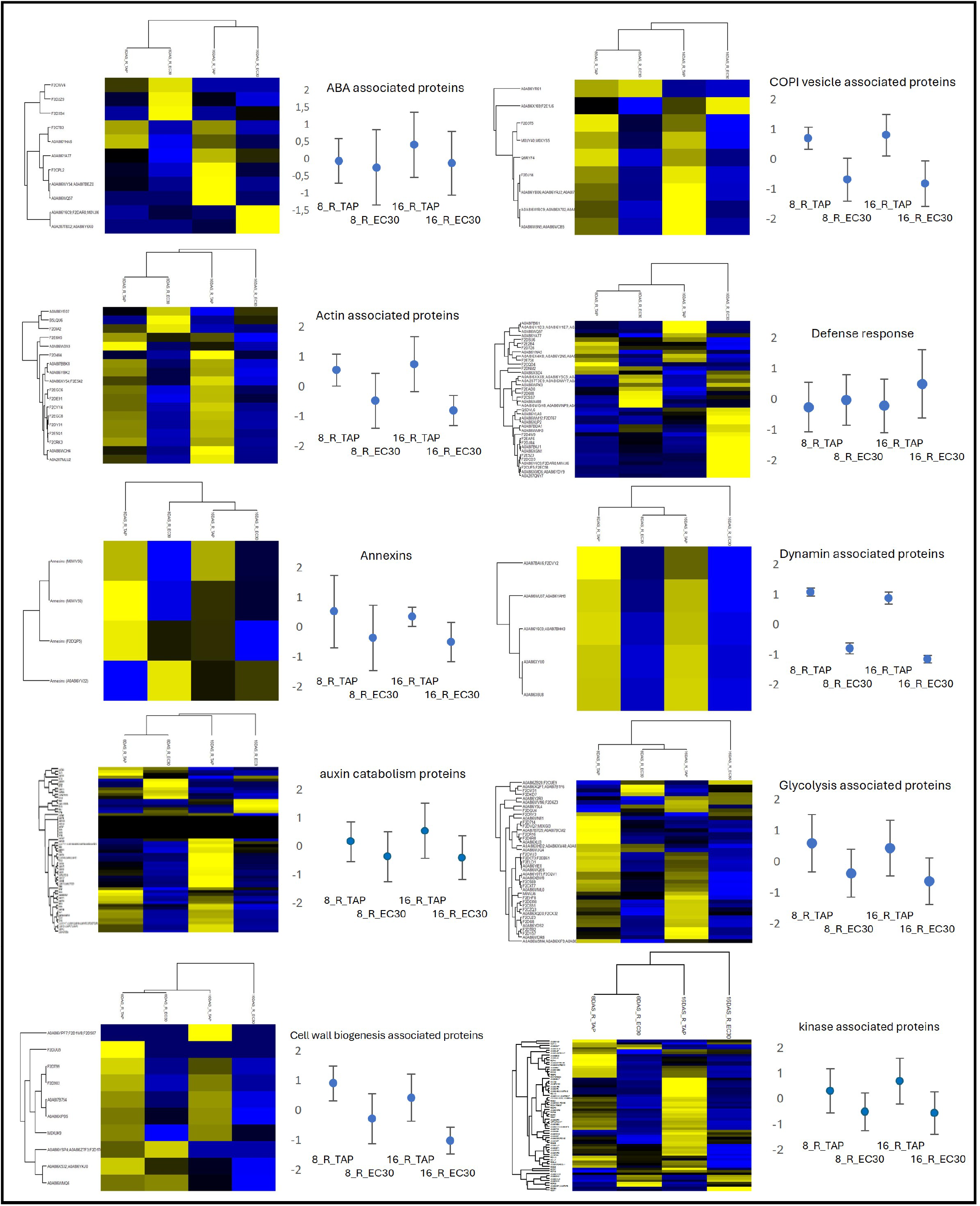
Hierarchical cluster analysis heat maps of z-scores over root data part 1. Means of original LFQ intensity value data were used for z-scores. Z-score results were used for hierarchical cluster analysis according to groups of **Table S 1**. Additionally group specific z-scores are depicted as a graph over root samples.

**Figure S 11.**
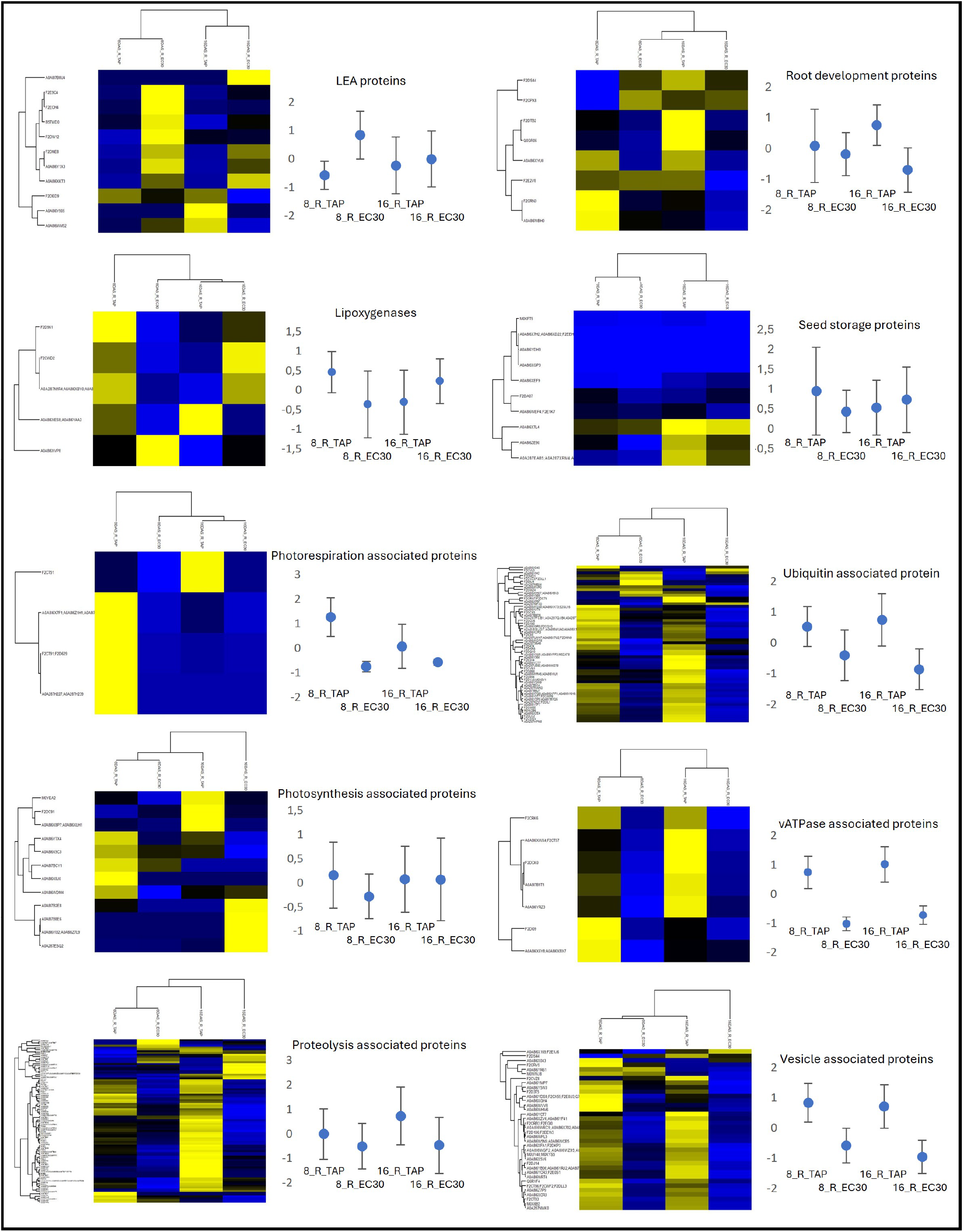
Hierarchical cluster analysis heat maps of z-scores over root data part 2. Means of original LFQ intensity value data were used for z-scores. Z-score results were used for hierarchical cluster analysis according to groups of **Table S 1**. Additionally group specific z-scores are depicted as a graph over root samples.

**Figure S 12.**
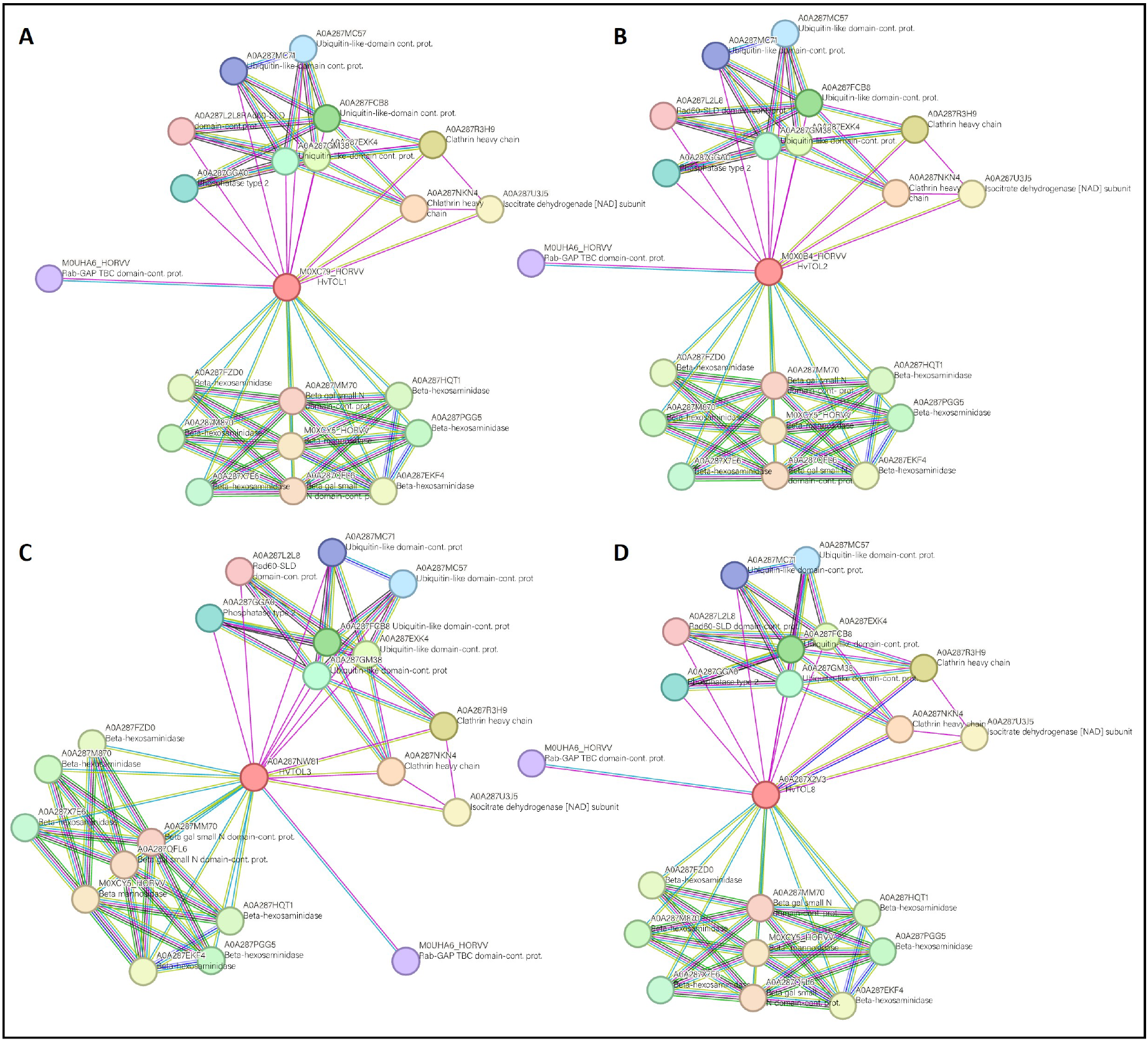
String data first shell interactors of HvTOL1, HvTOL2, HvTOL3, and HvTOL8. Precicted according to STRING database. Detailed description in Table 2.

**Figure S 13.**
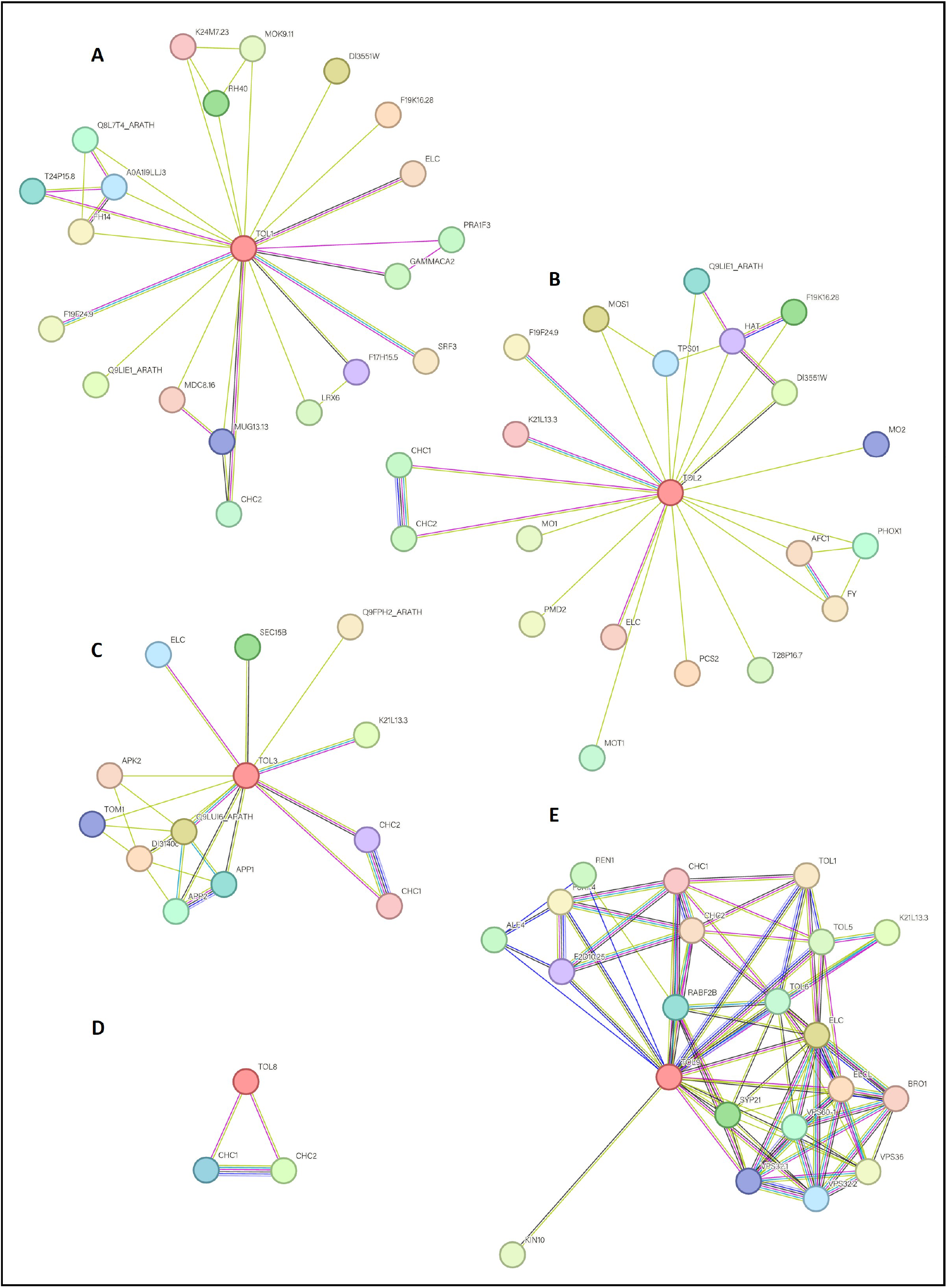
String data first shell interactors of AtTOL1, AtTOL2, AtTOL3, AtTOL8, and AtTOL9. Precicted according to STRING database. Detailed description in Table 3.

**Figure S 14.**
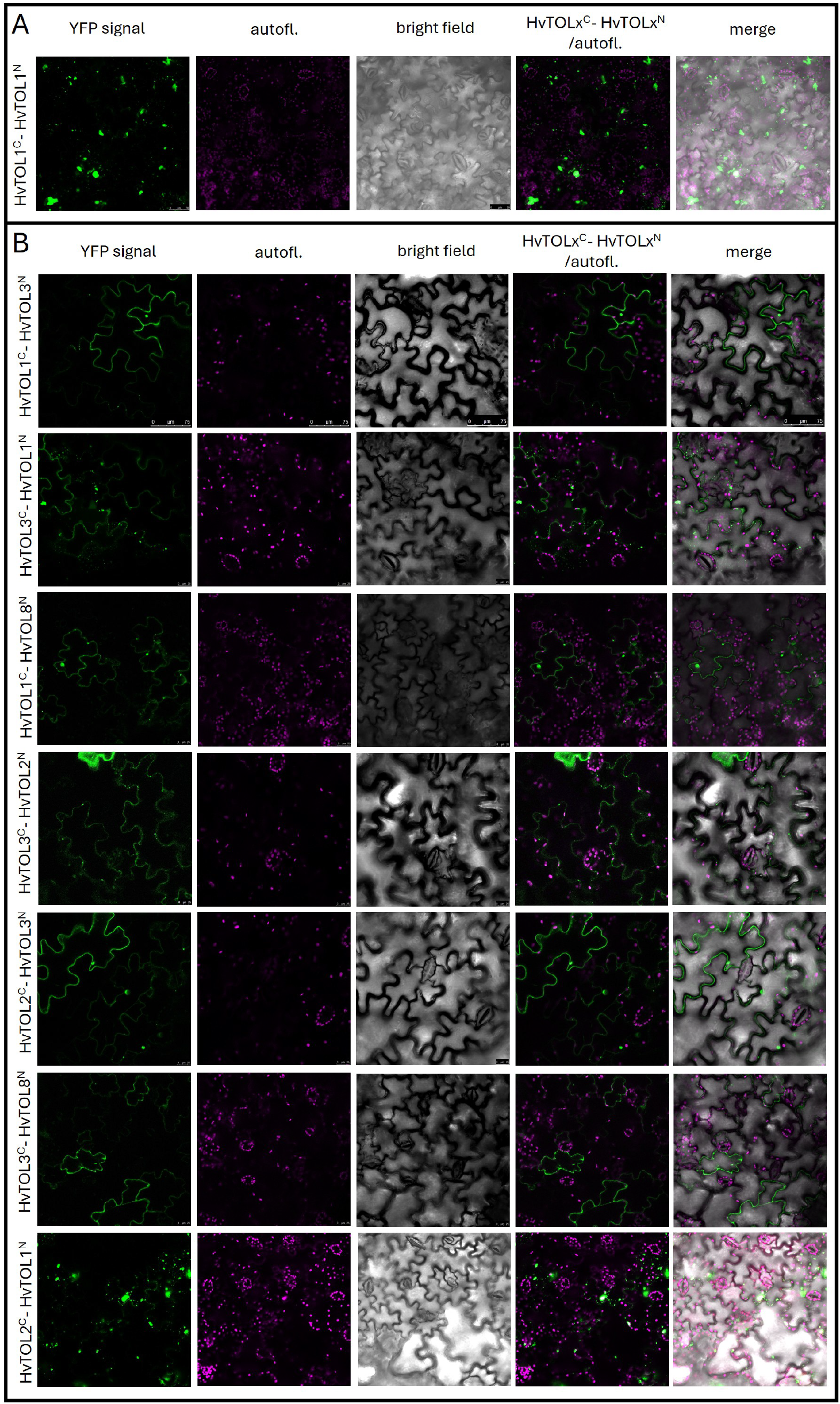
BiFC homodimeres and heterodimeres of HvTOL proteins. Different interaction results for BiFC in tobacco plants for HvTOL 1/2/3/8 upon split YFP combination on N(SPYNE) and C(SPYCE) term. (A) homodimeres,(B) heterodimeres; Pictures show from left to right the YFP signal in green, the autofluorescence in magenta, the bright field picture in grey tones, the merge between YFP and auto-fluorescence signal and the merge of YFP, auto-fluorescence and bright field signal. Negative controls are depicted in **(Figure 4A)**. An Argon laser and emission filter settings of 521nm - 587nm was used for the YFP signals and 608nm - 695nm for auto-fluorescence detection was used.

**Figure S 15.**
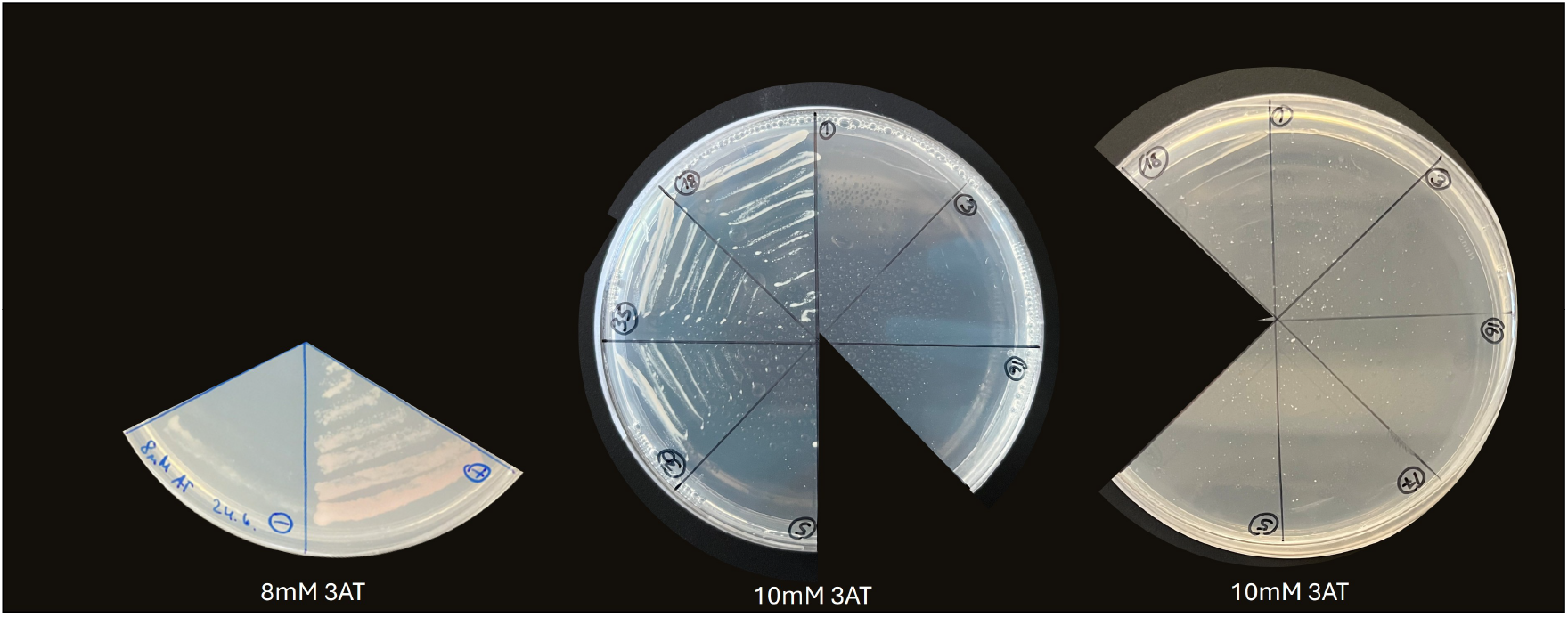
Y2H assay. Y2H results, as depicted in Figure4 B. Reporter gene assay - colony strike outs on SD-Leu-Trp-His + 8mM/10mM 3AT after 48h at 28 °C. (-) pGADT7 + pBTM116 (negative control), (+) pGADT7-AP2C1 + pBTM116-AtMPK6 (positive control), (1) pGADT7 + pBTM117 (negative control), (3) pGADT7-HvTOL3 + pBTM117, (16) pGADT7-HvTOL8 + pBTM117, (17) pGADT7 + pBTM117-HvTOL8, (5) pGADT7-AtMPK6 + pBTM117, (30) pGADT7-HvTOL3 + pBTM117-HvTOL8, (35) pGADT7-HvTOL8 + pBTM117-HvTOL8. Pictures were cropped for clearer visualization.

**Figure S 16.**
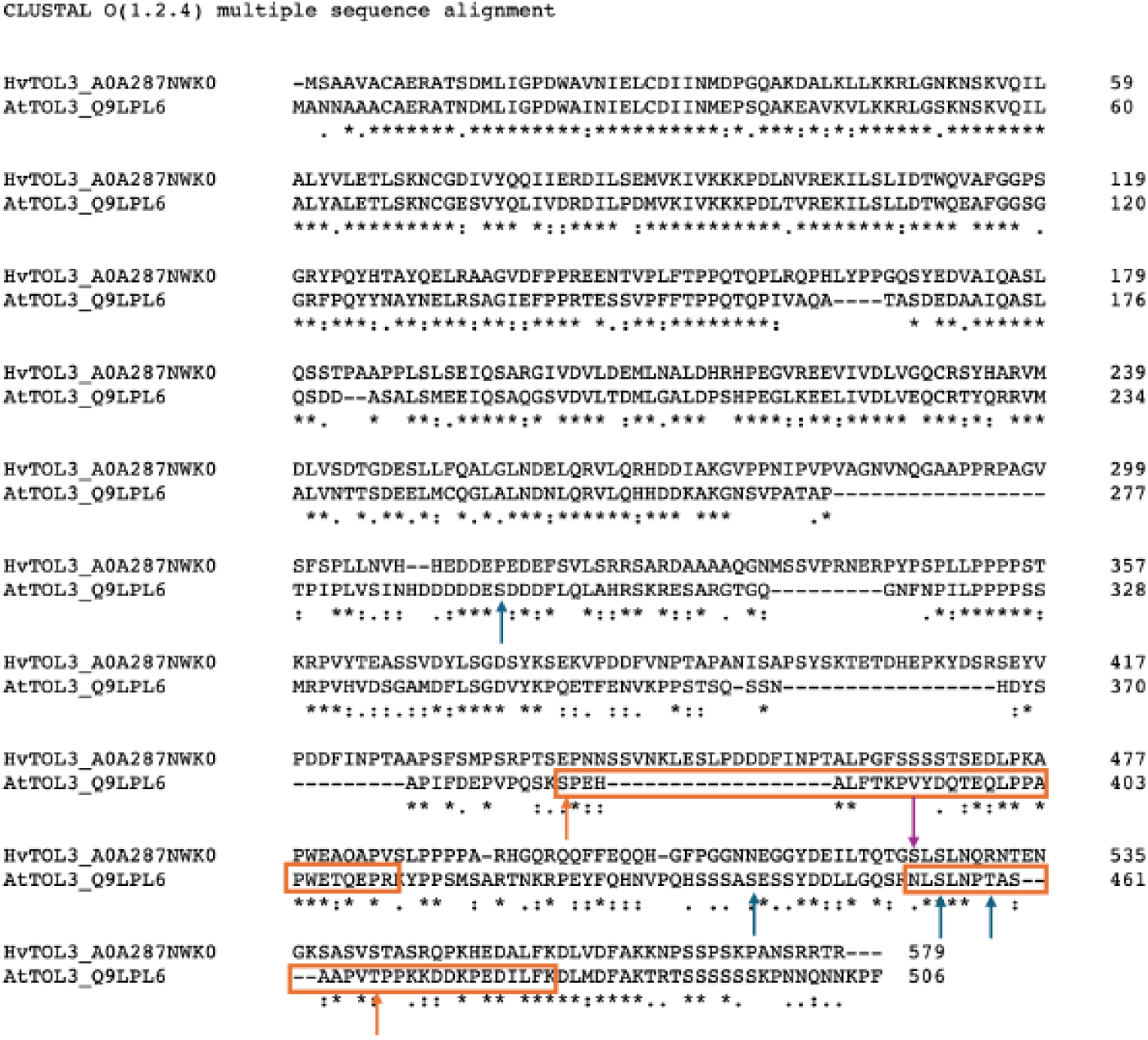
Alignment of HvTOL3 and AtTOL3. Phosphosites are indicated in arrows. Blue according to (https://www.p3db.org/p3dbid_1756277), orange according to (Rayapuram et al., 2018; Hoehenwarter et al., 2013), and magenta according to the homepage plant PTM viewer (https://www.psb.ugent.be/webtools/ptm-viewer/index.php).

**Figure S 17.**
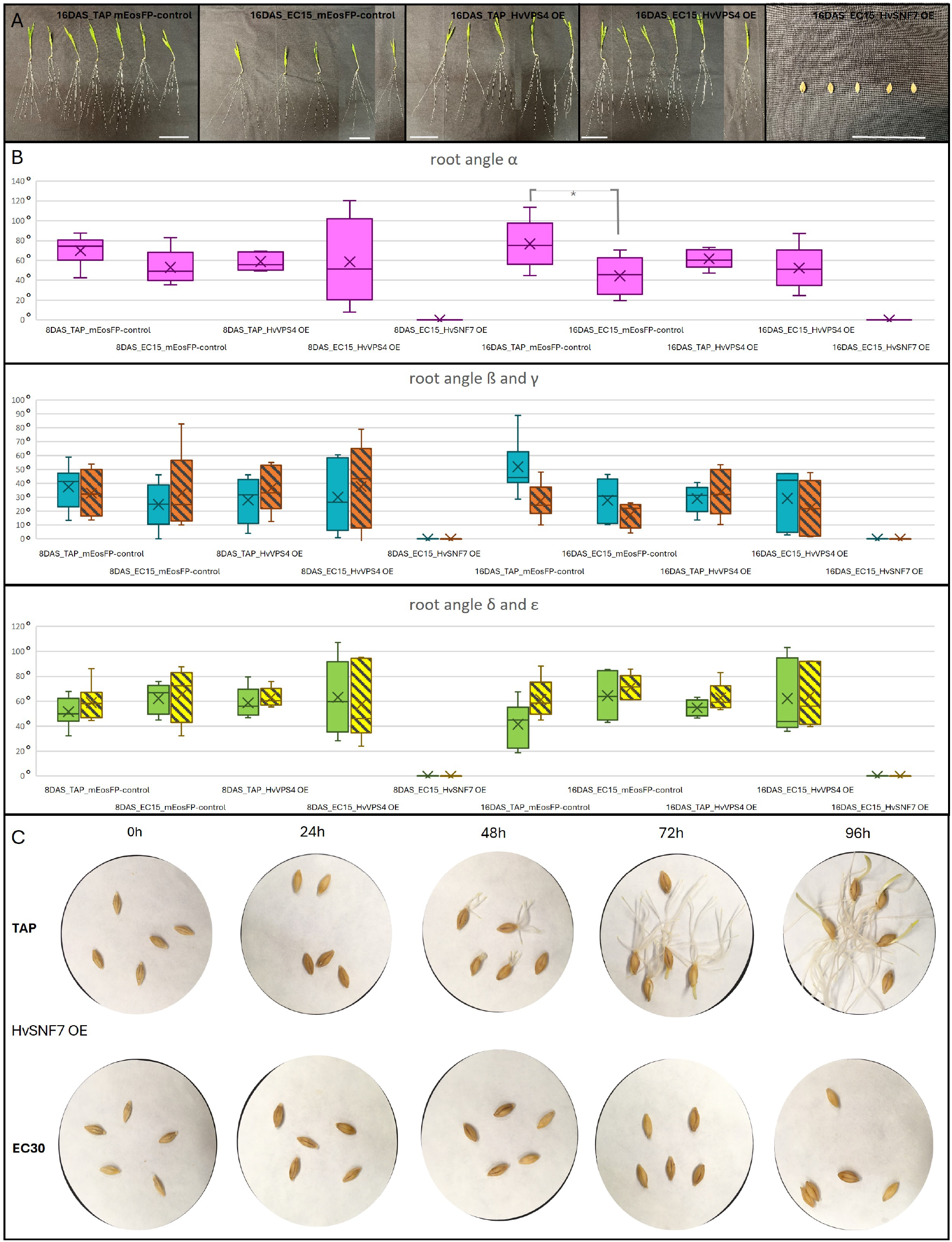
pAct-mEosFP, pAct-SNF7.1-mEosFP, and pAct-mEosFP-HvVPS4 plants grown for 8 DAS and 16 DAS in TAP or EC15 soil - measurements. (A) Plants harvested and cleaned from phenotyping boxes, pAct-mEosFP, pAct-HvSNF7.1-mEosFP, and pAct-mEosFP-HvVPS4, grown for 8 DAS and 16 DAS in TAP or salt stressed (EC15, 170 mM NaCl) soil. scale represents 8 cm.(B) root angle *α*, spanning along the two furthest apart seminal roots. Root angle *β* represents the angle between the (furthest) left seminal root and the primary root, *γ* represents the angle between the (furthest) right seminal root and the primary root. *δ* and *ε* represent the angles from the primary root to the respective right and left edge, perpendicular to the rhizobox. *β* (blue), *γ* (orange striped), *δ* (green), *ϵ* (yellow striped); n=14-33 plants per category; ** indicating a p-value of ≤ 0.01 and * indicating a p-value of ≤ 0.05 upon *t*-tests. n= 5-9 (C) Germination assay with HvSNF7.1-mEosFP over-expression line. Overview of germination of HvSNF7.1-mEosFP barley grains in Petri dishes over 0-96h after imbibition in TAP or EC30 solution.

## Footnotes

1 https://string-db.org/

2 https://bar.utoronto.ca/eplant/

3 https://bar.utoronto.ca

4 https://genecust.org

5 http://chopchop.cbu.uib.no

6 http://crispor.tefor.net

7 http://rna.tbi.univie.ac.at/cgi-bin/RNAWebSuite/RNAfold.cgi

8 https://webblast.ipk-gatersleben.de/barley_ibsc

9 https://ics.hutton.ac.uk/gmapper/gmap_page.html

10 http://n2t.net/addgene:62203

11 https://www.ncbi.nlm.nih.gov/

12 https://www.uniprot.org/

13 https://www.megasoftware.net

## REFERENCES

A. Alrajhi, S. Alharbi, S. Beecham, and F. Alotaibi. Regulation of root growth and elongation in wheat. Front Plant Sci, 15:1397337, 2024. ISSN 1664-462X (Print) 1664-462X (Electronic) 1664-462X (Linking). doi: 10.3389/fpls.2024.1397337. URL https://www.ncbi.nlm.nih.gov/pubmed/38835859.

M. Babst and G. Odorizzi. The balance of protein expression and degradation: an escrts point of view. Curr Opin Cell Biol, 25(4):489– 94, 2013. ISSN 0955-0674 (Print) 0955-0674. doi: 10.1016/j.ceb.2013.05.003.

D. D. Bilgin, J. A. Zavala, J. Zhu, S. J. Clough, D. R. Ort, and E. H. DeLucia. Biotic stress globally downregulates photosynthesis genes. Plant Cell Environ, 33(10):1597–613, 2010. ISSN 1365-3040 (Electronic) 0140-7791 (Linking). doi: 10.1111/j.1365-3040.2010.02167.x. URL https://www.ncbi.nlm.nih.gov/pubmed/20444224.

R. A. Buono, A. Leier, J. Paez-Valencia, J. Pennington, K. Goodman, N. Miller, P. Ahlquist, T. T. Marquez-Lago, and M. S. Otegui. Escrt-mediated vesicle concatenation in plant endosomes. J Cell Biol, 216 (7):2167–2177, 2017. ISSN 0021-9525 (Print) 0021-9525. doi: 10.1083/jcb.201612040.

C. S. Byrt, R. Munns, R. A. Burton, M. Gilliham, and S. Wege. Root cell wall solutions for crop plants in saline soils. Plant Sci, 269:47– 55, 2018. ISSN 1873-2259 (Electronic) 0168-9452 (Linking). doi: 10.1016/j.plantsci.2017.12.012. URL https://www.ncbi.nlm.nih.gov/pubmed/29606216.

Y. Cai, X. Zhuang, C. Gao, X. Wang, and L. Jiang. The arabidopsis endosomal sorting complex required for transport iii regulates internal vesicle formation of the prevacuolar compartment and is required for plant development. Plant Physiol, 165(3):1328–1343, 2014. ISSN 1532-2548 (Electronic) 0032-0889 (Print) 0032-0889 (Linking). doi: 10.1104/pp.114.238378. URL https://www.ncbi.nlm.nih.gov/pubmed/24812106.

J. P. Concordet and M. Haeussler. Crispor: intuitive guide selection for crispr/cas9 genome editing experiments and screens. Nucleic Acids Res, 46(W1):W242–W245, 2018. ISSN 1362-4962 (Electronic) 0305-1048 (Print) 0305-1048 (Linking). doi: 10.1093/nar/gky354. URL https://www.ncbi.nlm.nih.gov/pubmed/29762716.

J. Cox and M. Mann. Maxquant enables high peptide identification rates, individualized p.p.b.-range mass accuracies and proteome-wide protein quantification. Nat Biotechnol, 26(12):1367–72, 2008. ISSN 1546-1696 (Electronic) 1087-0156 (Linking). doi: 10.1038/nbt.1511. URL https://www.ncbi.nlm.nih.gov/pubmed/19029910 https://www.nature.com/articles/nbt.1511.

J. Cox, N. Neuhauser, A. Michalski, R. A. Scheltema, J. V. Olsen, and M. Mann. Andromeda: a peptide search engine integrated into the maxquant environment. J Proteome Res, 10(4):1794– 805, 2011. ISSN 1535-3907 (Electronic) 1535-3893 (Linking). doi: 10.1021/pr101065j. URL https://www.ncbi.nlm.nih.gov/pubmed/21254760 https://pubs.acs.org/doi/pdf/10.1021/pr101065j.

J. Cox, M. Y. Hein, C. A. Luber, I. Paron, N. Nagaraj, and M. Mann. Accurate proteome-wide label-free quantification by delayed normalization and maximal peptide ratio extraction, termed maxlfq. Mol Cell Proteomics, 13(9):2513–26, 2014. ISSN 1535-9484 (Electronic) 1535-9476 (Print) 1535-9476 (Linking). doi: 10.1074/mcp.M113.031591. URL https://www.ncbi.nlm.nih.gov/pubmed/24942700 https://www.mcponline.org/article/S1535-9476(20)33310-7/pdf.

G. Dermendjiev, M. Schnurer, J. Weiszmann, S. Wilfinger, E. Ott, C. Gebert, W. Weckwerth, and V. Ibl. Tissue-specific proteome and subcellular microscopic analyses reveal the effect of high salt concentration on actin cytoskeleton and vacuolization in aleurone cells during early germination of barley. Int J Mol Sci, 22(17), 2021. ISSN 1422-0067 (Electronic) 1422-0067 (Linking). doi: 10.3390/ijms22179642. URL https://www.ncbi.nlm.nih.gov/pubmed/34502558.

G. Dermendjiev, M. Schnurer, E. Stewart, T. Nagele, G. Marino, D. Leister, A. Thur, S. Plott, J. Jez, and V. Ibl. A bench-top dark-root device built with lego((r)) bricks enables a non-invasive plant root development analysis in soil conditions mirroring nature. Front Plant Sci, 14:1166511, 2023. ISSN 1664-462X (Print) 1664-462X (Electronic) 1664-462X (Linking). doi: 10.3389/fpls.2023.1166511. URL https://www.ncbi.nlm.nih.gov/pubmed/37324682 https://www.ncbi.nlm.nih.gov/pmc/articles/PMC10264708/pdf/fpls-14-1166511.pdf.

Muhammad Farooq, Suphia Rafique, Noreen Zahra, Abdul Rehman, and Kadambot H. M. Siddique. wRoot system architecture and salt stress responses in cereal crops. Journal of Agronomy and Crop Science, 210(6):e12776, 2024. ISSN 0931-2250. doi: 10.1111/jac.12776. URL https://onlinelibrary.wiley.com/doi/abs/10.1111/jac.12776.

Wieland Fricke, Gulya Akhiyarova, Wenxue Wei, Erik Alexandersson, Anthony Miller, Per Ola Kjellbom, Andrew Richardson, Tobias Wojciechowski, Lukas Schreiber, Dima Veselov, Guzel Kudoyarova, and Vadim Volkov. The short-term growth response to salt of the developing barley leaf. Journal of Experimental Botany, 57(5):1079–1095, 2006. ISSN 0022-0957. doi: 10.1093/jxb/erj095. URL https://doi.org/10.1093/jxb/erj095.

Carlos S Galvan-Ampudia, Magdalena M Julkowska, Essam Darwish, Jacinto Gandullo, Ruud A Korver, Geraldine Brunoud, Michel A Haring, Teun Munnik, Teva Vernoux, and Christa Testerink. Halotropism is a response of plant roots to avoid a saline environment. Current Biology, 23(20):2044–2050, 2013. ISSN 0960-9822. doi: 10.1016/j.cub.2013.08.042. URL https://doi.org/10.1016/j.cub.2013.08.042.

C. Gao, X. Zhuang, Y. Cui, X. Fu, Y. He, Q. Zhao, Y. Zeng, J. Shen, M. Luo, and L. Jiang. Dual roles of an arabidopsis escrt component free1 in regulating vacuolar protein transport and autophagic degradation. Proc Natl Acad Sci U S A, 112(6):1886– 91, 2015. ISSN 0027-8424 (Print) 0027-8424. doi: 10.1073/pnas.1421271112.

C. Gao, X. Zhuang, J. Shen, and L. Jiang. Plant escrt complexes: Moving beyond endosomal sorting. Trends Plant Sci, 22(11):986– 998, 2017. ISSN 1360-1385. doi: 10.1016/j.tplants.2017.08.003.

J. Garcia de la Garma, N. Fernandez-Garcia, E. Bardisi, B. Pallol, J. S. Asensio-Rubio, R. Bru, and E. Olmos. New insights into plant salt acclimation: the roles of vesicle trafficking and reactive oxygen species signalling in mitochondria and the endomembrane system. New Phytol, 205(1):216–39, 2015. ISSN 0028-646x. doi: 10.1111/nph.12997.

Sebastian Gasparis and Mateusz Przyborowski. An Optimized RNA-Guided Cas9 System for Efficient Simplex and Multiplex Genome Editing in Barley (Hordeum vulgare L.), pages 117–142. Springer US, New York, NY, 2020. ISBN 978-1-0716-0616-2. doi: 10.1007/978-1-0716-0616-2_8. URL https://doi.org/10.1007/978-1-0716-0616-2_8.

Z. Gong, H. Koiwa, M. A. Cushman, A. Ray, D. Bufford, S. Kore-eda, T. K. Matsumoto, J. Zhu, J. C. Cushman, R. A. Bressan, and P. M. Hasegawa. Genes that are uniquely stress regulated in salt overly sensitive (sos) mutants. Plant Physiol, 126(1):363–75, 2001. ISSN 0032-0889 (Print) 0032-0889. doi: 10.1104/pp.126.1.363.

T. J. Haas, M. K. Sliwinski, D. E. Martínez, M. Preuss, K. Ebine, T. Ueda, E. Nielsen, G. Odorizzi, and M. S. Otegui. The arabidopsis aaa atpase skd1 is involved in multivesicular endosome function and interacts with its positive regulator lyst-interacting protein5. Plant Cell, 19(4):1295–312, 2007. ISSN 1040-4651 (Print) 1040-4651. doi: 10.1105/tpc.106.049346.

R. P. Hellens, E. A. Edwards, N. R. Leyland, S. Bean, and P. M. Mullineaux. pgreen: a versatile and flexible binary ti vector for agrobacterium-mediated plant transformation. Plant Mol Biol, 42(6):819–32, 2000. ISSN 0167-4412 (Print) 0167-4412. doi: 10.1023/a:1006496308160.

R. P. Hellens, A. C. Allan, E. N. Friel, K. Bolitho, K. Grafton, M. D. Templeton, S. Karunairetnam, A. P. Gleave, and W. A. Laing. Transient expression vectors for functional genomics, quantification of promoter activity and rna silencing in plants. Plant Methods, 1:13, 2005. ISSN 1746-4811 (Electronic) 1746-4811 (Linking). doi: 10.1186/1746-4811-1-13. URL https://www.ncbi.nlm.nih.gov/pubmed/16359558.

W. M. Henne, N. J. Buchkovich, and S. D. Emr. The escrt pathway. Dev Cell, 21(1):77–91, 2011. ISSN 1878-1551 (Electronic) 1534-5807 (Linking). doi: 10.1016/j.devcel.2011.05.015. URL https://www.ncbi.nlm.nih.gov/pubmed/21763610.

J. Hilscher, E. Kapusi, E. Stoger, and V. Ibl. Cell layer-specific distribution of transiently expressed barley escrt-iii component hvvps60 in developing barley endosperm. Protoplasma, 253(1):137–53, 2016. ISSN 1615-6102 (Electronic) 0033-183X (Print) 0033-183X (Linking). doi: 10.1007/s00709-015-0798-1. URL https://www.ncbi.nlm.nih.gov/pubmed/25796522.

A. Hinchliffe and W. A. Harwood. Agrobacterium-mediated transformation of barley immature embryos. Methods Mol Biol, 1900:115–126, 2019. ISSN 1940-6029 (Electronic) 1064-3745 (Linking). doi: 10.1007/978-1-4939-8944-7_8. URL https://www.ncbi.nlm.nih.gov/pubmed/30460562 https://link.springer.com/protocol/10.1007/978-1-4939-8944-7_8.

Li-Wei Ho, Ting-Ting Yang, Shyan-Shu Shieh, Gerald E. Edwards, and Hungchen E. Yen. Reduced expression of a vesicle trafficking-related atpase skd1 decreases salt tolerance in arabidopsis. Functional Plant Biology, 37(10):962–973, 2010. doi: 10.1071/FP10049. URL https://www.publish.csiro.au/paper/FP10049.

W. Hoehenwarter, M. Thomas, E. Nukarinen, V. Egelhofer, H. Röhrig, W. Weckwerth, U. Conrath, and G. J. Beckers. Identification of novel in vivo map kinase substrates in arabidopsis thaliana through use of tandem metal oxide affinity chromatography. Mol Cell Proteomics, 12 (2):369–80, 2013. ISSN 1535-9476 (Print) 1535-9476. doi: 10.1074/mcp.M112.020560.

Y. W. Hsu and G. Y. Jauh. Vps36-mediated plasma membrane protein turnover is critical for arabidopsis root gravitropism. Plant Signal Behav, 12(4):e1307495, 2017. ISSN 1559-2316 (Print) 1559-2316. doi: 10.1080/15592324.2017.1307495.

Verena Ibl, Edina Csaszar, Nicole Schlager, Susanne Neubert, Christoph Spitzer, and Marie-Theres Hauser. Interactome of the plant-specific escrt-iii component atvps2.2 in arabidopsis thaliana. Journal of Proteome Research, 11(1):397–411, 2012. ISSN 1535-3893. doi: 10.1021/pr200845n. URL https://doi.org/10.1021/pr200845n.

E. Isono. Escrt is a great sealer: Non-endosomal function of the escrt machinery in membrane repair and autophagy. Plant Cell Physiol, 62(5):766–774, 2021. ISSN 1471-9053 (Electronic) 0032-0781 (Linking). doi: 10.1093/pcp/pcab045. URL https://www.ncbi.nlm.nih.gov/pubmed/33768242.

Y. Jou, C. P. Chiang, and H. E. Yen. Changes in cellular distribution regulate skd1 atpase activity in response to a sudden increase in environmental salinity in halophyte ice plant. Plant Signal Behav, 8(12):e27433, 2013. ISSN 1559-2316 (Print) 1559-2316. doi: 10.4161/psb.27433.

E. Kapusi, G. Hensel, Maria Coronado, Sylvia Broeders, Cornelia Marthe, I. Otto, and J. Kumlehn. Elimination of selectable marker genes via segregation of uncoupled t-dnas in populations of doubled haploid barley. Journal für Verbraucherschutz und Lebensmittelsicherheit, 2, 12 2007. doi: 10.1007/s00003-007-0277-5.

Muhammad Owais Khan, Muhammad Irfan, Asim Muhammad, Izhar Ullah, Sultan Nawaz, Mussaddiq Khan Khalil, and Manzoor Ahmad. A practical and economical strategy to mitigate salinity stress through seed priming. Frontiers in Environmental Science, 10, 2022. ISSN 2296-665X. doi: 10.3389/fenvs.2022.991977.

B. Korbei, J. Moulinier-Anzola, L. De-Araujo, D. Lucyshyn, K. Retzer, M. A. Khan, and C. Luschnig. Arabidopsis tol proteins act as gatekeepers for vacuolar sorting of pin2 plasma membrane protein. Curr Biol, 23(24):2500–5, 2013. ISSN 1879-0445 (Electronic) 0960-9822 (Linking). doi: 10.1016/j.cub.2013.10.036. URL https://www.ncbi.nlm.nih.gov/pubmed/24316203.

Pragati Kumari, Arvind Gupta, Harish Chandra, Pratibha Singh, and Saurabh Yadav. Effects of Salt Stress on the Morphology, Anatomy, and Gene Expression of Crop Plants, book section 6, pages 87–105. John Wiley & Sons, Ltd, 2021. ISBN 9781119700517. doi: 10.1002/9781119700517.ch6. URL https://onlinelibrary.wiley.com/doi/abs/10.1002/9781119700517.ch6.

Labun, T. G. Montague, M. Krause, Y. N. Torres Cleuren, H. Tjeldnes, and E. Valen. Chopchop v3: expanding the crispr web toolbox beyond genome editing. Nucleic Acids Res, 47(W1):W171–W174, 2019. ISSN 1362-4962 (Electronic) 0305-1048 (Print) 0305-1048 (Linking). doi: 10.1093/nar/gkz365. URL https://www.ncbi.nlm.nih.gov/pubmed/31106371.

Bingying Leng, Fei Geng, Xinxiu Dong, Fang Yuan, and Baoshan Wang. Sodium is the critical factor leading to the positive halotropism of the halophyte limonium bicolor. Plant Biosystems - An International Journal Dealing with all Aspects of Plant Biology, 153(4):544–551, 2019. ISSN 1126-3504. doi: 10.1080/11263504.2018.1508085. URL https://doi.org/10.1080/11263504.2018.1508085.

P. Li, X. Yang, H. Wang, T. Pan, Y. Wang, Y. Xu, C. Xu, and Z. Yang. Genetic control of root plasticity in response to salt stress in maize. Theor Appl Genet, 134(5):1475–1492, 2021. ISSN 1432-2242 (Electronic) 0040-5752 (Linking). doi: 10.1007/s00122-021-03784-4. URL https://www.ncbi.nlm.nih.gov/pubmed/33661350 https://link.springer.com/article/10.1007/s00122-021-03784-4.

C. Liu, X. Lin, M. Xu, Z. Zheng, Z. Wang, X. Huang, F. Wu, G. Liu, W. Liu, C. Peng, Y. Guo, Y. Zheng, C. Gao, W. Shen, and H. Li. The escrt component fyve4 modulates salt stress response by strengthening the sos1-sos2 interaction in arabidopsis. Plant Commun, page 101428, 2025. ISSN 2590-3462. doi: 10.1016/j.xplc.2025.101428.

W. Liu, R. J. Li, T. T. Han, W. Cai, Z. W. Fu, and Y. T. Lu. Salt stress reduces root meristem size by nitric oxide-mediated modulation of auxin accumulation and signaling in arabidopsis. Plant Physiol, 168 (1):343–56, 2015. ISSN 1532-2548 (Electronic) 0032-0889 (Print) 0032-0889 (Linking). doi: 10.1104/pp.15.00030. URL https://www.ncbi.nlm.nih.gov/pubmed/25818700.

L. Lou, F. Yu, M. Tian, G. Liu, Y. Wu, Y. Wu, R. Xia, J. M. Pardo, Y. Guo, and Q. Xie. Escrt-i component vps23a sustains salt tolerance by strengthening the sos module in arabidopsis. Mol Plant, 13(8):1134–1148, 2020. ISSN 1752-9867 (Electronic) 1674-2052 (Linking). doi: 10.1016/j.molp.2020.05.010. URL https://www.ncbi.nlm.nih.gov/pubmed/32439321.

Liming Luo, Pingping Zhang, Ruihai Zhu, Jing Fu, Jing Su, Jing Zheng, Ziyue Wang, Dan Wang, and Qingqiu Gong. Autophagy is rapidly induced by salt stress and is required for salt tolerance in arabidopsis. Frontiers in Plant Science, Volume 8 - 2017, 2017. ISSN 1664-462X. doi: 10.3389/fpls.2017.01459. URL https://www.frontiersin.org/journals/plant-science/articles/10.3389/fpls.2017.01459.

Ma, X. Liu, W. Lv, and Y. Yang. Molecular mechanisms of plant responses to salt stress. Front Plant Sci, 13:934877, 2022. ISSN 1664-462X (Print) 1664-462x. doi: 10.3389/fpls.2022.934877.

Luke McCormack, Ian A. Dickie, David M. Eissenstat, Timothy J. Fahey, Christopher W. Fernandez, Dali Guo, Heljä-Sisko Helmisaari, Erik A. Hobbie, Colleen M. Iversen, Robert B. Jackson, Jaana Leppälammi-Kujansuu, Richard J. Norby, Richard P. Phillips, Kurt S. Pregitzer, Seth G. Pritchard, Boris Rewald, and Marcin Zadworny. Redefining fine roots improves understanding of below-ground contributions to terrestrial biosphere processes. New Phytologist, 207(3):505–518, 2015. ISSN 0028-646X. doi: 10.1111/nph.13363. URL https://nph.onlinelibrary.wiley.com/doi/abs/10.1111/nph.13363.

Niccolò Mosesso Niharika Savant Lerner, Tobias Bläske, Felix Groh, Shane Maguire, Marie Laura Niedermeier, Eliane Landwehr, Karin Vogel, Konstanze Meergans, Marie-Kristin Nagel, Malte Drescher, Florian Stengel, Karin Hauser, and Erika Isono. Arabidopsis calb1 undergoes phase separation with the escrt protein alix and modulates autophagosome maturation. Nature Communications, 15(1):5188, 2024. ISSN 2041-1723. doi: 10.1038/s41467-024-49485-6. URL https://doi.org/10.1038/s41467-024-49485-6.

A. Mostek, A. Borner, A. Badowiec, and S. Weidner. Alterations in root proteome of salt-sensitive and tolerant barley lines under salt stress conditions. J Plant Physiol, 174:166–76, 2015. ISSN 1618-1328 (Electronic) 0176-1617 (Linking). doi: 10.1016/j.jplph.2014.08.020. URL https://www.ncbi.nlm.nih.gov/pubmed/25462980.

J. Moulinier-Anzola, L. De-Araujo, and B. Korbei. Expression of arabidopsis tol genes. Plant Signal Behav, 9(4):e28667, 2014. ISSN 1559-2316 (Print) 1559-2316. doi: 10.4161/psb.28667.

J. Moulinier-Anzola, M. Schwihla, L. De-Araujo, C. Artner, L. Jorg, N. Konstantinova, C. Luschnig, and B. Korbei. Tols function as ubiquitin receptors in the early steps of the escrt pathway in higher plants. Mol Plant, 13(5):717–731, 2020. ISSN 1752-9867 (Electronic) 1674-2052 (Linking). doi: 10.1016/j.molp.2020.02.012. URL https://www.ncbi.nlm.nih.gov/pubmed/32087370.

J. Moulinier-Anzola, M. Schwihla, R. Lugsteiner, N. Leibrock, M. I. Feraru, I. Tkachenko, C. Luschnig, E. Arcalis, E. Feraru, J. Lozano-Juste, and B. Korbei. Modulation of abscisic acid signaling via endosomal tol proteins. New Phytol, 243(3):1065–1081, 2024. ISSN 1469-8137 (Electronic) 0028-646X (Linking). doi: 10.1111/nph.19904. URL https://www.ncbi.nlm.nih.gov/pubmed/38874374 https://nph.onlinelibrary.wiley.com/doi/pdfdirect/10.1111/nph.19904?download=true.

R. Munns and M. Tester. Mechanisms of salinity tolerance. Annu Rev Plant Biol, 59:651–81, 2008. ISSN 1543-5008 (Print) 1543-5008 (Linking). doi: 10.1146/annurev.arplant.59.032607.092911. URL https://www.ncbi.nlm.nih.gov/pubmed/18444910.

Rana Munns and Matthew Gilliham. Salinity tolerance of crops – what is the cost? New Phytologist, 208(3):668–673, 2015. ISSN 0028-646X. doi: 10.1111/nph.13519. URL https://nph.onlinelibrary.wiley.com/doi/abs/10.1111/nph.13519.

M. S. Otegui and C. Spitzer. Endosomal functions in plants. Traffic, 9(10):1589–98, 2008. ISSN 1600-0854 (Electronic) 1398-9219 (Linking). doi: 10.1111/j.1600-0854.2008.00787.x. URL https://www.ncbi.nlm.nih.gov/pubmed/18627577.

N. Rayapuram, J. Bigeard, H. Alhoraibi, L. Bonhomme, A. M. Hesse, J. Vinh, H. Hirt, and D. Pflieger. Quantitative phosphoproteomic analysis reveals shared and specific targets of arabidopsis mitogen-activated protein kinases (mapks) mpk3, mpk4, and mpk6. Mol Cell Proteomics, 17(1):61–80, 2018. ISSN 1535-9476 (Print) 1535-9476. doi: 10.1074/mcp.RA117.000135.

V. Roustan, J. Hilscher, M. Weidinger, S. Reipert, A. Shabrangy, C. Gebert, B. Dietrich, G. Dermendjiev, M. Schnurer, P. J. Roustan, E. Stoger, and V. Ibl. Protein sorting into protein bodies during barley endosperm development is putatively regulated by cytoskeleton members, mvbs and the hvsnf7s. Sci Rep, 10(1):1864, 2020. ISSN 2045-2322 (Electronic) 2045-2322 (Linking). doi: 10.1038/s41598-020-58740-x. URL https://www.ncbi.nlm.nih.gov/pubmed/32024857.

Miguel Sampaio, João Neves, Tatiana Cardoso, José Pissarra Susana Pereira, and Cláudia Pereira. Coping with abiotic stress in plants— an endomembrane trafficking perspective. Plants, 11(3):338, 2022. ISSN 2223-7747. URL https://www.mdpi.com/2223-7747/11/3/338.

M. Sauer and J. Friml. Plant biology: gatekeepers of the road to protein perdition. Curr Biol, 24(1):R27–R29, 2014. ISSN 1879-0445 (Electronic) 0960-9822 (Linking). doi: 10.1016/j.cub.2013.11.019. URL https://www.ncbi.nlm.nih.gov/pubmed/24405674.

Alois Schweighofer, Vaiva Kazanaviciute, Elisabeth Scheikl, Markus Teige, Robert Doczi, Heribert Hirt, Manfred Schwanninger, Merijn Kant, Robert Schuurink, Felix Mauch, Antony Buchala, Francesca Cardinale, and Irute Meskiene. The pp2c-type phosphatase ap2c1, which negatively regulates mpk4 and mpk6, modulates innate immunity, jasmonic acid, and ethylene levels in arabidopsis. The Plant Cell, 19(7):2213–2224, 2007. ISSN 1040-4651. doi: 10.1105/tpc.106.049585. URL https://doi.org/10.1105/tpc.106.049585.

Alois Schweighofer, Volodymyr Shubchynskyy, Vaiva Kazanaviciute, Armin Djamei, and Irute Meskiene. Bimolecular Fluorescent Complementation (BiFC) by MAP Kinases and MAPK Phosphatases, pages 147–158. Springer New York, New York, NY, 2014. ISBN 978-1-4939-0922-3. doi: 10.1007/978-1-4939-0922-3_12. URL https://doi.org/10.1007/978-1-4939-0922-3_12.

Megan C. Shelden and Rana Munns. Crop root system plasticity for improved yields in saline soils. Frontiers in Plant Science, Volume 14 - 2023, 2023. ISSN 1664-462X. doi: 10.3389/fpls.2023.1120583. URL https://www.frontiersin.org/journals/plant-science/articles/10.3389/fpls.2023.1120583.

O. Shelef, N. Lazarovitch, B. Rewald, A. GolanGoldhirsh, and S. Rachmilevitch. Root halotropism: Salinity effects on bassia indica root. Plant Biosystems - An International Journal Dealing with all Aspects of Plant Biology, 144(2):471–478, 2010. ISSN 1126-3504. doi: 10.1080/11263501003732001. URL https://doi.org/10.1080/11263501003732001.

J. A. Siqueira, W. C. Otoni, and W. L. Araujo. The hidden half comes into the spotlight: Peeking inside the black box of root developmental phases. Plant Commun, 3(1):100246, 2022. ISSN 2590-3462 (Electronic) 2590-3462 (Linking). doi: 10.1016/j.xplc.2021.100246. URL https://www.ncbi.nlm.nih.gov/pubmed/35059627.

C. Spitzer, F. Li, R. Buono, H. Roschzttardtz, T. Chung, M. Zhang, K. W. Osteryoung, R. D. Vierstra, and M. S. Otegui. The endosomal protein charged multivesicular body protein1 regulates the autophagic turnover of plastids in arabidopsis. Plant Cell, 27(2):391–402, 2015. ISSN 1040-4651 (Print) 1040-4651. doi: 10.1105/tpc.114.135939.

S. H. Su, N. M. Gibbs, A. L. Jancewicz, and P. H. Masson. Molecular mechanisms of root gravitropism. Curr Biol, 27(17):R964–R972, 2017. ISSN 1879-0445 (Electronic) 0960-9822 (Linking). doi: 10.1016/j.cub.2017.07.015. URL https://www.ncbi.nlm.nih.gov/pubmed/28898669.

F. Sun, W. Zhang, H. Hu, B. Li, Y. Wang, Y. Zhao, K. Li, M. Liu, and X. Li. Salt modulates gravity signaling pathway to regulate growth direction of primary roots in arabidopsis. Plant Physiol, 146(1):178–88, 2008. ISSN 0032-0889 (Print) 0032-0889. doi: 10.1104/pp.107.109413.

Marci Surpin and Natasha Raikhel. Traffic jams affect plant development and signal transduction. Nature Reviews Molecular Cell Biology, 5 (2):100–109, 2004. ISSN 1471-0080. doi: 10.1038/nrm1311. URL https://doi.org/10.1038/nrm1311.

D. Szklarczyk, R. Kirsch, M. Koutrouli, K. Nastou, F. Mehryary, Hachilif, A. L. Gable, T. Fang, N. T. Doncheva, S. Pyysalo, P. Bork, L. J. Jensen, and C. von Mering. The string database in 2023: protein-protein association networks and functional enrichment analyses for any sequenced genome of interest. Nucleic Acids Res, 51(D1):D638–d646, 2023. ISSN 0305-1048 (Print) 0305-1048. doi: 10.1093/nar/gkac1000.

Tyanova, T. Temu, P. Sinitcyn, A. Carlson, M. Y. Hein, Geiger, M. Mann, and J. Cox. The perseus computational platform for comprehensive analysis of (prote)omics data. Nat Methods, 13(9):731–40, 2016. ISSN 1548-7105 (Electronic) 1548-7091 (Linking). doi: 10.1038/nmeth.3901. URL https://www.ncbi.nlm.nih.gov/pubmed/27348712 https://www.nature.com/articles/nmeth.3901.

A. A. Tyurin, K. V. Kabardaeva, M. A. Berestovoy, Yu V. Sidorchuk, A. A. Fomenkov, A. V. Nosov, and I. V. Goldenkova-Pavlova. Simple and reliable system for transient gene expression for the characteristic signal sequences and the estimation of the localization of target protein in plant cell. Russian Journal of Plant Physiology, 64(5):672– 679, 2017. ISSN 1608-3407. doi: 10.1134/S1021443717040173. URL https://doi.org/10.1134/S1021443717040173.

Thea van den Berg, Ruud A. Korver, Christa Testerink, and Kirsten H. W. J. ten Tusscher. Modeling halotropism: a key role for root tip architecture and reflux loop remodeling in redistributing auxin. Development, 143(18):3350–3362, 2016. ISSN 0950-1991. doi: 10.1242/dev.135111. URL https://doi.org/10.1242/dev.135111.

E. van Zelm, Y. Zhang, and C. Testerink. Salt tolerance mechanisms of plants. Annu Rev Plant Biol, 71:403–433, 2020. ISSN 1545-2123 (Electronic) 1543-5008 (Linking). doi: 10.1146/annurev-arplant-050718-100005. URL https://www.ncbi.nlm.nih.gov/pubmed/32167791.

A. B. Vojtek, S. M. Hollenberg, and J. A. Cooper. Mammalian ras interacts directly with the serine/threonine kinase raf. Cell, 74(1):205–14, 1993. ISSN 0092-8674 (Print) 0092-8674 (Linking). doi: 10.1016/0092-8674(93)90307-c. URL https://www.ncbi.nlm.nih.gov/pubmed/8334704.

X. Wang, M. Xu, C. Gao, Y. Zeng, Y. Cui, W. Shen, and L. Jiang. The roles of endomembrane trafficking in plant abiotic stress responses. J Integr Plant Biol, 62(1):55–69, 2020. ISSN 1672-9072. doi: 10.1111/jipb.12895.

G. West, D. Inzé, and G. T. Beemster. Cell cycle modulation in the response of the primary root of arabidopsis to salt stress. Plant Physiol, 135(2):1050–8, 2004. ISSN 0032-0889 (Print) 0032-0889. doi: 10.1104/pp.104.040022.

Honghong Wu, Lana Shabala, Elisa Azzarello, Yuqing Huang, Camilla Pandolfi, Nana Su, Qi Wu, Shengguan Cai, Nadia Bazihizina, Lu Wang, Meixue Zhou, Stefano Mancuso, Zhonghua Chen, and Sergey Shabala. Na+ extrusion from the cytosol and tissue-specific na+ sequestration in roots confer differential salt stress tolerance between durum and bread wheat. Journal of Experimental Botany, 69(16):3987–4001, 2018. ISSN 0022-0957. doi: 10.1093/jxb/ery194. URL https://doi.org/10.1093/jxb/ery194.

Lei Wu, Pan Luo, Dong-Wei Di, Li Wang, Ming Wang, Cheng-Kai Lu, Shao-Dong Wei, Li Zhang, Tian-Zi Zhang, Petra Amakorová, Miroslav Strnad, Ondřej Novák, and Guang-Qin Guo. Forward genetic screen for auxin-deficient mutants by cytokinin. Scientific Reports, 5(1):11923, 2015. ISSN 2045-2322. doi: 10.1038/srep11923. URL https://doi.org/10.1038/srep11923.

Z. Xia, Y. Huo, Y. Wei, Q. Chen, Z. Xu, and W. Zhang. The arabidopsis lyst interacting protein 5 acts in regulating abscisic acid signaling and drought response. Front Plant Sci, 7:758, 2016. ISSN 1664-462X (Print) 1664-462x. doi: 10.3389/fpls.2016.00758.

Zongliang Xia, Yangyang Wei, Kaile Sun, Jianyu Wu, Yongxia Wang, and Ke Wu. The maize aaa-type protein skd1 confers enhanced salt and drought stress tolerance in transgenic tobacco by interacting with lyst-interacting protein 5. PLOS ONE, 8(7):e69787, 2013. doi: 10.1371/journal.pone.0069787. URL https://doi.org/10.1371/journal.pone.0069787.

H. L. Xing, L. Dong, Z. P. Wang, H. Y. Zhang, C. Y. Han, B. Liu, X. C. Wang, and Q. J. Chen. A crispr/cas9 toolkit for multiplex genome editing in plants. BMC Plant Biol, 14:327, 2014. ISSN 1471-2229. doi: 10.1186/s12870-014-0327-y.

M. Xiong, J. Xu, Z. Zhou, B. Peng, Y. Shen, H. Shen, X. Xu, C. Li, L. Deng, and G. Feng. Salinity inhibits seed germination and embryo growth by reducing starch mobilization efficiency in barley. Plant Direct, 8(2):e564, 2024. ISSN 2475-4455 (Electronic) 2475-4455 (Linking). doi: 10.1002/pld3.564. URL https://www.ncbi.nlm.nih.gov/pubmed/38312996.

Hongli Yang, Jing Liu, Jiulu Lin, Linbin Deng, Shihang Fan, Yan Guo, Fengming Sun, and Wei Hua. Overexpression of chmp7 from rapeseed and arabidopsis causes dwarfism and premature senescence in arabidopsis. Journal of Plant Physiology, 204:16– 26, 2016. ISSN 0176-1617. doi: 10.1016/j.jplph.2016.06.023. URL https://www.sciencedirect.com/science/article/pii/S0176161716301493.

Y. Zeng, B. Li, S. Huang, H. Li, W. Cao, Y. Chen, G. Liu, Z. Li, C. Yang, L. Feng, J. Gao, S. W. Lo, J. Zhao, J. Shen, Y. Guo, C. Gao, Y. Dagdas, and L. Jiang. The plant unique escrt component free1 regulates autophagosome closure. Nat Commun, 14(1):1768, 2023. ISSN 2041-1723. doi: 10.1038/s41467-023-37185-6.

B. Zhang, C. Deng, S. Wang, Q. Deng, Y. Chu, Z. Bai, A. Huang, Q. Zhang, and Q. He. The rna landscape of dunaliella salina in response to short-term salt stress. Front Plant Sci, 14:1278954, 2023. ISSN 1664-462X (Print) 1664-462X (Electronic) 1664-462X (Linking). doi: 10.3389/fpls.2023.1278954. URL https://www.ncbi.nlm.nih.gov/pubmed/38111875.

